# Structural and Functional Glycosylation of the Abdala COVID-19 Vaccine

**DOI:** 10.1101/2024.09.24.614759

**Authors:** Sean A. Burnap, Valeria Calvaresi, Gleysin Cabrera, Satomy Pousa, Miladys Limonta, Yassel Ramos, Luis Javier González, David J. Harvey, Weston B. Struwe

## Abstract

Abdala is a COVID-19 vaccine produced in *Pichia pastoris* and is based on the receptor-binding domain (RBD) of the SARS-CoV-2 spike. Abdala is currently approved for use in multiple countries with clinical trials confirming its safety and efficacy in preventing severe illness and death. Although *P. pastoris* is used as an expression system for protein-based vaccines, yeast glycosylation remains largely uncharacterised across immunogens. Here, we characterise N-glycan structures and their site of attachment on Abdala and show how yeast-specific glycosylation decreases binding to the ACE2 receptor and a receptor-binding motif (RBM) targeting antibody compared to the equivalent mammalian-derived RBD. Reduced receptor and antibody binding is attributed to changes in conformational dynamics resulting from N-glycosylation. These data highlight the critical importance of glycosylation in vaccine design and demonstrate how individual glycans can influence host interactions and immune recognition via protein structural dynamics.

## INTRODUCTION

As of the end of 2023, the severe acute respiratory syndrome (SARS) coronavirus (CoV)-2 outbreak was attributed to over 7 million deaths worldwide by the World Health Organisation (WHO)^1^. The rapid development, approval and deployment of multiple COVID-19 vaccines was critical for limiting viral spread and deaths, with over 13 billion vaccine doses administered to date^1^. The major SARS-CoV-2 immunogen and neutralisation target for vaccine design is the surface “spike” glycoprotein, which recognises host cells via binding between the spike (S) receptor-binding domain (RBD) and the human angiotensin-converting enzyme-2 (ACE2)^2^. The use of full-length S trimers as immunogens required extensive protein engineering for stabilisation of pre-fusion states that resulted in altered glycosylation, protein dynamics and overall antigen structural integrity compared to spike trimers on infectious virions^3,4^.

The SARS-CoV-2 spike is a trimeric class I fusion glycoprotein comprised of an S1 outer domain, containing the RBD and N-terminal domain (NTD), and an S2 domain responsible for host membrane fusion. Spikes are heavily glycosylated, with 66 N-glycans per S trimer with each RBD domain containing 2 N-glycans that modulate protein structural states through “up” and “down” RBD dynamics^5,6^. Varying O-glycosylation has been reported, with a known functional role in modulation of furin cleavage^7-9^. Glycosylation is therefore particularly important for cellular processing and functional dynamics of COVID immunogens.

Effective worldwide vaccination relies on distributable and cost-effective production of stable vaccines. The Abdala COVID-19 vaccine, produced by the Center for Genetic Engineering and Biotechnology in Cuba ^10^, is based on a recombinantly produced RBD subunits expressed in *Pichia pastoris*, a yeast system previously used for vaccines, including SARS-CoV, hepatitis B and other SARS-CoV-2 candidates^11-15^. Clinical trials of Abdala showed it to be cost-effective, stable, safe and efficacious, fulfilling the WHO criteria for COVID-19 vaccines, and Abdala is currently authorised for emergency use in multiple countries^16-18^.

Glycoproteins produced in *P. pastoris* have different glycan structures to those found on circulating infectious viruses as well as recombinantly produced spikes or RBDs from mammalian systems, such as Chinese hamster ovary (CHO) or human embryonic kidney (HEK) cell lines^19^. In contrast to mammalian glycosylation, yeast glycans are generally high-mannose structures extending from a Man_8_GlcNAc_2_ core^20^. Therapeutic glycoproteins from *P. pastoris* remain largely uncharacterised at the glycan level but are recognised as important for vaccine immunogenicity and/or half-life, particularly through mannose-dependent clearance mechanisms, which has led to efforts for glycoengineering *P. pastoris* to produce human complex-type N-glycans^21-23^. *P. pastoris* have hyper-mannosylated glycans, however the extent of mannosylation, degree of N-glycan phosphorylation and precise structural information of the individual glycans and their protein site of attachment are currently undefined for Abdala. Equally, the effect of yeast glycosylation and comparisons with mammalian-derived RBD in antibody and receptor binding, which is critical for immunogenicity and overall vaccine efficacy, are unknown.

Here, we combine mass spectrometry (MS) methods in glycomics, glycoproteomics and hydrogen-deuterium exchange (HDX) to provide a comprehensive structural and dynamical understanding of the Abdala RBD vaccine. Using mass photometry (MP), a label-free single molecule imaging method ^3,24^, we quantify binding to ACE2 and an RBD-targeting antibody. Comparison with HEK cell derived RBD, we show how differences in site-specific glycosylation affect antibody and receptor binding that arise from conformational dynamics of the receptor binding motif (RBM) within the RBD glycoprotein. Taken together, these data represent the first full site-specific glycan characterisation of a structure-based vaccine from *P. pastoris* that uncovers how subtle changes in RBD glycosylation can shape immunogen functionality.

## RESULTS

### Batch-to-batch characterisation of Abdala N-glycosylation

SDS-PAGE of Abdala and HEK-derived RBD (RBD_HEK_), showed Abdala to exhibit a broader mass distribution that migrated at a higher molecular weight, between 40 and 70 kDa, compared to RBD_HEK_ which migrated at approximately 37 kDa (**Fig. 1a**). The theoretical masses are 25.9 kDa for RBD_HEK_ and 26.1 kDa for Abdala, and following PNGaseF treatment both proteins had similar migration patterns, confirming differences in observed mass are from N-glycosylation. Additionally, the measured mass by MP confirmed a difference of approximately 11 kDa between RBD_HEK_ (37 kDa) and three separate production batches of Abdala (48 ± 1 kDa) (**Fig. 1b, Supplementary Fig. 1**). MP also showed a minor population of dimeric species for both RBD_HEK_ and Abdala, at approximately 19.6 and 5.6 %, respectively. Dimer formation was consistently lower across three batches of Abdala compared to RBD_HEK_ (**Supplementary Fig. 1**) but was not detected by size exclusion chromatography. Our results were consistent with a previous report that detected dimerization of HEK derived RBD using native MS^25^.

**Fig. 1:**
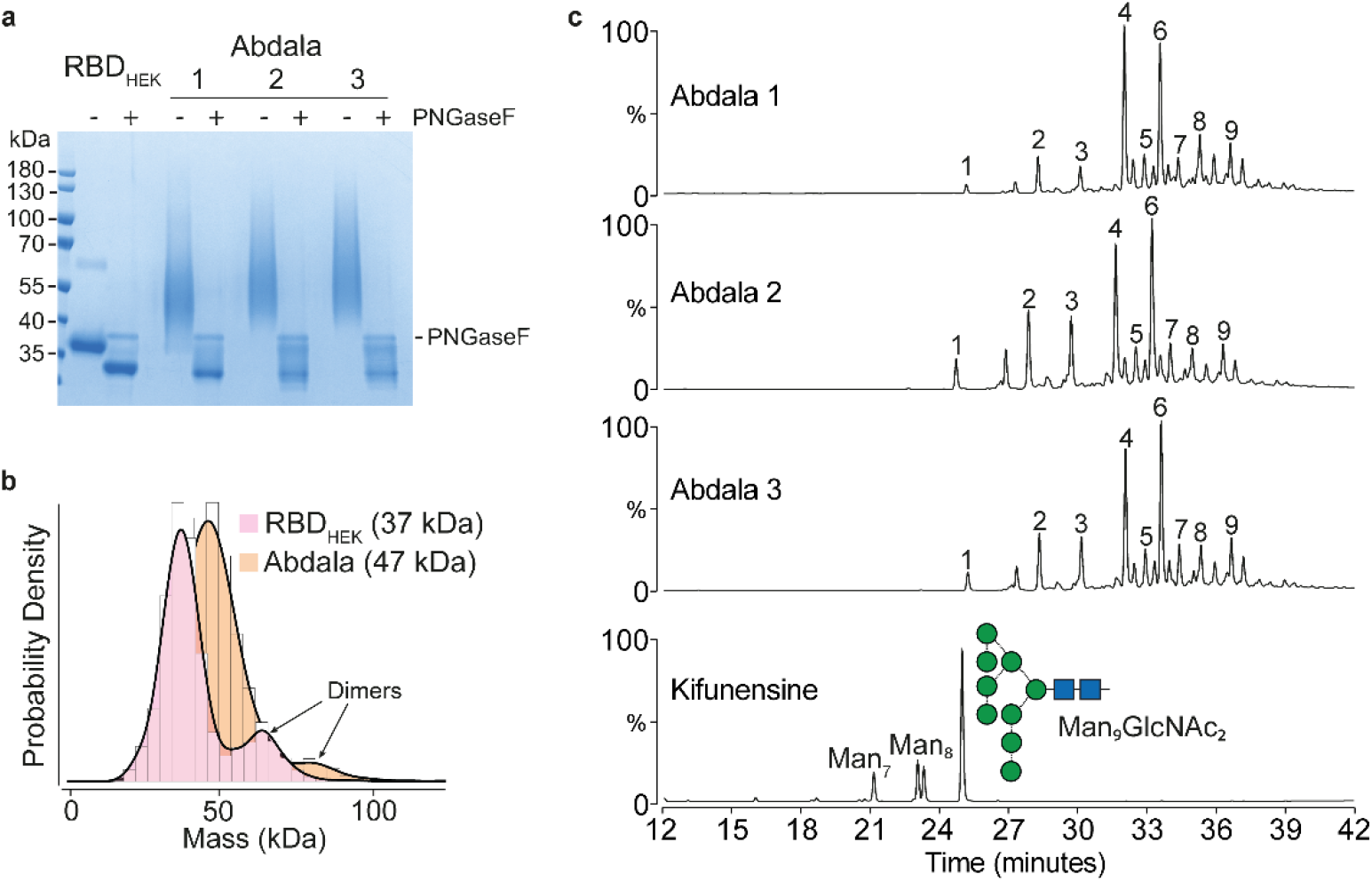
HPLC-based analysis of Abdala N-glycans. (**a**) SDS-PAGE of RBD_HEK_ and three separate production batches of Abdala +/-PNGaseF digestion. (**b**) Mass photometry of RBD_HEK_ and Abdala. (**c**) HPLC of N-glycans from three production batches of Abdala and a kifunensine-derived RBD control.

To quantify N-glycosylation across Abdala batches, we analysed fluorescently labelled N-glycans via high performance liquid chromatography (HPLC) (**Fig. 1c**)^3^. As a control, we included a glycoengineered form of RBD that was recombinantly expressed in the presence of kifunensine to contain predominantly Man_9_GlcNAc_2_ *N*-glycans, which assists preliminary assignments of Abdala. Expectedly, Abdala profiles were distinct from kifunensine-derived RBD as N-glycans eluted later than the major Man_9_GlcNAc_2_ peak, indicative of hyper-mannosylated structures. RBD_HEK_ N-glycan profiles contained 11 major N-glycan peaks all eluting earlier than Man_9_GlcNAc_2_ (<26 minutes), suggestive of complex-type N-glycans (**Supplementary Fig. 2a**). Batch-to-batch variation was minimal across the 9 major N-glycan HPLC peaks, with a mean coefficient of variation equal to 22%. The relative abundances of peaks 4 and 6 varied most across the three batches (**Fig. 1c, Supplementary Fig. 2b, Supplementary Table. 1**).

### *N*-glycan structural analysis by ion mobility-MS/MS

Ion mobility (IM)-MS/MS of Abdala *N*-glycans revealed detailed information of their hyper-mannosylated structures as well as the extent of phosphorylation (**Fig. 2a**). N-glycan ions were predominantly doubly charged and were detected as deprotonated [M-2H]^2-^ species. The mass difference between each peak was ∼81 *m/z*, corresponding to a hexose (presumably mannose) residue (162 Da). The overall monosaccharide compositions correspond to Man_9-16_GlcNAc_2_Phos_2_ and their proposed structures are schematically represented in the spectrum. The MS spectra of the three Abdala batches were consistent and it underscored batch-to-batch production consistency with near identical glycan peak distribution and abundances (**Supplementary Fig. 3**). One notable difference was that Abdala batch 1 had a greater Man_10_/Man_11_ ratio compared to batches 2 and 3, which indicates HPLC peaks 4 and 6 are likely Man_10_GlcNAc_2_Phos_2_ and Man_11_GlcNAc_2_Phos_2_, respectively.

**Fig. 2:**
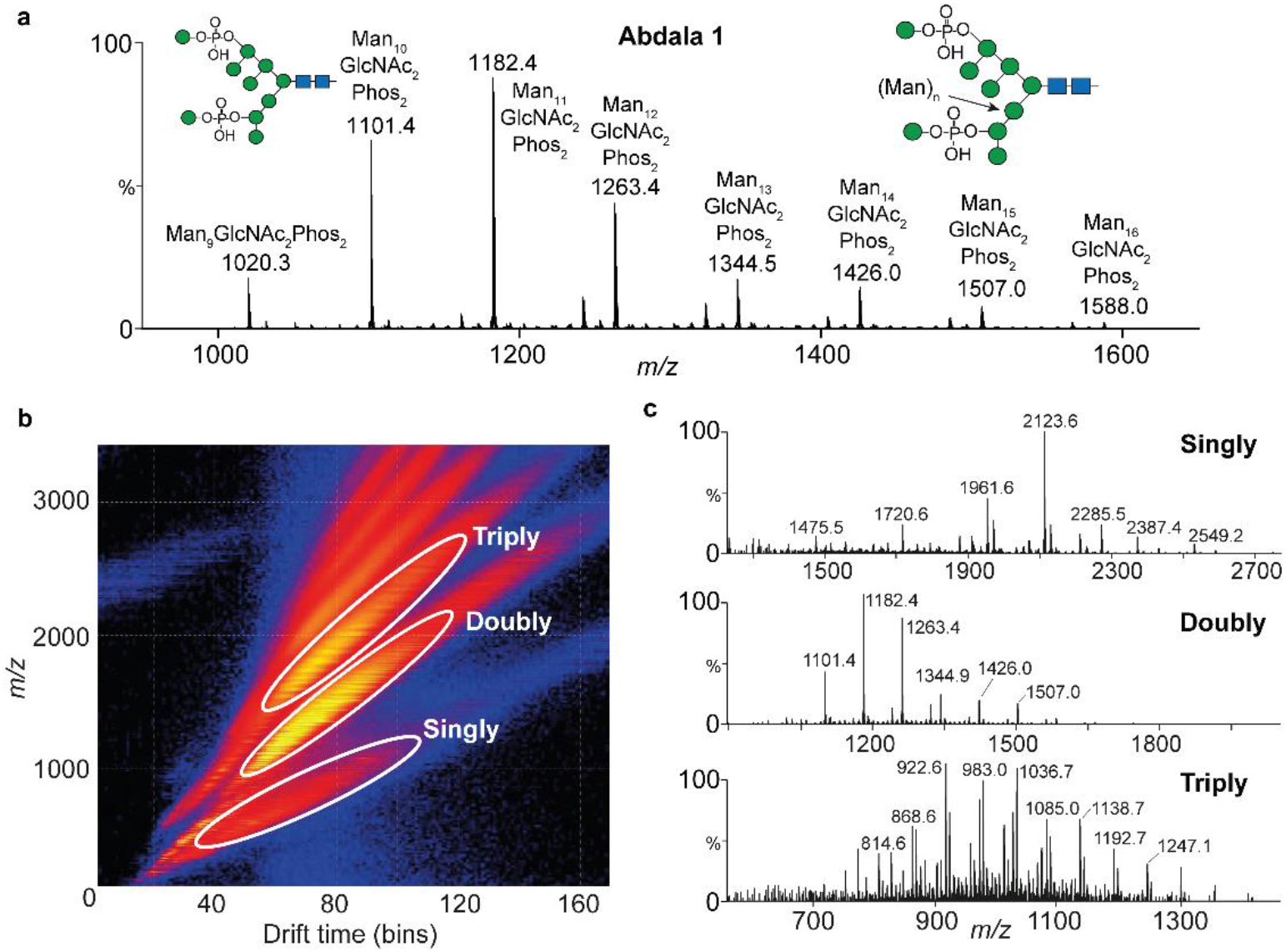
IM-MS of Abdala N-glycans. (**a**) MS spectrum of Abdala N-glycans (batch 1). (**b**) Representative drift plot of Abdala N-glycans showing the distribution of singly, doubly and triply charged ions. (**c**) Corresponding extracted MS spectra of singly, doubly and triply charged N-glycans. Glycan compositions and ion assignments are annotated in Supplementary Tables 2-5.

IM separation (**Fig. 2b**) and extraction of singly, doubly and triply charged ions revealed additional structural information, specifically the presence of glycans with 1-3 phosphate groups and up to 22 mannose residues. Singly charged ions had a major series of Man_5-14_GlcNAc_2_Phos_1_ detected as [M – H]^-^ ions and a minor series of Man_9-13_GlcNAc_2_Phos_2_ as [M – H + Na]^-^ (**Fig. 2c** (top spectrum), **Supplementary Table. 2**). The major N-glycan ion was 2123.6, which is Man_10_GlcNAc_2_Phos_1_, followed by Man_9_GlcNAc_2_Phos_1_ (1961.6 *m/z*) and Man_11_GlcNAc_2_Phos_1_ (2285.5 *m/z*). IM extraction of doubly charged ions uncovered additional mannose residues beyond those shown in Fig. 2a, specifically N-glycans with 2 phosphates and up to 21 mannose residues (Man_9-21_GlcNAc_2_Phos_2_). Ions were detected as [M – 2H]^2-^ and/or [M – H + Na + H_2_PO_4_]^2-^ species (**Fig. 2c** (middle spectrum), **Supplementary Table. 3**). The spectrum of doubly charged ions is similar to the MS spectra of all N-glycans shown in **Fig. 2a**, as Man_9-16_GlcNAc_2_Phos_2_ are the major N-glycan structures present on Abdala. Triply charged ions were predominantly Man_10-22_GlcNAc_2_Phos_3_ N-glycans that were detected as [M – 3H]^3-^, [M – 2H + H_2_PO_4_]^3-^ and [M – 2H + Na + H_2_PO_4_]^3-^ ions (**Fig. 2c** bottom spectrum, **Supplementary Table. 4**). A minor amount of N-glycans with two phosphates were also detected as Man_11-19_GlcNAc_2_Phos_2_. Overall, Abdala glycans were Man_5-22_GlcNAc_2_, with the dominant species being di-phosphorylated species (**Supplementary Table. 5**).

MS-based fragmentation revealed structural information of the underlying glycan (representative MS/MS spectra from Abdala batch 1 are shown). Collision induced dissociation of all N-glycans resulted primarily in the neutral loss of mannose residues with minimal diagnostic A-type cross-ring fragments (**Fig. 3**). One cross-ring fragment was found, specifically ^0,2^A_R_ of the reducing end GlcNAc. Fragmentation across glycan species revealed similar spectra, with all ions increasing by the mass of mannose residues, suggestive of the core structure of all compounds to be the same (Man_10-14_GlcNAc_2_Phos_2_).

**Fig. 3:**
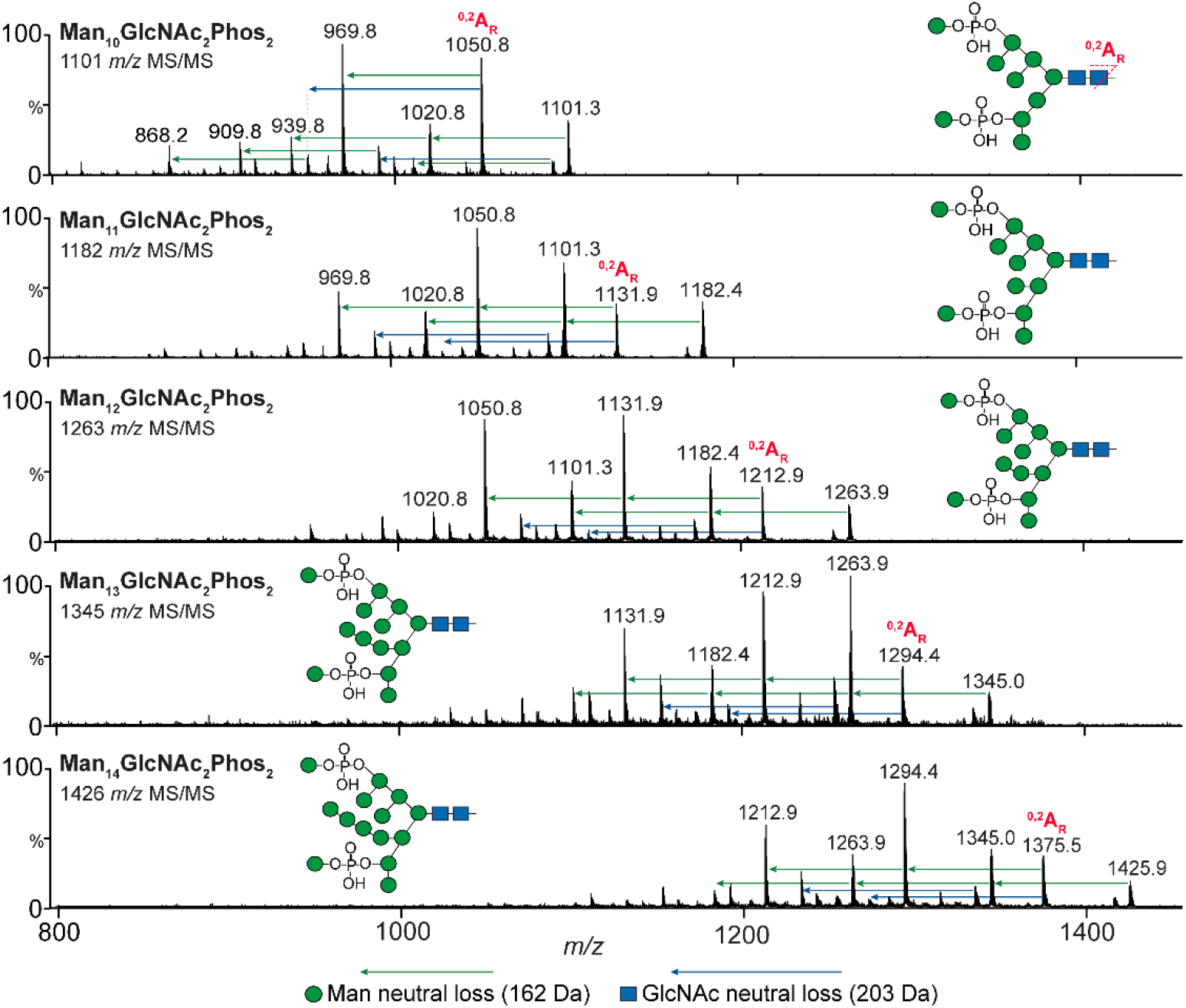
Fragmentation spectra of Abdala N-glycans. MS/MS fragmentation spectra of Man_10-14_GlcNAc_2_Phos_2_ N-glycans showing characteristic loss of mannose (green arrows) and N-acetylglucosamine (GlcNAc) residues (blue arrows). Proposed N-glycan structures are shown. Precursor ions are [M – 2H]^2-^ species.

IM-MS analysis was also conducted on RBD_HEK_ N-glycans. The total acquired MS spectra (**Supplementary Fig. 4a**), as well as mobility extracted doubly (**Supplementary Fig. 4b**) and triply (**Supplementary Fig. 4c**) charged ions were primary doubly charged complex type N-glycans, specifically core-fucosylated, bi and tri-antennary structures with terminal sialyation, with Man_5_GlcNAc_4_Fuc_1_NeuAc_2_ as the most abundant glycan (**Supplementary Fig. 4b**). IM extraction of triply charged ions revealed a minor presence of tetra-antennary, complex glycans with varying levels of terminal sialyation (**Supplementary Fig. 4c**). While multiple studies have performed glycan analyses on recombinant full-length spikes or isolated S1 domains, glycoproteomics has been the method of choice to characterise RBD N-glycosylation^26,27^. Our IM-MS analysis of RBD_HEK_ aligns with previous bottom-up glycoproteomic studies that show RBD N-glycans as primarily bi-or tri-antennary with varying degrees of sialyation. Glycan analysis via IM-MS revealed a greater degree of sialyation than previously shown, potentially due to a reduced loss of labile sialic acids during ionisation of glycopeptides.

### Site-specific N- and O-glycan analysis

Having characterised Abdala and RBD_HEK_ *N*-glycans, we next identified their sites of attachment via bottom-up glycoproteomics. It is important to note that both Abdala and RBD_HEK_ contain amino acids 319-541 present on the native Wuhan-Hu-1 SARS-CoV-2 virus, but the Abdala immunogen contains additional sequences at both termini (**Fig. 4a**). There are two N-glycan sites in this region, N331 and N343 which are distal to the RBM region responsible for ACE2 binding. The additional Abdala N*-*terminal sequence is NWSFFSNIGGSSGGS, which is flexible and polar, and added to prevent protein aggregation ^10^. This introduces an additional N-glycan sequon (N-X-S/T, X≠P) at the first residue, **<u>N</u>**WS. The Abdala C-terminal sequence is GGSGGSSSSSSSSSSIEHHHHHH and was added to improve His-tag protein purification^10^. Both Abdala and RBD_HEK_ have the endogenous N331 and N343 N-glycan glycosylation sites. Furthermore, the spike RBD is known to have varying levels of O-glycosylation, with a report showing up to 8 glycoforms at T323 with a core 1, di-sialyated O-glycan as the most abundant structure^28^. *P. pastoris* has a distinct type of protein O-glycosylation in the form of O-mannosylation, where up to 6 mannose residues can be added primarily to S/T residues, primarily in α1,2 linkages^29^. Importantly, O-mannosylation is not present on human-derived glycoproteins, and has been shown to enhance antigen immunogenicity specifically through CD4^+^ T cell responses, highlighting the importance of characterising the extent of O-glycosylation in addition to N-glycosylation on Abdala^30^.

**Fig. 4:**
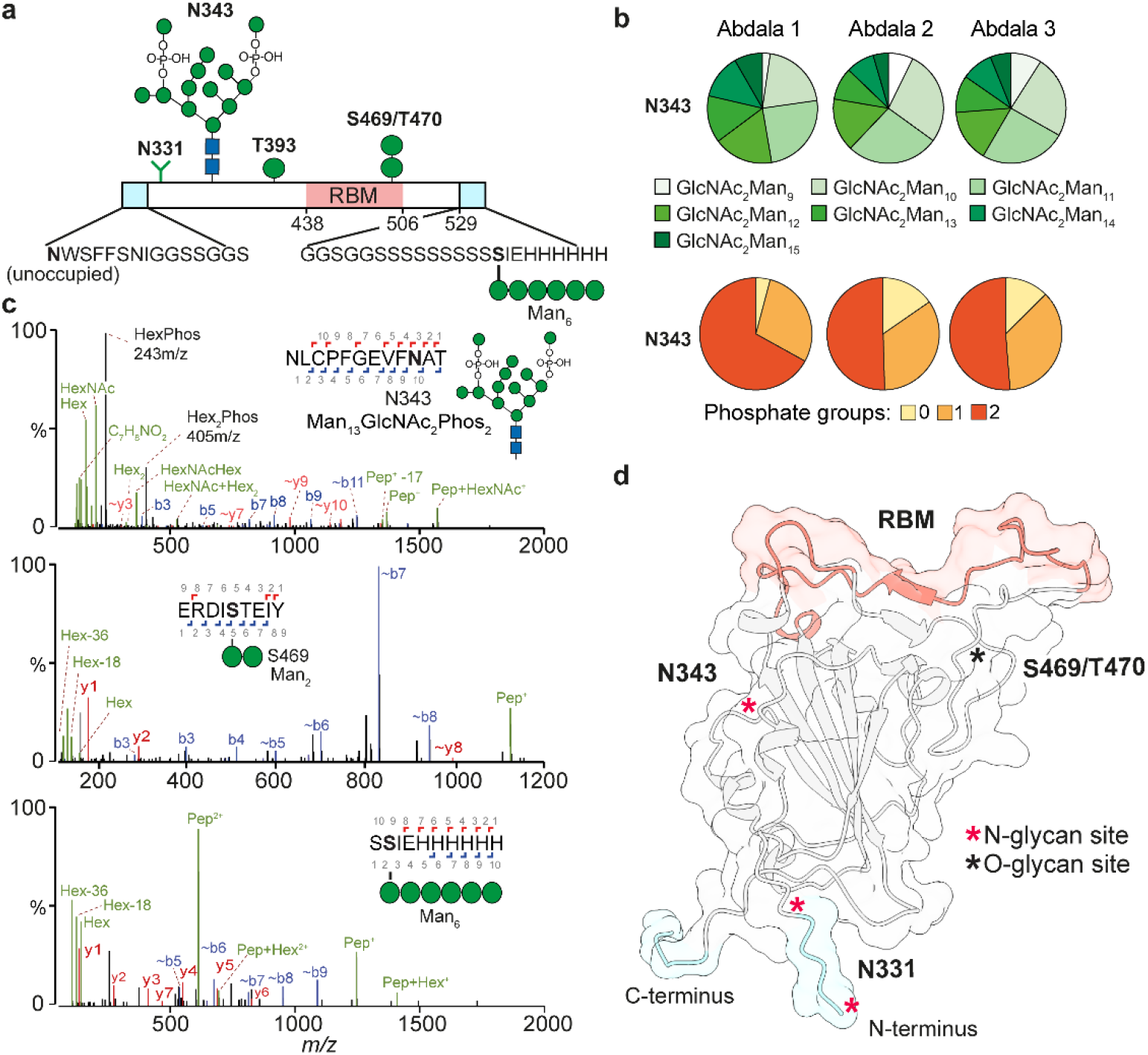
Glycoproteomic characterisation of Abdala. (**a**) Schematic representation of Abdala amino acid sequence, highlighting N- and C-terminal regions, RBM and the major N- and O-glycans identified from this study. (**b**) Pie charts reflecting site-specific glycosylation analysis by glycoproteomics. Glycans are characterised by the number of mannose residues and phosphate groups. **c** MS/MS spectra of a glycopeptide containing Man_13_GlcNAc_2_Phos_2_ at N343 (top). O-mannosylation at position S469 (middle) and O-mannosylation in the C-terminal extension (bottom). **d** AlphaFold model of Abdala showing the location of N-glycans, RBM and Abdala N- and C-termini.

Consistent with IM-MS data of released *N*-glycans, glycoproteomics of the three Abdala batches identified Man_9-15_GlcNAc_2_ with between 0-2 phosphate groups as the major species at N343 (**Fig. 4b**, green pie charts). The majority glycans contained two phosphates (56%) followed by one (33%) and zero (11%) (**Fig. 4b**, orange pie charts). Consistent with the HPLC data, glycoproteomics identified more similarity between batches 2 and 3 than batch 1. Opposed to glycan MS, the bottom-up analysis did not detect glycopeptides containing three phosphate groups, which is most likely due to their low abundance and poor detection efficiency. The presence of phosphorylated glycans was evidenced by 243 *m/z* (HexPhos) and 405 *m/z* (Hex_2_Phos) oxonium ion fragments in the MS/MS spectra (**Fig. 4c**, top spectrum). Chymotrypsin, alpha-lytic or ProAlanase proteases were predicted to generate theoretical peptides spanning N331 with Abdala, but we did not identify any glycopeptides spanning this site, nor did we detect the equivalent non-glycosylated peptide(s). The final N-glycosylation site at the N-terminal sequon (**<u>N</u>**WS) was only detected as unoccupied (**Supplementary Fig. 5a**). On the other hand, analysis of RBD_HEK_ identified complex-type glycans at positions N331 and N343, consisting of bi-, tri- and tetra-antennary structures (**Supplementary Fig. 6a**). While bi- and tri-antennary glycans were also most abundant in the IM-MS analysis, a lesser degree of sialyation was observed by glycoproteomics with the most abundant glycoform harbouring a single sialic acid (**Supplementary Fig. 6b**). We did not detect any non-glycosylated peptides spanning both N331 and N343 sites and contrary to a previous report we only identified one *O*-glycan at position T430 on RBD_HEK_ (**Supplementary Fig. 6b**)^25^.

Multiple O-mannosylated peptides were observed on Abdala and their presence was evidenced by hexose oxonium ion fragments (162 m/z, with and without the loss of H_2_O) within the MS/MS spectra. Two mannose residues were identified at S469/T470 (**Fig. 4c**, middle spectrum), while one mannose residue was identified at position T393 (**Supplementary Fig. 5b**). A longer Man_6_ glycan was detected within the C-terminal region of Abdala (S**S**IEHHHHHH), highlighting a previously unknown site arising from the addition of a repeating serine sequence (**Fig. 4c**, bottom spectrum). Importantly, O-mannosylated peptides were also observed in a non-glycosylated state with varying degrees of glycan occupancy. To understand the extent of glycan occupancy, the abundances of non-glycosylated peptides were summed and made relative to their glycosylated counterparts. The O-glycan sites had different degrees of occupancy: T393 was <1% occupied, S469/T470 were approximately 2.1% occupied while the Man_6_ glycan at S544 within the C-terminal region S**S**IEHHHHHH had an occupancy of 12.7%. Overall, O-mannosylation is relatively low on Abdala.

### Quantifying receptor and antibody binding by mass photometry

Glycomics and glycoproteomics show clear differences between Abdala and RBD_HEK_, including the presence of O-mannosylation. Although RBD N-glycans are distal to the ACE2 binding interface, molecular dynamics simulations have shed light on the role N343 plays in maintaining RBD structural integrity via shielding of the hydrophobic RBD core^6^. Secondly, it is thought that N343 can influence opening and closing of the RBD in the context of pre-fusion spike trimers^5^. The extent of glycan-driven structural change in isolated RBD as a result of the presence of hyper-mannosylated N-glycans is unknown. Quantifying ACE2 binding would inform whether the structural integrity (i.e. protein folding) of Abdala is maintained and comparable to RBD_HEK_.

Elicitation of high-affinity, neutralising antibodies in response to Abdala is critical for vaccine efficacy. The primary mode of SARS-CoV-2 neutralisation is through direct interaction between antibodies and the RBM, through similar contact residues with ACE2 binding^31,32^. Class 3/4 neutralising antibodies however, defined through binding distal to the RBM, have distinct neutralisation mechanisms with mAb S309 as a notable example that makes direct contact with the N343 glycan^31,32^. Due to Abdala glycans being greatly different to those derived from mammalian culture, and glycans found on the spike of live virus, antibodies targeting Abdala glycan epitopes are unlikely to be effective in combatting SARS-CoV-2 infection. A quantitative understanding of binding between Abdala and RBD_HEK_ is important for understanding possible differences in functionality between glycovariants, in particular testing if phosphorylated oligomannose N-glycans or O-mannosylation alters receptor or antibody binding.

Here, we us MP to quantify binding affinities (K_d_ values) via single molecule counting, as we have done previously^3,24,33^. MP has the advantage over conventional biophysical methods in its ability to quantify each K_d_ in a multivalent interaction, which are characteristic of IgGs. To ensure accurate quantification, all proteins were first measured individually to confirm the concentration (particle counts), expected mass distribution (monomer vs. oligomers) and purity (**Supplementary Figs. 1 and 7a**). Prior to each measurement, proteins were mixed at a 1:1 molar ratio and measured after 5 minutes incubation at room temperature. We first calculated the dissociation constant between Abdala and a monomeric form of ACE2 (mACE2). Abdala bound mACE2 with a K_d_ = 55.5 ± 6.1 nM, which was 60% weaker than RBD_HEK_ (K_d_ = 22 ± 1.8 nM (**Fig. 5a, Supplementary Figs. 7b and 6c**). The interaction between RBD and mACE2 can also be expressed in terms of occupancy whereby the sum of all mACE2 counts, including species with 0 and 1 RBD molecule bound, is compared to the total counts of RBD-bound species. RBD_HEK_ exhibited a binding occupancy of 33% compared to Abdala at 24% (**Fig. 5a, Supplementary Fig. 7c**).

**Fig. 5:**
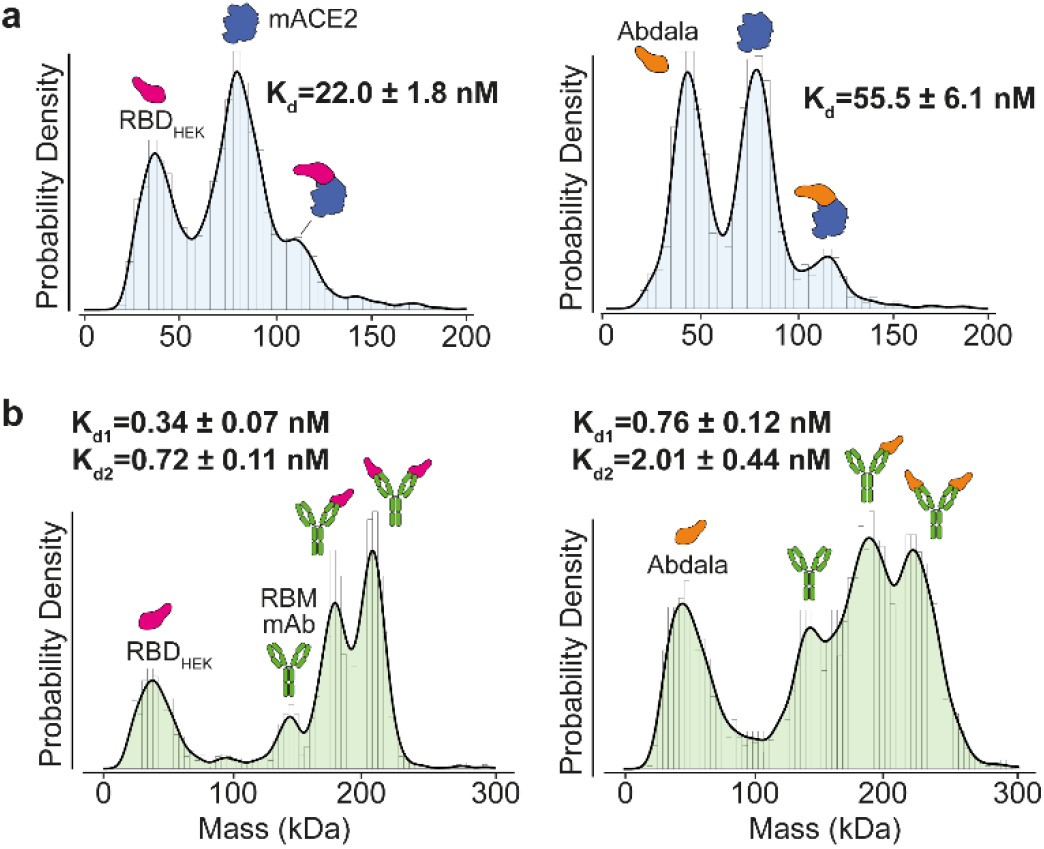
Quantifying Abdala interactions with mass photometry. (**a**) MP RBD_HEK_ (pink) and Abdala (orange) with monomeric ACE2 (mACE2) at a 1:1 molar ratio. (**b**) MP of RBD_HEK_ and Abdala with a receptor-binding motif targeting antibody at a 1:1 molar ratio.

To determine whether K_d_ differences were consistent in the context of antibody binding, we tested an RBM targeting monoclonal antibody, which is currently under investigation (**Supplementary Fig. 7d**). Each IgG bound two RBDs when mixed and measured at an equimolar ratio following equilibration at room temperature. In line with the ACE2 results, the antibody bound Abdala with weaker affinities compared to RBD_HEK_, with an average percent occupancy of bound mAb compared to free mAb equal to 82% (Abdala) compared to 89% (**Supplementary Figs. 7e and f**). For RBD_HEK_, the first K_d1_ was 0.34 ± 0.07 nM (mAb_1_:RBD_1_) with the second K_d2_ = 0.72 ± 0.11 nM (mAb_1_:RBD_2_). Abdala had a K_d1_ = 0.76 ± 0.12 nM (mAb_1_:RBD_1_), which is 55% weaker compared to mAb_1_:RBD_1HEK_. The second interaction (mAb_1_:RBD_2_) was an order of magnitude weaker (K_d2_ = 2.01 ± 0.44 nM) but still in the low nM range (**Fig. 5b**).

### Probing RBD dynamics by HDX-MS

We compared dynamics of Abdala and RBD_HEK_ by following the HDX of approximately 80 peptides spanning 83% of the protein sequences with a redundancy of 5.8, including the O-mannosylated peptides (S469/T470 and S543/544) (**Supplementary Fig. 8**). We tracked HDX at 4°C, 23°C and 28°C and over 5 time points (10s, 100s, 1000s, 10000s and 9 h) (**Supplementary Table 6, 7**). We observed an increase in HDX with Abdala that arises from an increase in flexibility, dynamics and solvent accessibility, specifically regions spanning residues 349-361, 407-415 and 454-471. Conversely, a decrease in HDX was observed between residues 472-487 indicating conformational restriction and reduced accessibility compared to RBD_HEK_ (**Supplementary Figs. 9 and 10**). Mapping the effects on the structure of the RBD-ACE2 complex (pdb:6M0J), reveals how Abdala N-glycans induce a destabilization of the adjacent protein segment (+20% of relative fractional uptake (RFU)), and the long-range allosteric propagation of destabilizing effects results in the rigidification of the RBM segment that directly engages ACE2 (−25% RFU) (**Fig. 6a**). Notably, decreased HDX in the RBM was observed across multiple time intervals. Overall, the reduced HDX in the Abdala RBM supports the observed reduction in binding to ACE2 and an RBM-targeting mAb compared to RBD_HEK_ by MP.

**Fig. 6:**
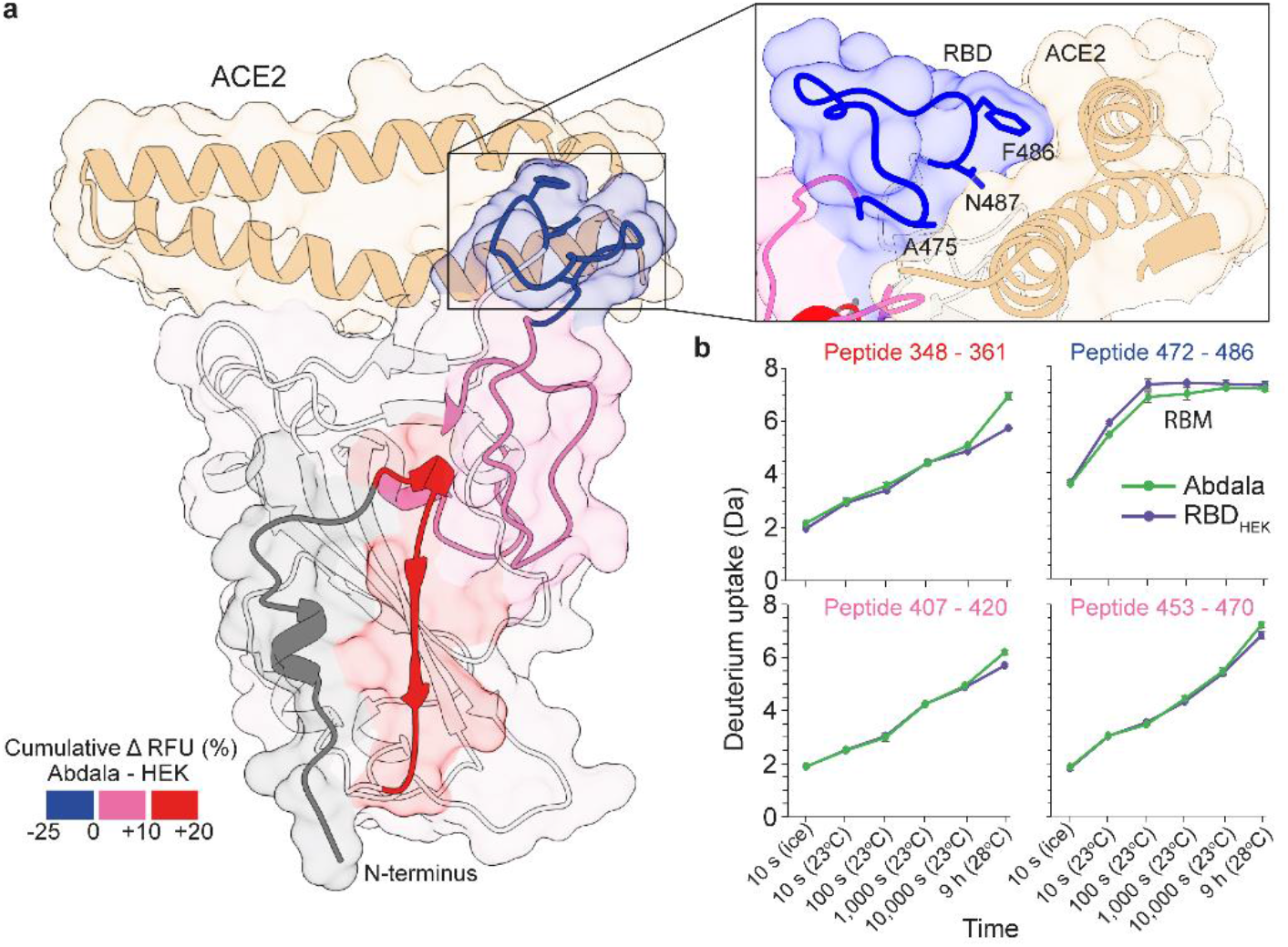
Differences in conformational dynamics probed by HDX-MS. (**a**) Protein regions with differences in HDX superimposed onto the crystal structure of RBD with ACE2 (pdb:6M0J). Red and pink indicate areas with increased dynamics (HDX) and blue indicates reduced dynamics (HDX) of Abdala compared to RBD_HEK_. N-glycan positions are indicated by asterisks. RBD-ACE2 interface with RBD contact residues (inset). (**b**) Deuterium uptake plots of peptides spanning regions with differences in HDX (colour coded as in panel **a**).

## DISCUSSION

Here, we characterise the glycosylation of the COVID-19 vaccine Abdala which uncovers distinct structures, namely phosphorylated hyper-mannosylated N-glycans and the presence of O-mannosylation. Despite the distance of these glycan sites to the RBM, we quantify how differences in glycosylation have a tractable effect in ACE2 binding as well as a neutralising IgG antibody as compared to the mammalian-derived equivalent RBD. Variations in binding affinities can be attributed to changes in protein conformational dynamics, which we believe to be caused by *P. pastoris* N- and O-glycosylation. While *P. pastoris* derived glycans are vastly different to the glycan structures observed on mammalian-derived proteins, and therefore glycans found on virions, Abdala has been shown to be efficacious in reducing severe illness and death caused by SARS-CoV-2.

Although Abdala has hyper-mannosylated, primarily phosphorylated Man_9-16_GlcNAc_2_ structures, it remains a viable immunogen. To this point, separate parallel efforts by the Argentinian AntiCovid Consortium are ongoing that also uses the RBD subunit expressed in *P. pastoris* as a vaccine candidate. It was recently demonstrated that an enzymatically deglycosylated RBD from *P. pastoris* elicited a 10-fold greater anti-RBD IgG titre in a preclinical mouse model, which plausibly results from altered protein half-life^38^. The effect of yeast glycans on protein half-life are well known, primarily driven by the action of carbohydrate receptors, specifically mannose-binding, promoting glycoprotein clearance^23,39,40^. In our hands, both PNGaseF and EndoH treatment of the Abdala vaccine led to evident protein precipitation (data not shown), highlighting the expected role that glycans play in protein solubility^41^. Abdala is stable up to 25-37°C as demonstrated in an accelerated thermal stress study conducted at five temperatures for 15 days^42^. Protein stability was evaluated by circular dichroism spectroscopy analysis, mass spectrometry (including disulfide bond assignment), functionality by immunogenicity assays and inhibiting RBD-ACE2 binding^41^. It is reasonable that glycosylation contributes to these results.

The observation that Abdala binds ACE2 and an IgG with lower affinities was not expected based on the view that N- and O-glycans are not close to the RBM, a region that shares 100% sequence homology between Abdala and RBD_HEK_. Although the measure K_d_ values with MP are all in the nM range, Abdala binding to mACE2 and IgG was approximately twice as weak compared to RBD_HEK_. We cannot rule out these differences arise from the additional C- and N-termini, in addition to changes in N-/O-glycosylation. We postulated changes in binding at the RBM is driven by altered protein structural dynamics within the core RBD. This hypothesis was influenced by molecular dynamics simulations showing the RBD N343 glycan, which is amphipathic in nature when mammalian-derived, is integral to preserving structural integrity of the RBD hydrophobic core. Loss of this glycan was shown to consistently trigger conformational change^6^. It could therefore be envisaged that replacement of a complex, amphipathic glycan on RBD_HEK_, with a larger phosphorylated hyper-mannosylated structure would disrupt the structural integrity of the RBD core in a similar manner.

Our HDX-MS results indeed uncovered regions of Abdala with greater deuterium uptake in regions proximal to the beta-sheet core, suggestive of a destabilizing effect, that propagates to the RBM, explaining differences in ACE2 and IgG binding affinities. The impact of changes in N-glycosylation on protein structure and dynamics, as tracked by HDX-MS, has also been exemplified in other proteins, highlighting N-glycosylation to regulate long-range protein conformational dynamics, impacting ligand binding and enzymatic activity^43,44^. Alterations in protein dynamics in regions distal to N-glycans within the RBD N-terminus suggests allosteric effects from *P. pastoris N*-glycosylation that are transmitted to regions directly engaging with ACE2 and the RBM-targeting mAb investigated in our study.

In conclusion, Abdala glycosylation modulates protein structural dynamics and binding to RBM targeting proteins. While N-glycan differences modulate Abdala protein function in the context of receptor and antibody binding, neutralising epitopes are preserved as evidenced by clinical trials validating Abdala to be efficacious in eliciting cross-neutralising antibody responses and reducing severe illness and death caused by SARS-CoV-2.

## MATERIALS AND METHODS

### Protein Expression

Vector pCAGGS containing the SARS-CoV-2, Wuhan-Hu-1 Spike Glycoprotein receptor binding domain (RBD) was a kind gift from the Krammer Laboratory, Department of Microbiology, Icahn School of Medicine at Mount Sinai, New York. pHL-sec vector encoding *C*-terminally His-tagged monomeric (a.a 19-611) ACE2 was a kind gift from the Zitzmann Laboratory, Department of Biochemistry, University of Oxford. The production of Abdala RBD in *P. pastoris* has been described extensively elsewhere^10^. Three production lots of Abdala RBD were assessed in this manuscript, one lot from a 75 L production scale and two lots from a 3000 L production scale. RBD and monomeric ACE2 were transiently expressed in HEK293F (FreeStyle™, Thermo Fisher Scientific). Cells were cultured in Freestyle 293 expression media (ThermoFisher Scientific) and incubated at 37°C, 8% CO2 and 120 rpm. Transfection was achieved using FreeStyle™ MAX reagent (Invitrogen) and OptiMEM™ (Gibco) following a published protocol ^45^. For the kifunensine control RBD protein, kifunensine was added at time of transfection at a final concentration 10 µM. Five days post transfection, cell culture supernatant was harvested by centrifugation at 3000 x g for 10 min and then filtered using 0.45 µM pore size filters (Merck). The supernatant was supplemented with 10 mM imidazole prior to purification using a HisTrap HP, 5mL column (Cytiva) connected to an ÄKTA pure protein purification system (Cytiva). Both RBD and monomeric ACE2 were further purified by size exclusion chromatography (SEC) using a Superose 6 increase 10/300 GL column (GE Healthcare) equilibrated in Dulbecco’s phosphate-buffered saline (DPBS, pH 7.4, ThermoFisher Scientific). SEC fractions were pooled and concentrated using Amicon molecular weight cut-off centrifugal filters (GE Healthcare). Protein concentrations were determined using a Nanodrop spectrophotometer (Thermo Fisher Scientific) at absorbance 280 nM and corrected for protein molecular weight and extinction coefficient.

### Glycan HPLC

Approximately 10 µg of RBD_HEK_ and Abdala was loaded onto SDS-PAGE gels, run at 200V for 30 min and Coomassie stained. Gel bands were excised and de-stained in 50:50 MeCN: water. PNGase F (generated in-house^46^) was added to each gel-band and incubated for 16 hours at 37°C. Released N-glycans were labelled with 2-aminoanthranilic acid (2-AA) as previously described^47^. Briefly, glycans were resuspended in 30 μL of HPLC-grade H_2_O followed by addition of 80 μL of labelling mixture (30 mg/mL 2-AA and 45 mg/mL sodium cyanoborohydride in a solution of sodium acetate trihydrate [4% w/v] and boric acid [2% w/v] in methanol). N-glycans were incubated at 80°C for 1 hour. Excess label was removed using Spe-ed Amide-2 cartridges (Applied Separation) as described ^47^.

Fluorescently labelled N-glycans were profiled by hydrophilic interaction liquid chromatography-high-performance liquid chromatography (HILIC-HPLC) using a 2.1 mm × 10 mm Acquity BEH Amide Column (1.7 μm particle size) (Waters, Elstree, UK) attached to an Agilent 1260 Infinity II (Agilent, Manchester, UK). Mobile phase was solvent A: 50 mM ammonium formate, pH 4.4 and solvent B: MeCN. The gradient was: (t = 0): 22.0% A, 78.0% B (flow rate of 0.5 mL/min); t = 38.5: 44.1% A, 55.9% B (0.5 mL/min); t = 39.5: 100% A, 0% B (0.25 mL/min); t = 44.5: 100% A, 0% B (0.25 mL/min); t = 46.5: 22.0% A, 78.0% B (0.5 ml/min), t = 48: 22.0% A, 78.0% B (0.5 mL/min). Fluorescence was measured using an excitation wavelength of 360 nm and a detection wavelength of 425 nm.

### Ion mobility – MS/MS

IM-MS/MS measurements were performed on a Synapt G2 instrument (Waters, Manchester, UK). For each analysis, 2 μL of N-glycan sample material was ionized by nano-electrospray ionization (nano-ESI) from gold-coated borosilicate glass capillaries prepared in-house ^48^. The instrument was set in sensitivity mode with the following: capillary voltage 0.8–1.2 kV, sample cone 65 V, extraction cone 3.3 V, source temperature 80°C. Collision-induced dissociation was performed both before and after mobility separation in the trap and transfer cells respectively with argon as the collision gas. The instrument was externally mass-calibrated with sodium iodide and the mobility cell was calibrated with dextran ((Glc_2–13_) (from Leuconostoc mesenteroides). Data acquisition and processing were carried out using Waters Driftscope (version 2.8) software and MassLynx™ (version 4.1). The scheme devised by Domon and Costello^49^ was used to name the fragment ions with the following exception: the subscript R (for reducing terminal) is used when general reference is made for loss or fragmentation of a GlcNAc residue from the reducing terminus of the glycan in order to avoid confusion caused by the subscript number changing as the result of altered chain lengths. Interpretation of the spectra followed rules developed earlier in this laboratory^50-53^.

### Glycoproteomics

Approximately 5 µg protein was loaded and run on an SDS-PAGE. Gel bands were excised and washed sequentially with HPLC grade water followed by 1:1 (v/v) MeCN/H_2_O. Gel bands were dried (via vacuum centrifuge), treated with 10 mM dithiothreitol (DTT) in 100mM NH_4_HCO_3_ and incubated for 45 minutes at 56°C with shaking. DTT was removed and 55 mM iodoacetamide (in 100 mM NH_4_HCO_3_) was added and incubated for 30 minutes in the dark. All liquid was removed and gels were washed with 100 mM NH_4_HCO_3_/MeCN as above. Gels were dried and 12.5 ng/µl trypsin, chymotrypsin or alpha lytic protease was added separately and incubated overnight at 37°C. Samples were then washed and (glyco)peptides were extracted and pooled with sequential washes with 5% (v/v) formic acid (FA) in H_2_O and MeCN. Dried samples were reconstituted in 2% MeCN, 0.05% trifluoroacetic acid and run by LC-MS.

Samples were analysed using an Ultimate 3000 UHPLC coupled to an Orbitrap Q Exactive mass spectrometer (Thermo Fisher Scientific). (Glyco)peptides were loaded onto a 75 µm × 2 cm pre-column and separated on a 75 µm × 15 cm Pepmap C18 analytical column (Thermo Fisher Scientific). Buffer A was 0.1% FA in H_2_O and buffer B was 0.1% FA in 80% MeCN with 20% H_2_O. A 40-minute linear gradient (0% to 40% buffer B) was used. Data was collected in data-dependent acquisition mode with a mass range 375 to 1500 m/z and at a resolution of 70,000. For MS/MS scans, stepped HCD normalized energy was set to 27, 30 and 33% with orbitrap detection at a resolution of 35,000.

Glycopeptide data was analysed with Byonic (Protein Metrics). Digestion was set to RK, TASV and FYWML for trypsin, alpha-lytic and chymotrypsin digests, respectively and fully specific with a maximum of two miss cleavages allowed. Carbamidomethylation (57.02 Da) was set as a fixed modification, while methionine oxidation (15.99 Da), deamidation (0.98 Da) and Gln -> pyro-glutamate (−17.03 Da) were set as variable modifications. The Byonic in-built common human N-linked (182 glycans), and O-linked glycan (9 glycans) databases were used to identify glycopeptides for RBD_HEK_. The Byonic in-built yeast glycan library was used to identify glycopeptides for Abdala, containing high mannose structures with and without phosphate groups and O-mannosylation. Byonic output files were imported into Byologic for quantification (Protein Metrics). A minimum Byonic threshold score of 300 was used for glycopeptide identification. All glycopeptide assignments were manually validated. For quantification, the extracted ion chromatogram intensities for each glycopeptide were summed and plotted relative to the total intensity for each glycosite.

### Mass Photometry

Mass photometry measurements were conducted using a Refeyn TwoMP system (Refeyn Ltd) as previously described ^24^. High Precision No. 1.5H glass coverslips were cleaned via sonication in Milli-Q H_2_O, followed by isopropanol and Milli-Q H_2_0 then dried under nitrogen flow. Sample chambers were assembled using silicone gaskets (CultureWell™ reusable gasket, 3mm diameter x 1 mm depth, Grace Bio-Labs). Coverslips were placed on the MP sample stage and a single gasket was filled with 5-20 µL DPBS (without calcium, without magnesium, pH 7.4 ThermoFisher Scientific) to find focus and ensure low background signal-to-noise. For interaction experiments, proteins were mixed at a 1:1 molar ratio and equilibrated for 5 minutes prior to data acquisition.

Acquisition settings within AcquireMP (v2.5.0, Refeyn Ltd) were as follows: small field of view, frame binning = 14, frame rate = 685.0 Hz, pixel binning = 6, exposure time 1.41 ms and movies were taken over 60 seconds. Mass calibration was conducted using an in-house protein standard. Data was analysed using DiscoverMP (v2.5.0, Refeyn Ltd). Molecule counts were used to determine levels of ACE2-RBD occupancy. The interaction between RBD and ACE2 is represented as % total occupancy, which was calculated using the sum of all ACE2 counts including species with 0 and 1 RBD molecules bound and expressed as a percentage of RBD bound counts compared to total ACE2 counts. The interaction between RBD and RBM targeting mAb is represented as % total occupancy, which was calculated using the sum of all mAb counts including species with 0, 1 and 2 RBD molecules bound and expressed as a percentage of 1 and 2 RBD bound counts compared to total mAb counts. Calculation of approximate K_d_ values was done as previously described ^24,33^. Representative histograms with overlaid kernel density estimates were generated in R (v4.2.1) using event exports from DiscoverMP.

### Hydrogen-deuterium exchange (HDX)-MS

Prior to conducting HDX-MS experiments, peptides were identified by digesting RBD_HEK_ and Abdala using the same protocol and identical liquid chromatographic (LC) gradient as detailed below and performing MSE analysis with a Synapt G2-Si mass spectrometer (Waters), applying collision energy ramping from 20 to 30 kV. Sodium iodide was used for calibration and leucine enkephalin was applied for mass accuracy correction. MSE runs were analysed with ProteinLynx Global Server (PLGS) 3.0 (Waters) and peptides identified in 3 out of 4 runs, with at least 0.2 fragments per amino acid (at least 2 fragments in total) and at least 1 consecutive product, with mass error below 10 ppm were selected in DynamX 3.0 (Waters).

HDX was conducted with differential temperature labelling, in a similar manner as previously described ^54,55^. RBD_HEK_ and Abdala (20 uM) were diluted 1:20 in deuterated PBS buffer (95% final D2O fraction, pH read 7.3) and the exchange reaction was conducted for 10 s on ice, for 10 s, 100 s (1 min 40 s), 1,000 s (16 min 40 s) and 10,000 s (2 h 46 min 40 s) at 23°C, and for 9 h at 28°C. The exchange reactions were quenched by 1:1 dilution into ice-cold 100 mM phosphate buffer containing 3 M Urea and 70 mM TCEP (final pHread 2.3). Samples were held on ice for 30 s and snap-frozen in liquid nitrogen and kept frozen at –80°C until LC-MS analysis. Triplicates were performed for time points 10 s on ice and 9 h at 28°C; duplicates were performed for all the other time points. Maximally labelled samples were produced by labelling RBDs with 3 M fully deuterated urea in D_2_O and 4 mM tris(2-carboxyethyl) phosphine (TCEP), resulting in a final deuterium content as for the other labelled samples. The maximally labelled samples were quenched after 6 h by 1:1 dilution with ice-cold 100 mM phosphate buffer (final pH read 2.3), held for 30 s on ice and snap-frozen in liquid nitrogen and kept frozen at -80°C until LC-MS analysis.

Frozen protein samples were quickly thawed and injected into an Acquity UPLC M-Class System with HDX Technology (Waters). Proteins were on-line digested at 20°C into a home-made Pepsin column and trapped/desalted with solvent A (0.23% formic acid in water, pH 2.5) for 3 min at 200 μL/min and at 0 °C through an Acquity BEH C18 VanGuard pre-column (1.7 μm, 2.1 mm × 5 mm, Waters). Peptides were eluted into an Acquity UPLC BEH C18 analytical column (1.7 μm, 2.1 mm × 100 mm, Waters) with a 7 min-linear gradient raising from 8 to 35% of solvent B (0.23% formic acid in acetonitrile) at a flow rate of 40 μL/min and at 0 °C. Then, peptides went through electrospray ionization in positive mode and underwent MS analysis with ion mobility separation.

Peptide level deuterium uptake was calculated with DynamX 3.0 and data visually inspected and curated. The threshold for the statistically significant difference in HDX was established at the significance level of 99%, based on an approach described earlier ^56^.

## DATA AVAILABILITY

Mass spectrometry raw data have been deposited to the ProteomeXchange Consortium (http://proteomecentral.proteomexchange.org) via the PRIDE partner repository with the dataset identifier <PXD>.^57^ All other data included in the manuscript and supplemental materials is available upon request.

## ACKNOWLEDGEMENTS

W.B.S. and S.A.B. recognise funding from the UKRI Future Leaders Fellowship MR/V02213*X*/1 and the University of Oxford’s COVID-19 Research Response Fund. We would like to thank Arthur Huang (Chang Gung Memorial Hospital, Taiwan), Tiong Tan, Lisa Schimanski and Alain Townsend (MRC Weatherall Institute of Molecular Medicine, University of Oxford) for providing a SARS-CoV-2 spike receptor-binding motif (RBM) targeting antibody.

## COMPETING INTERESTS

W.B.S. is a shareholder of Refeyn Ltd.

