## Supplemental Information for "Structural and Functional Glycosylation of the Abdala COVID-19 Vaccine"

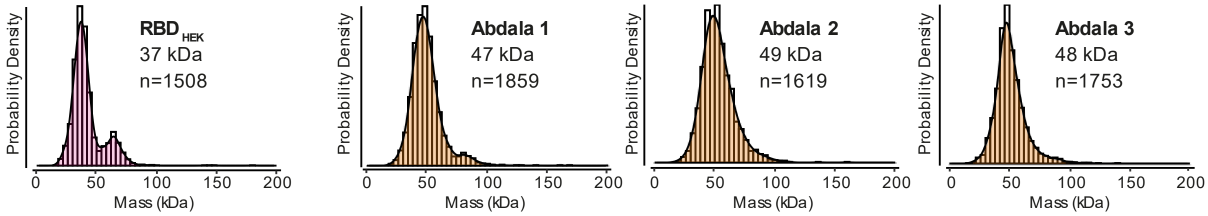

**Supplementary Figure 1. Mass photometry of RBD$_{HEK}$ and Abdala from three production batches**. The number of detected particles are also given (n).

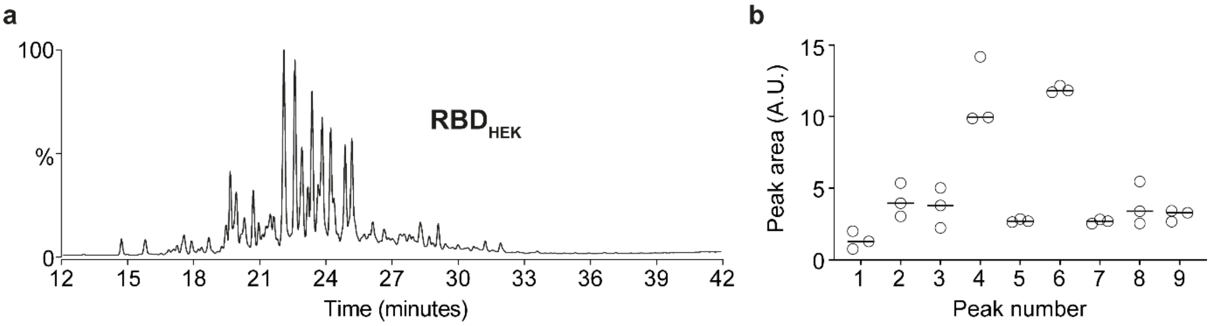

**Supplementary Figure 2. UPLC-based characterisation of RBD$_{HEK}$ and Abdala N-glycans.** (**a**) UPLC profile of RBD$_{HEK}$ protein N-glycans. (**b**) Peak areas of the 9 most abundant Abdala UPLC glycan peaks across three production batches.

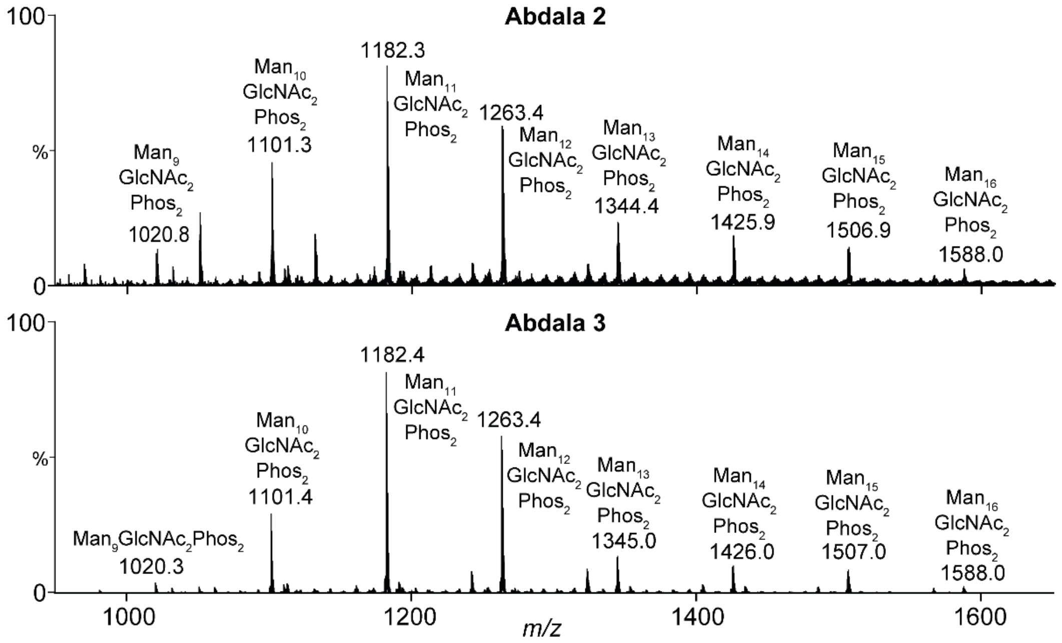

**Supplementary Figure 3. Glycan analysis of released Abdala N-glycans.** Total N-glycan MS spectrum of Abdala batch 2 and 3. Glycan compositions and ion assignments are annotated in Supplementary Tables 2-5.

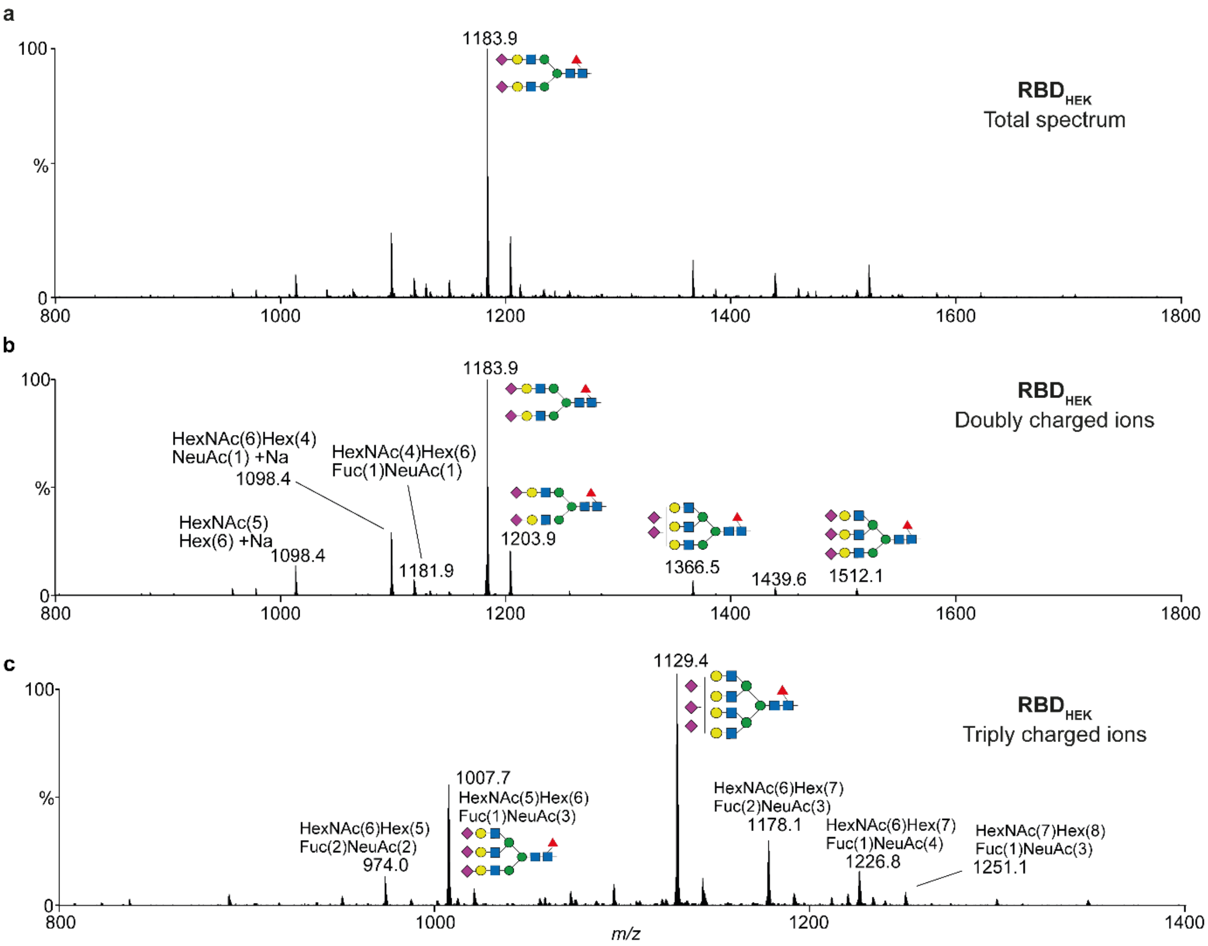

**Supplementary Figure 4. Glycan analysis of released $RBD_{HEK}$ N-glycans.** (**a**) N-glycan MS spectra of $RBD_{HEK}$ (no IM extraction). (**b**) Ion mobility extracted MS spectrum of doubly charged ions. (**c**) Ion mobility extracted MS spectrum of triply charged ions.

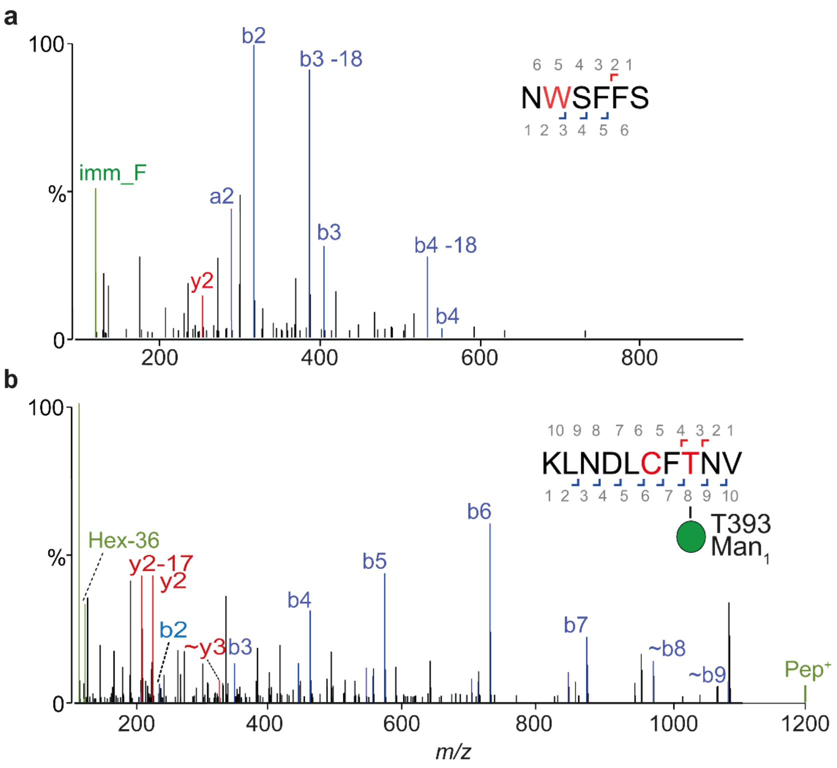

**Supplementary Figure 5.** (**a**) MS/MS spectrum of the non-glycosylated N-terminal extension of Abdala. (**b**) MS/MS spectrum of O-mannosylation at position T393.

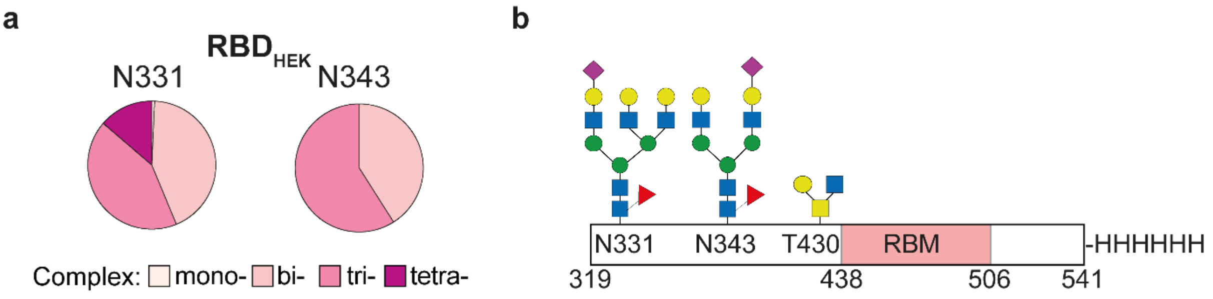

**Supplementary Figure 6. Glycoproteomic characterisation of HEK-derived RBD.** (**a**) Pie charts showing site-specific glycosylation at N331 and N343. N-glycans are group based on complex-type branching (i.e. mono- bi- tri- and tetra-antennary structures). (**b**) Schematic representation of the RBD amino acid sequence, showing location of N- and O-glycosylation sites.

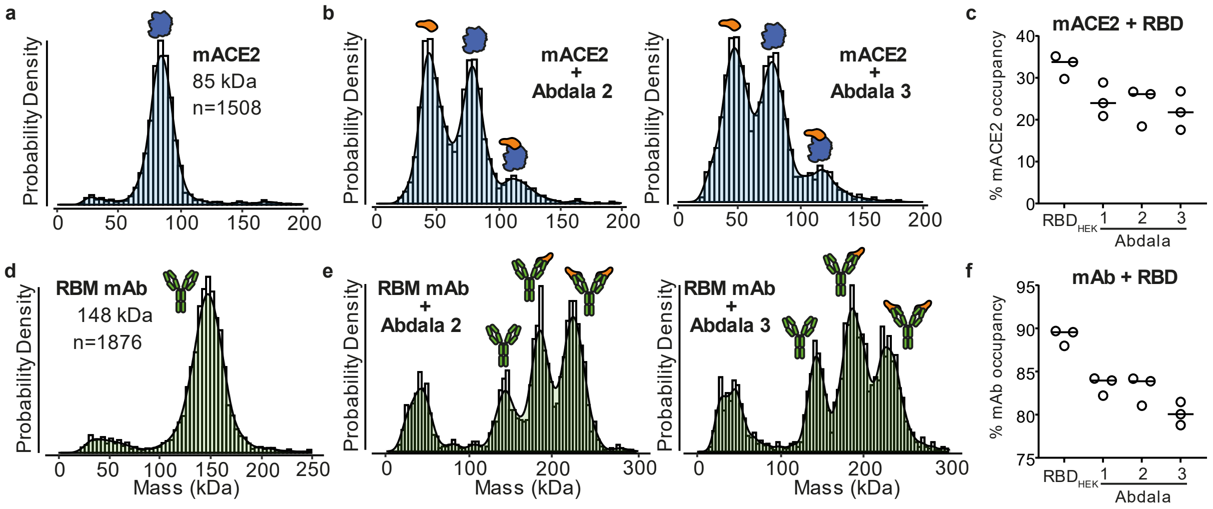

**Supplementary Figure 7. Mass photometry of Abdala and RBD$_{HEK}$ with ACE2 and IgG.** (**a**) MP of monomeric ACE2 (mACE2) alone and (**b**) with Abdala RBD batches 2 and 3 at a 1:1 molar ratio. (**c**) The interaction between mACE2 and RBD expressed as percentage occupancy accounting for unbound and bound mACE2 counts. (**d**) MP of RBM-targeting mAb alone and (**e**) with Abdala RBD batches 2 and 3 at a 1:1 molar ratio. (**f**) The interaction between the RBM-targeting mAb and RBD expressed as percentage occupancy accounting for unbound and bound mAb counts.

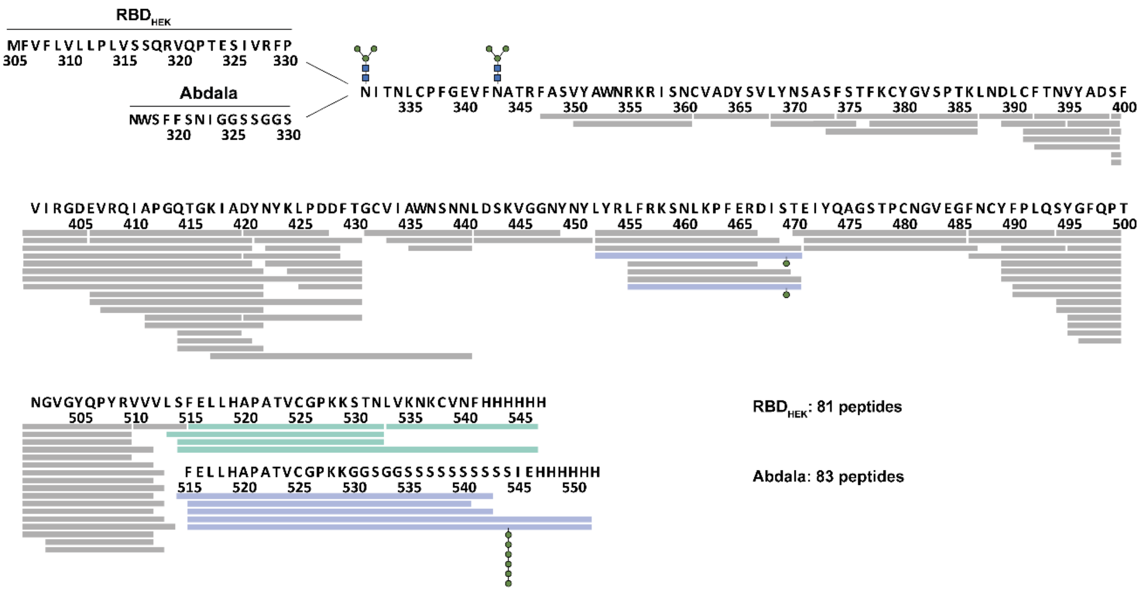

**Supplementary Figure 8. Effective HDX-MS sequence coverage of RBD$_{HEK}$ and Abdala.** Peptide HDX represented along the protein sequences: peptides in common between the two forms (grey), peptides belonging uniquely to HEK-derived RBD (green), peptides belonging uniquely to Abdala (blue). Positions of N-glycans (N331 and N344) and O-glycans (S469 and S544) are also schematically represented.

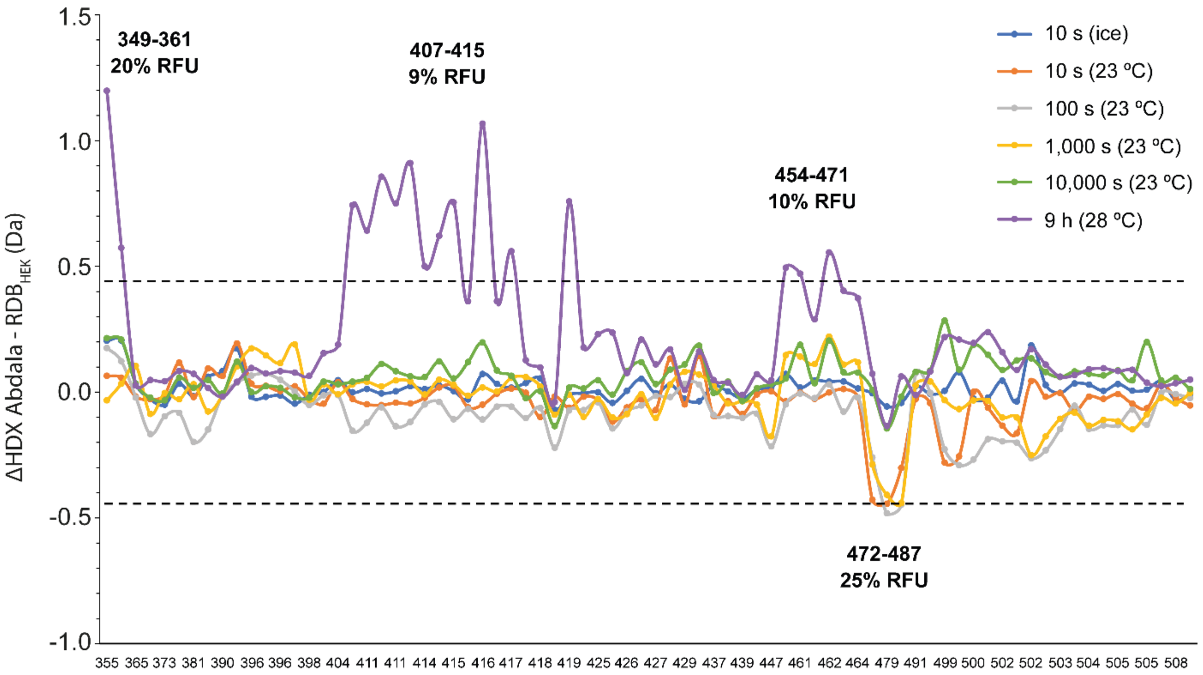

**Supplementary Figure 9. HDX peptide-level difference plot between Abdala and RBD$_{HEK}$ over the measured time points.** Peptides are arranged based on their central residue according to their position from the N- and C-terminus. The dotted grey lines indicate the threshold of significance (±0.44 Da) calculated at 99% confidence interval. The four regions whose peptides had significant differences in HDX are specified (bold text) with their cumulative difference in relative fractional uptake (RFU).

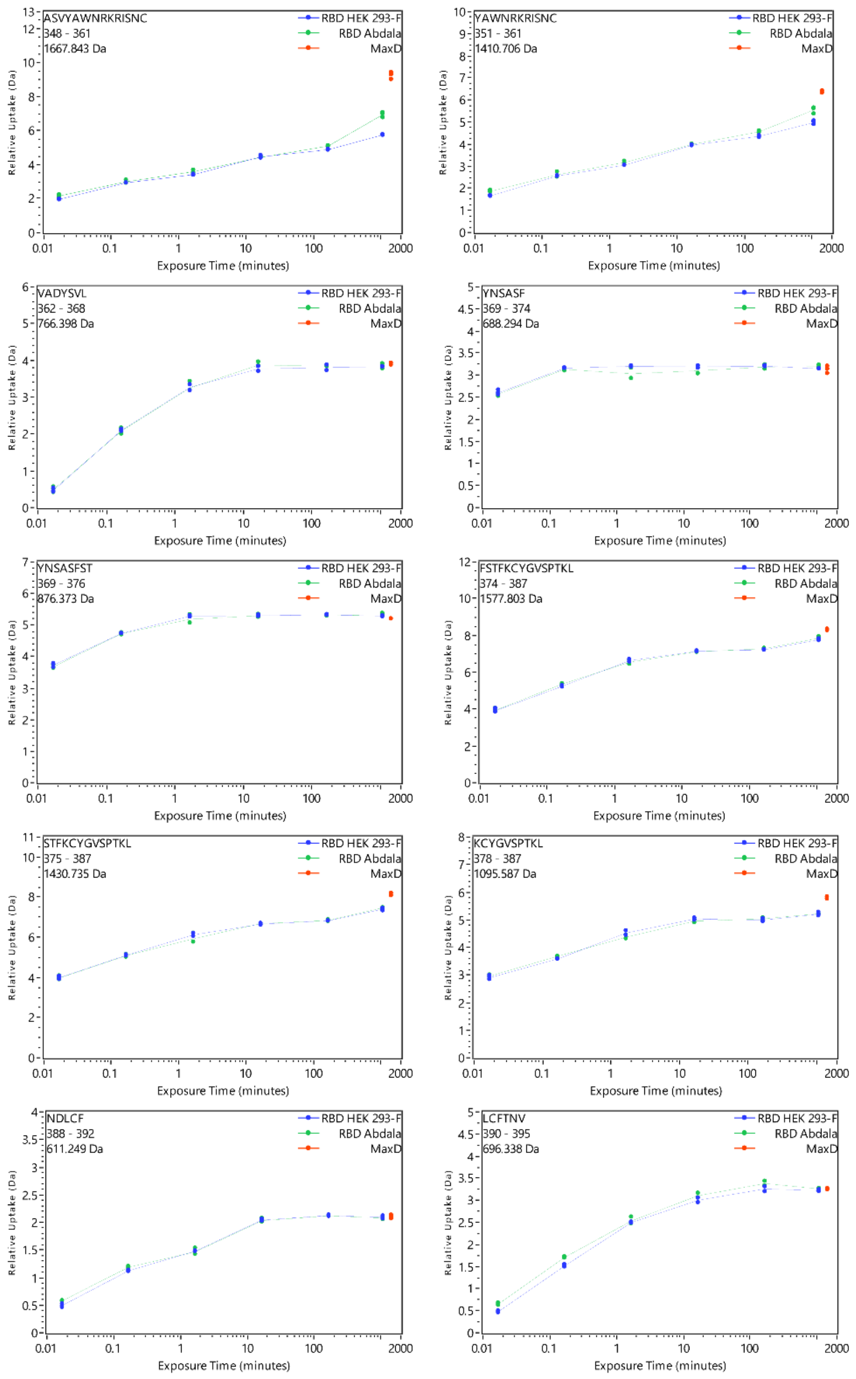
ASVYAWNRKRISNC
348 - 361
1667.843 Da
YAWNRKRISNC
351 - 361
1410.706 Da
VADYSVL
362 - 368
766.398 Da
YNSASF
369 - 374
688.294 Da
YNSASFST
369 - 376
876.373 Da
FSTFKCYGVSPTKL
374 - 387
1577.803 Da
STFKCYGVSPTKL
375 - 387
1430.735 Da
KCYGVSPTKL
378 - 387
1095.587 Da
NDLCF
388 - 392
611.249 Da
LCFTNV
390 - 395
696.338 Da
RBD HEK 293-F
RBD Abdala
MaxD
Relative Uptake (Da)
Exposure Time (minutes)
0.01
0.1
1
10
100
2000

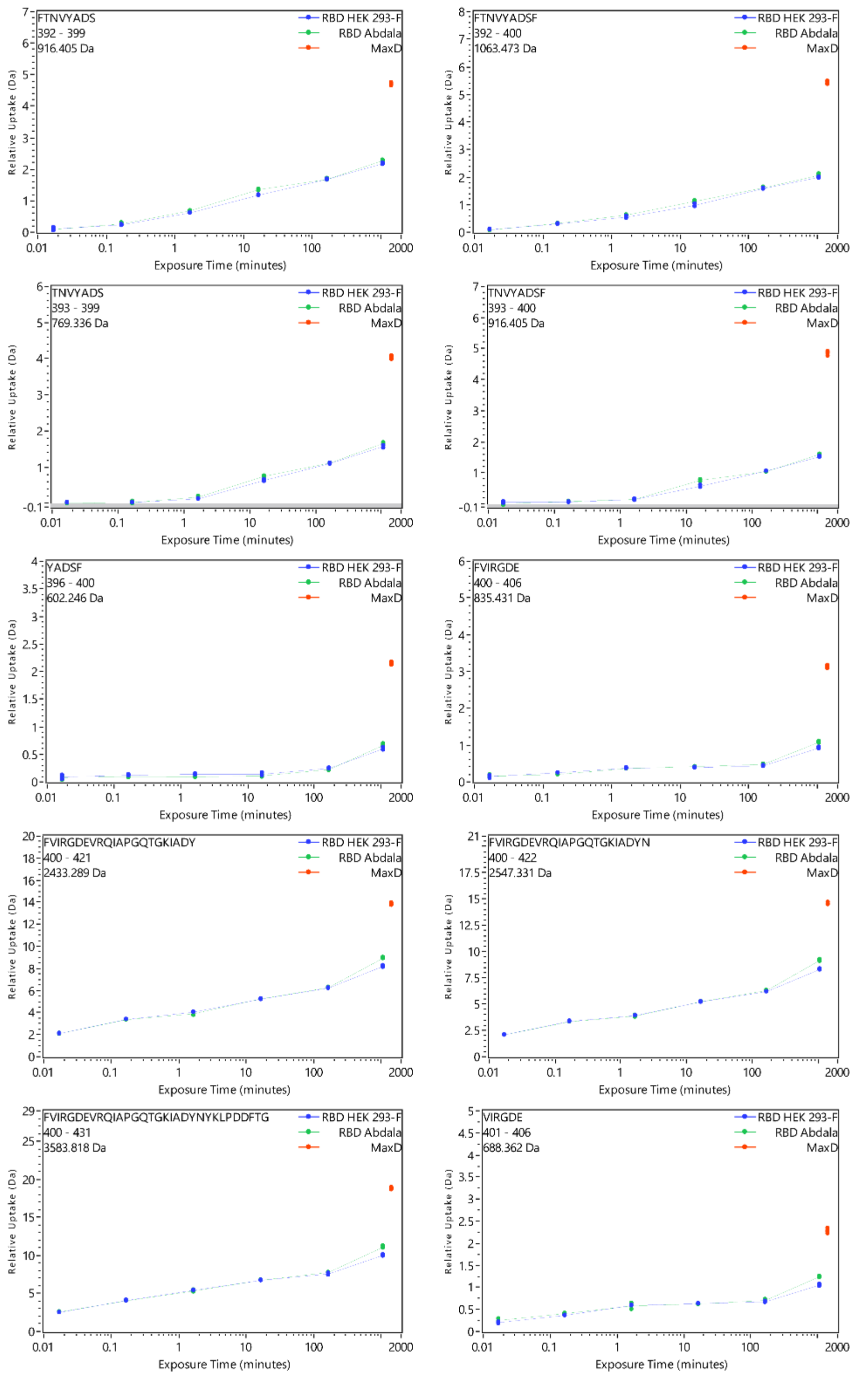
FTNVYADS
392 - 399
916.405 Da
RBD HEK 293-F
RBD Abdala
MaxD
Relative Uptake (Da)
Exposure Time (minutes)
FTNVYADSF
392 - 400
1063.473 Da
TNVYADS
393 - 399
769.336 Da
TNVYADSF
393 - 400
916.405 Da
YADSF
396 - 400
602.246 Da
FVIRGDE
400 - 406
835.431 Da
FVIRGDEVRQIAPGQTGKIADY
400 - 421
2433.289 Da
FVIRGDEVRQIAPGQTGKIADYN
400 - 422
2547.331 Da
FVIRGDEVRQIAPGQTGKIADYNYKLPDDFTG
400 - 431
3583.818 Da
VIRGDE
401 - 406
688.362 Da

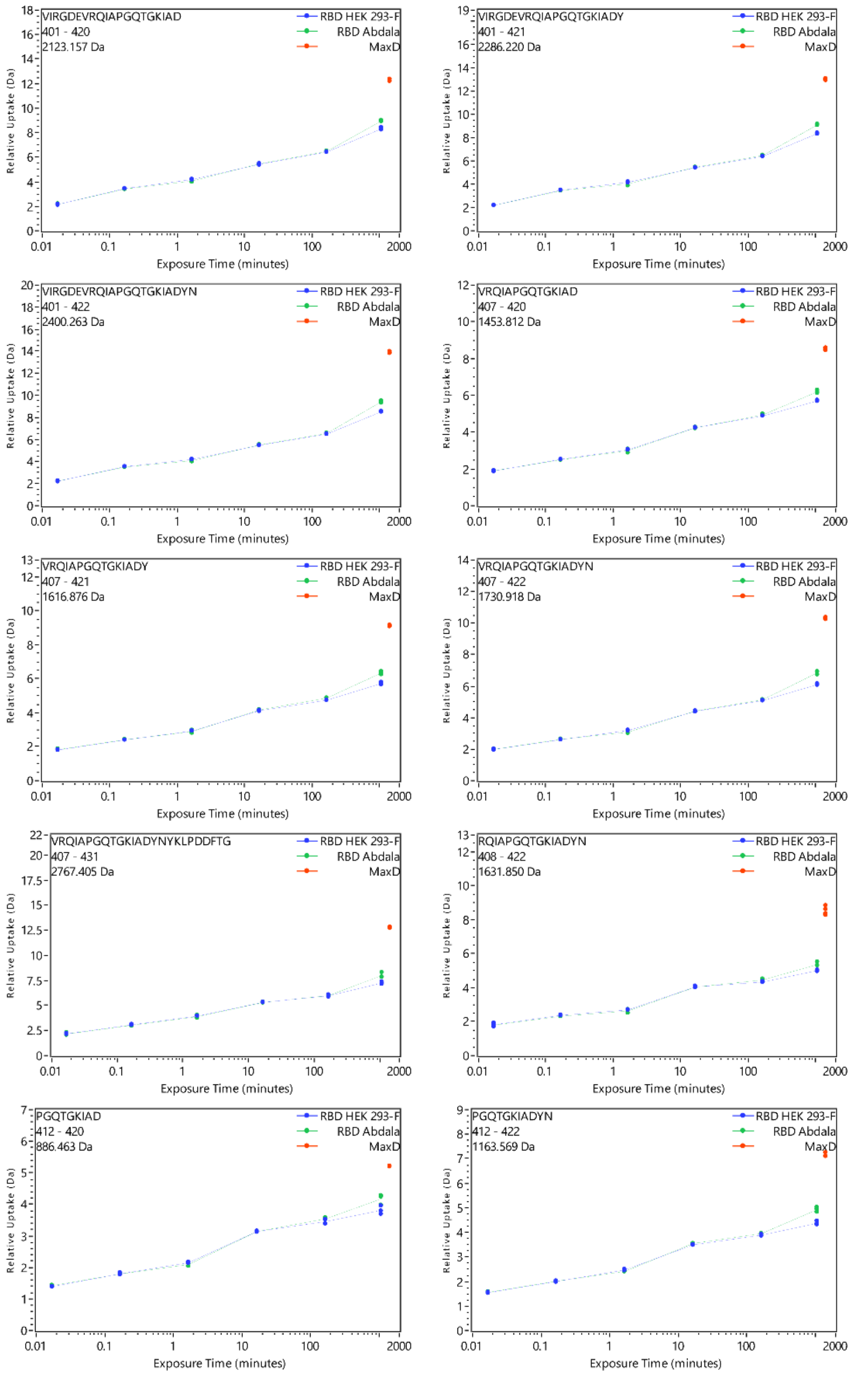
VIRGDEVRQIAPGQTGKIAD
401 - 420
2123.157 Da
VIRGDEVRQIAPGQTGKIADY
401 - 421
2286.220 Da
VIRGDEVRQIAPGQTGKIADYN
401 - 422
2400.263 Da
VRQIAPGQTGKIAD
407 - 420
1453.812 Da
VRQIAPGQTGKIADY
407 - 421
1616.876 Da
VRQIAPGQTGKIADYN
407 - 422
1730.918 Da
VRQIAPGQTGKIADYNYKLPDDFTG
407 - 431
2767.405 Da
RQIAPGQTGKIADYN
408 - 422
1631.850 Da
PGQTGKIAD
412 - 420
886.463 Da
PGQTGKIADYN
412 - 422
1163.569 Da
RBD HEK 293-F
RBD Abdala
MaxD
Relative Uptake (Da)
Exposure Time (minutes)
0.01
0.1
1
10
100
2000

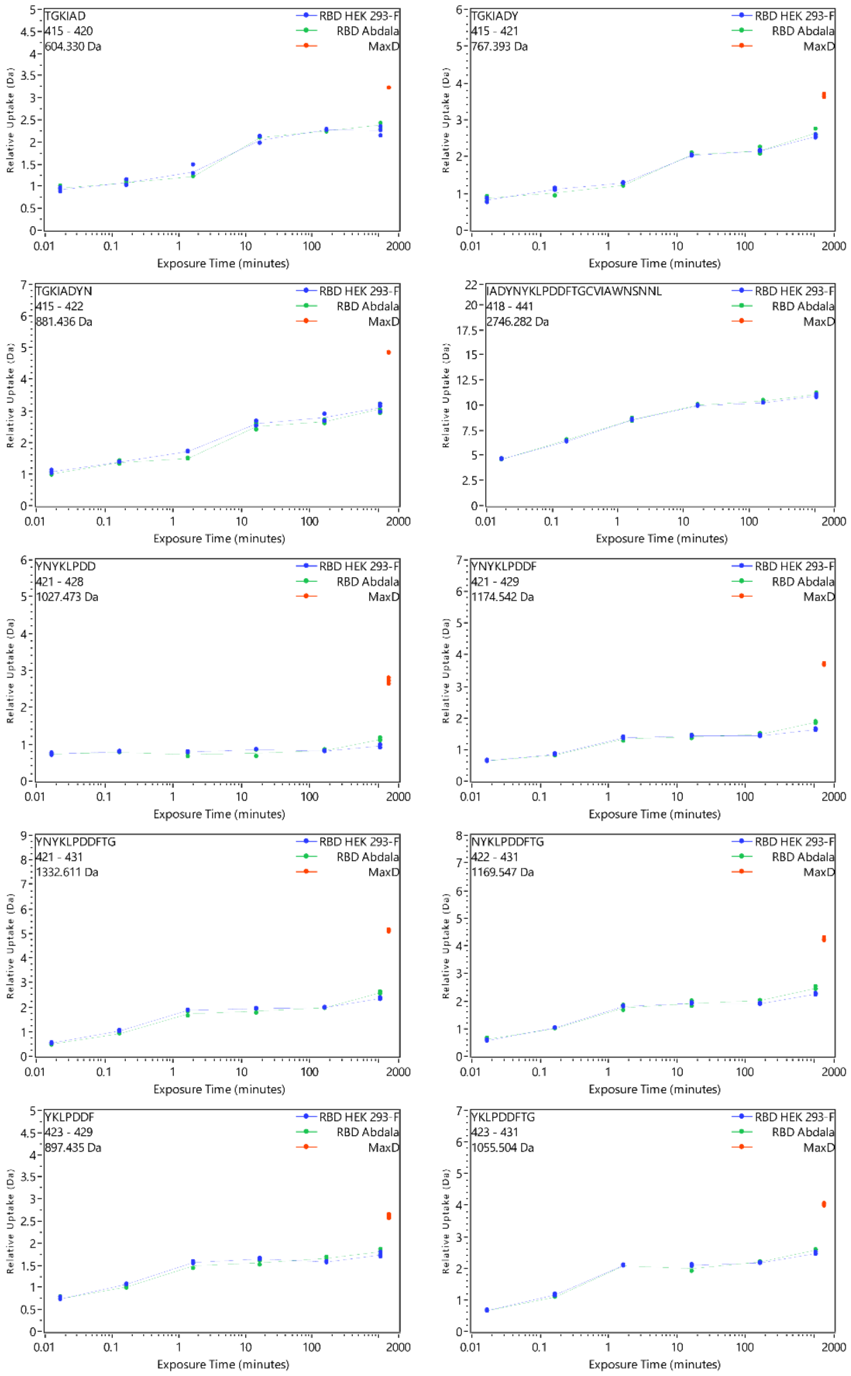
TGKIAD
415 - 420
604.330 Da
RBD HEK 293-F
RBD Abdala
MaxD
Relative Uptake (Da)
Exposure Time (minutes)
TGKIADY
415 - 421
767.393 Da
TGKIADYN
415 - 422
881.436 Da
IADYNYKLPDDFTGCVIAWNSNNL
418 - 441
2746.282 Da
YNYKLPDD
421 - 428
1027.473 Da
YNYKLPDDF
421 - 429
1174.542 Da
YNYKLPDDFTG
421 - 431
1332.611 Da
NYKLPDDFTG
422 - 431
1169.547 Da
YKLPDDF
423 - 429
897.435 Da
YKLPDDFTG
423 - 431
1055.504 Da

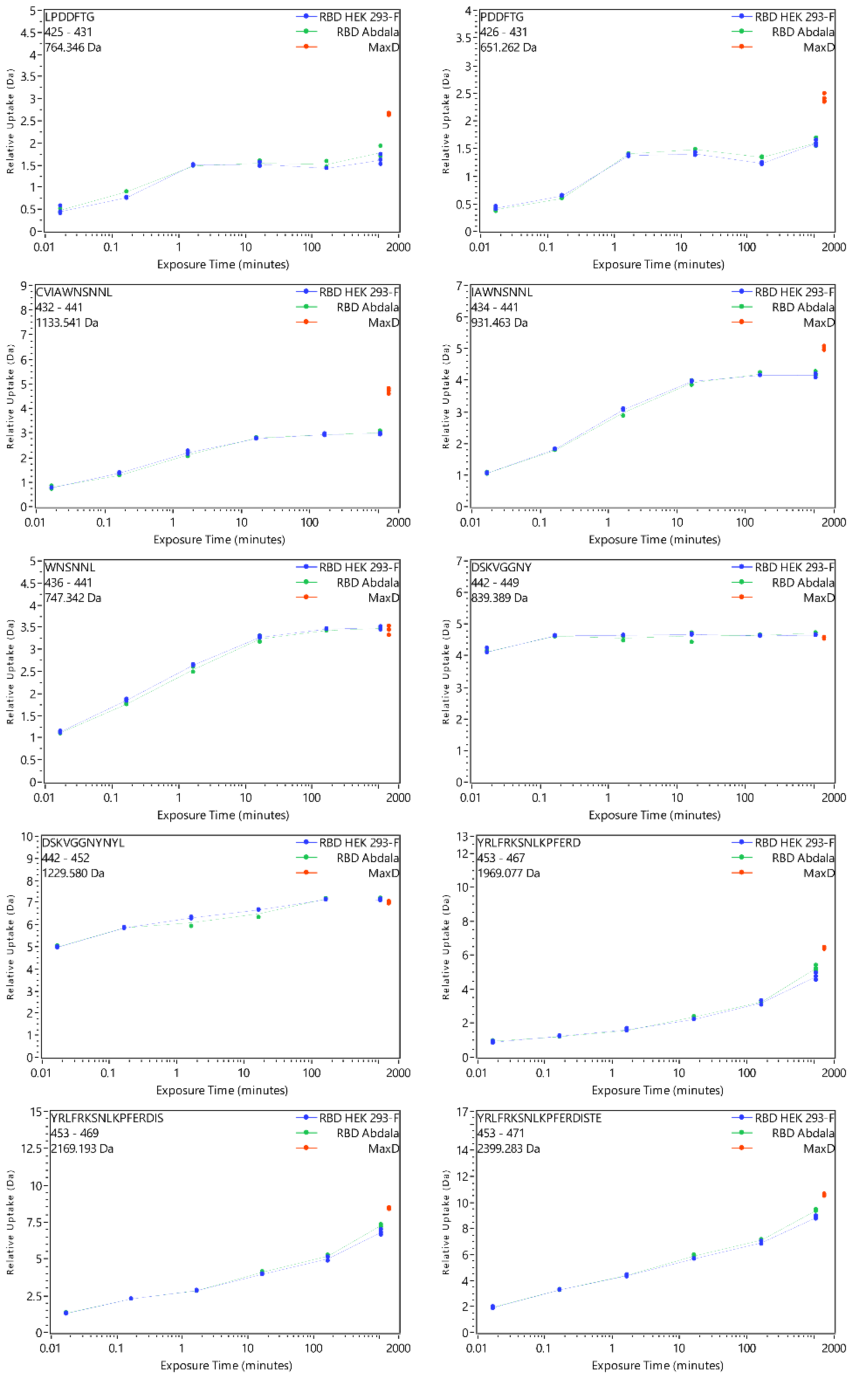
LPDDFTG
425 - 431
764.346 Da
PDDFTG
426 - 431
651.262 Da
CVIAWNSNNL
432 - 441
1133.541 Da
IAWNSNNL
434 - 441
931.463 Da
WNSNNL
436 - 441
747.342 Da
DSKVGGNY
442 - 449
839.389 Da
DSKVGGNYNYL
442 - 452
1229.580 Da
YRLFRKSNLKPFERD
453 - 467
1969.077 Da
YRLFRKSNLKPFERDIS
453 - 469
2169.193 Da
YRLFRKSNLKPFERDISTE
453 - 471
2399.283 Da
RBD HEK 293-F
RBD Abdala
MaxD
Relative Uptake (Da)
Exposure Time (minutes)
0.01
0.1
1
10
100
2000

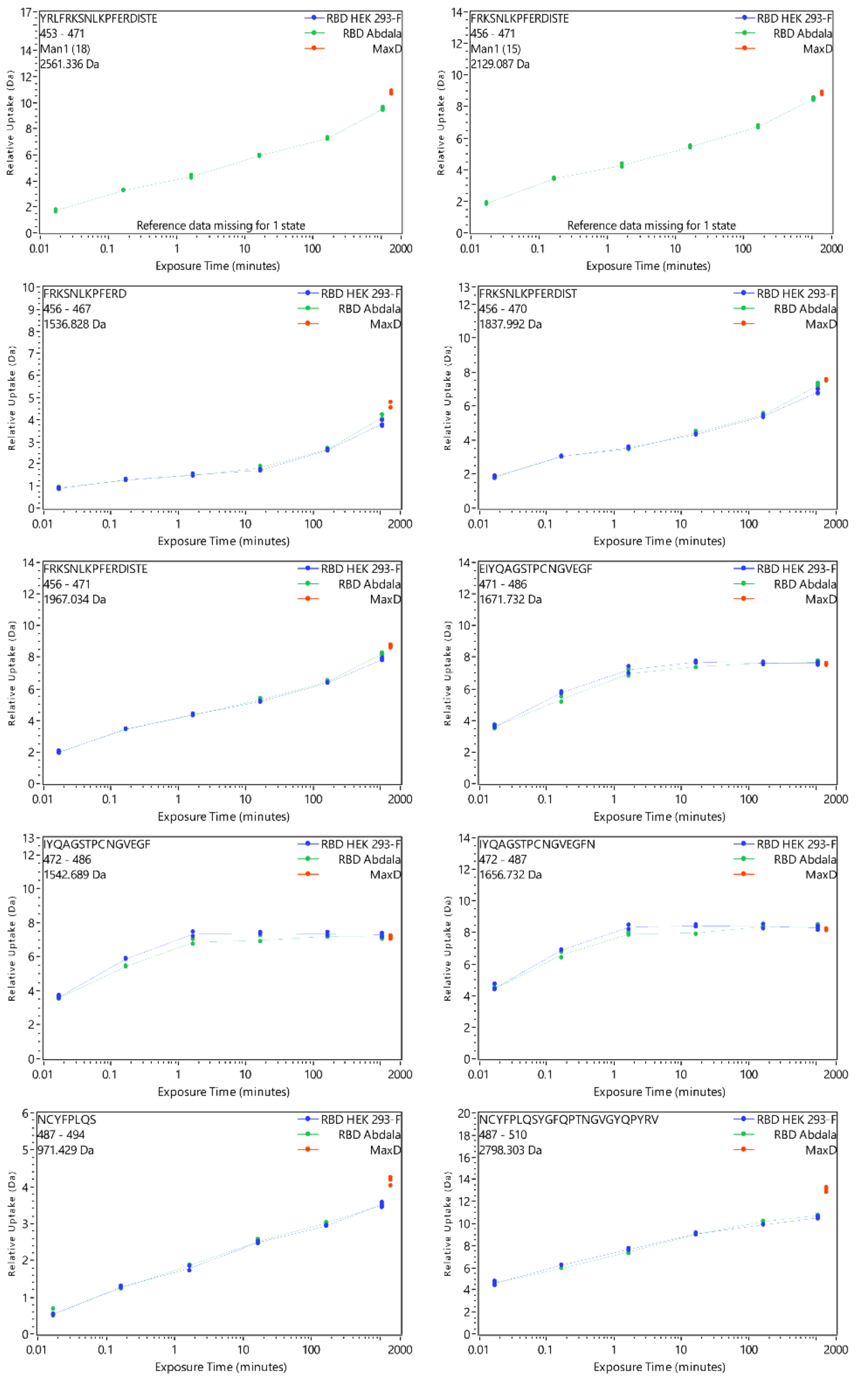
YRLFRKSNLKPFERDISTE
453 - 471
Man1 (18)
2561.336 Da
RBD HEK 293-F
RBD Abdala
MaxD
Reference data missing for 1 state
Relative Uptake (Da)
Exposure Time (minutes)
FRKSNLKPFERDISTE
456 - 471
Man1 (15)
2129.087 Da
RBD HEK 293-F
RBD Abdala
MaxD
Reference data missing for 1 state
Relative Uptake (Da)
Exposure Time (minutes)
FRKSNLKPFERD
456 - 467
1536.828 Da
RBD HEK 293-F
RBD Abdala
MaxD
Relative Uptake (Da)
Exposure Time (minutes)
FRKSNLKPFERDIST
456 - 470
1837.992 Da
RBD HEK 293-F
RBD Abdala
MaxD
Relative Uptake (Da)
Exposure Time (minutes)
FRKSNLKPFERDISTE
456 - 471
1967.034 Da
RBD HEK 293-F
RBD Abdala
MaxD
Relative Uptake (Da)
Exposure Time (minutes)
EIYQAGSTPCNGVEGF
471 - 486
1671.732 Da
RBD HEK 293-F
RBD Abdala
MaxD
Relative Uptake (Da)
Exposure Time (minutes)
IYQAGSTPCNGVEGF
472 - 486
1542.689 Da
RBD HEK 293-F
RBD Abdala
MaxD
Relative Uptake (Da)
Exposure Time (minutes)
IYQAGSTPCNGVEGFN
472 - 487
1656.732 Da
RBD HEK 293-F
RBD Abdala
MaxD
Relative Uptake (Da)
Exposure Time (minutes)
NCYFPLQS
487 - 494
971.429 Da
RBD HEK 293-F
RBD Abdala
MaxD
Relative Uptake (Da)
Exposure Time (minutes)
NCYFPLQSYGFQPTNGVGYQPYRV
487 - 510
2798.303 Da
RBD HEK 293-F
RBD Abdala
MaxD
Relative Uptake (Da)
Exposure Time (minutes)

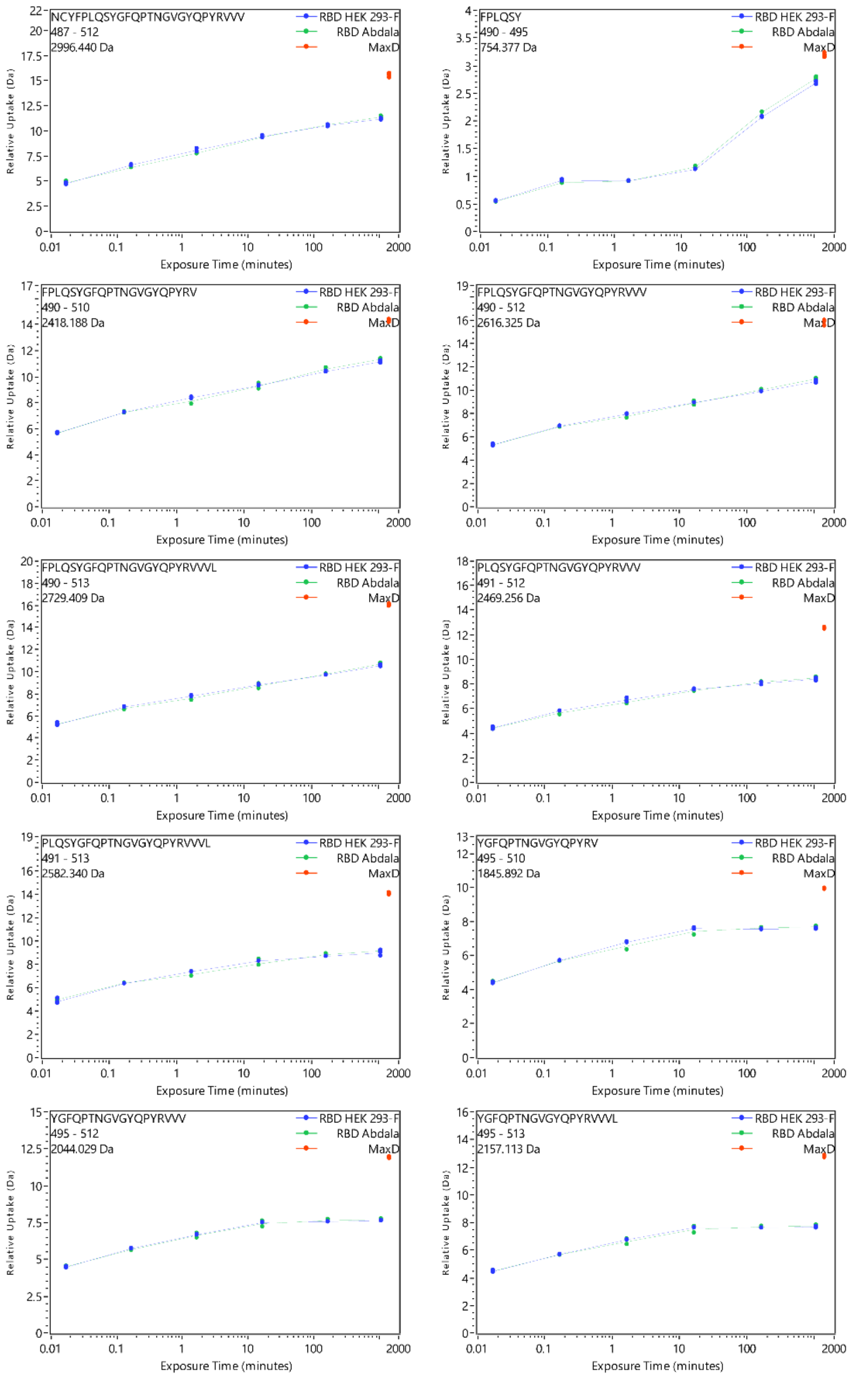
NCYFPLQSYGFQPTNGVGYQPYRVVV
487 - 512
2996.440 Da
FPLQSY
490 - 495
754.377 Da
FPLQSYGFQPTNGVGYQPYRV
490 - 510
2418.188 Da
FPLQSYGFQPTNGVGYQPYRVVV
490 - 512
2616.325 Da
FPLQSYGFQPTNGVGYQPYRVVVL
490 - 513
2729.409 Da
PLQSYGFQPTNGVGYQPYRVVV
491 - 512
2469.256 Da
PLQSYGFQPTNGVGYQPYRVVVL
491 - 513
2582.340 Da
YGFQPTNGVGYQPYRV
495 - 510
1845.892 Da
YGFQPTNGVGYQPYRVVV
495 - 512
2044.029 Da
YGFQPTNGVGYQPYRVVVL
495 - 513
2157.113 Da
RBD HEK 293-F
RBD Abdala
MaxD
Relative Uptake (Da)
Exposure Time (minutes)

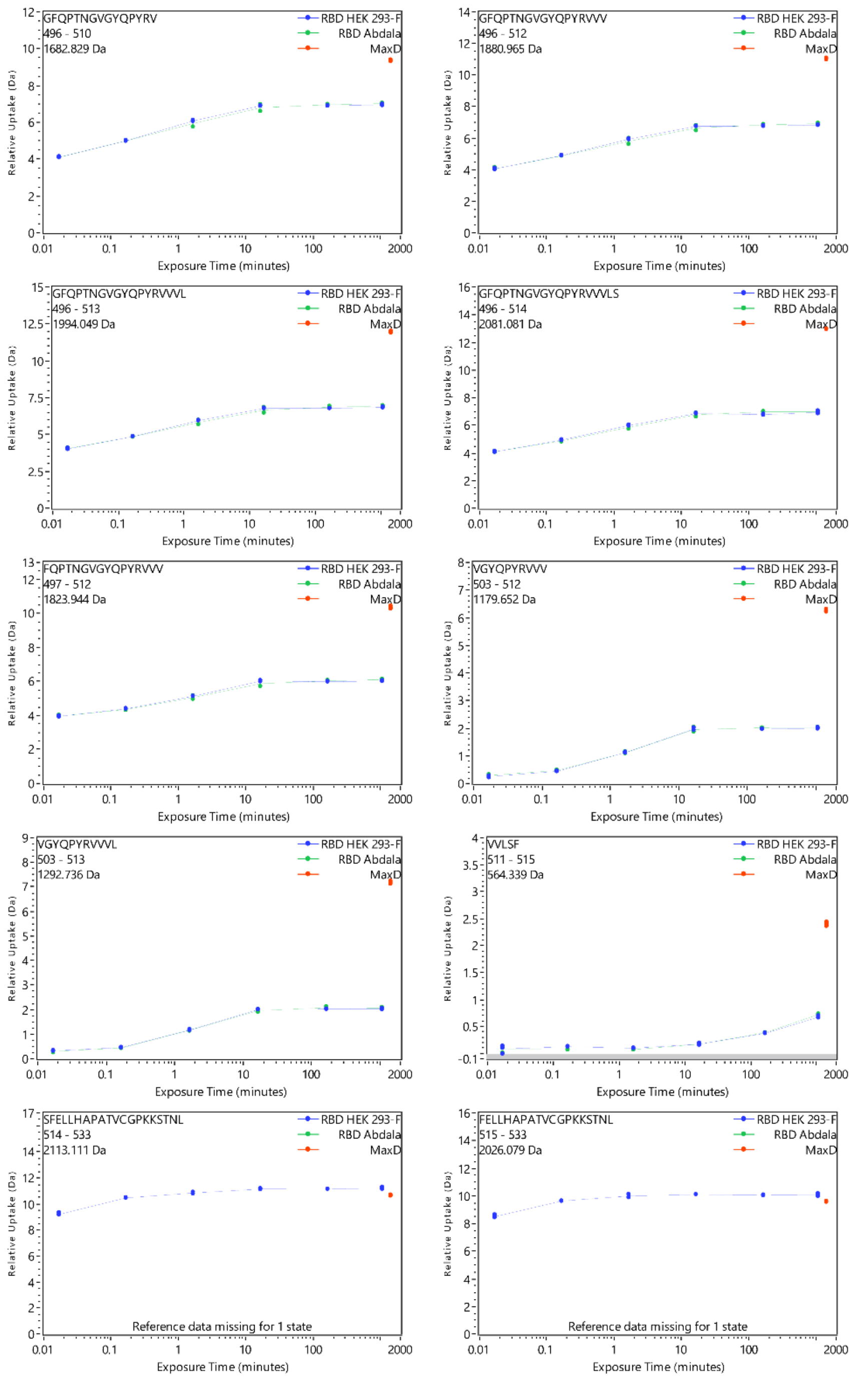
GFQPTNGVGYQPYRV
496 - 510
1682.829 Da
RBD HEK 293-F
RBD Abdala
MaxD
Relative Uptake (Da)
Exposure Time (minutes)
GFQPTNGVGYQPYRVVV
496 - 512
1880.965 Da
GFQPTNGVGYQPYRVVVL
496 - 513
1994.049 Da
GFQPTNGVGYQPYRVVVLS
496 - 514
2081.081 Da
FQPTNGVGYQPYRVVV
497 - 512
1823.944 Da
VGYQPYRVVV
503 - 512
1179.652 Da
VGYQPYRVVVL
503 - 513
1292.736 Da
VVLSF
511 - 515
564.339 Da
SFELLHAPATVCGPKKSTNL
514 - 533
2113.111 Da
Reference data missing for 1 state
FELLHAPATVCGPKKSTNL
515 - 533
2026.079 Da
Reference data missing for 1 state

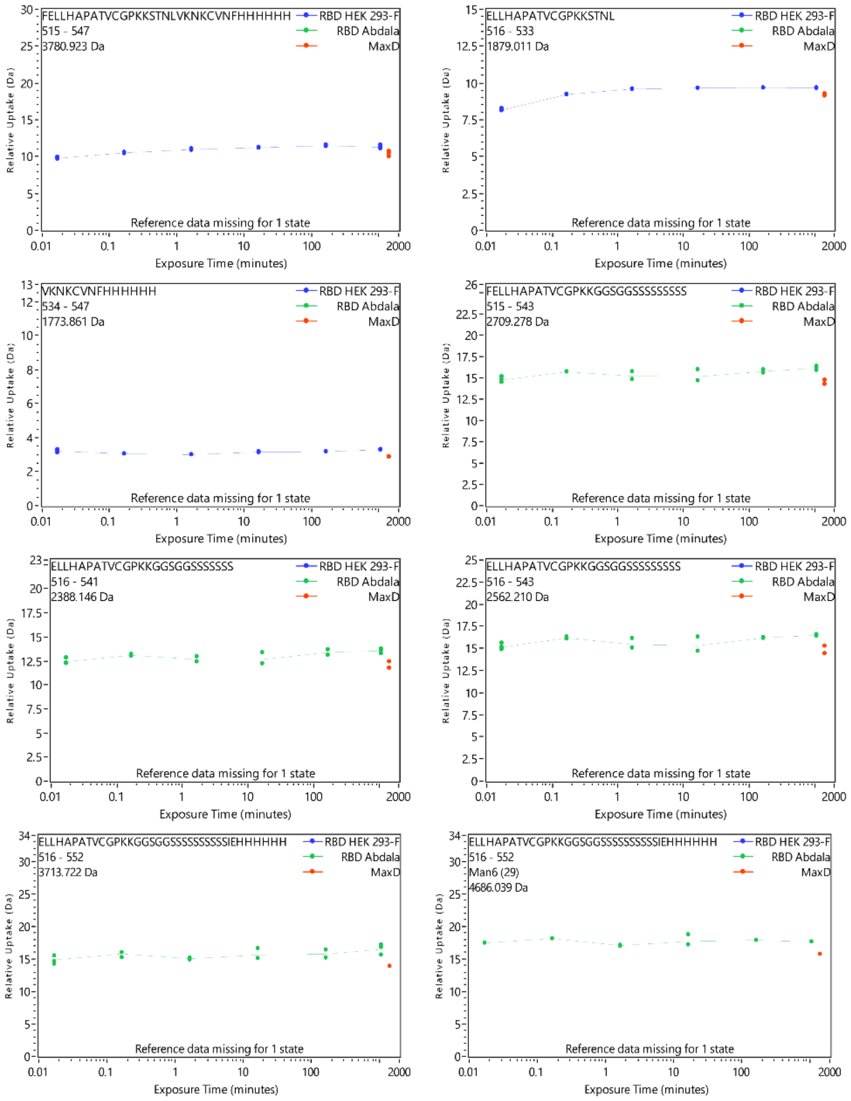

**Supplementary Figure 10. Deuterium uptake plots of $RBD_{HEK}$ and Abdala.** HDX of HEK-derived RBD and Abdala across all measured time points (represented separately for each peptide).

| | Peak area (A.U.) | | |
|---|---|---|---|
| **HPLC Peak** | **Abdala 1** | **Abdala 2** | **Abdala 3** |
| 1 | 0.76 | 2.00 | 1.29 |
| 2 | 3.04 | 5.36 | 3.96 |
| 3 | 2.23 | 5.03 | 3.80 |
| 4 | 14.18 | 9.95 | 9.87 |
| 5 | 2.86 | 2.63 | 2.71 |
| 6 | 12.16 | 11.81 | 11.69 |
| 7 | 2.54 | 2.71 | 2.83 |
| 8 | 5.47 | 2.55 | 3.40 |
| 9 | 3.43 | 2.67 | 3.28 |

**Supplementary Table. 1**: HPLC peak areas for the 9 major N-glycans from three production batches of Abdala.

| *m/z* | | Composition | | | | Ion |
|---|---|---|---|---|---|---|
| Found | Calc. | Man | GlcNAc | Phosphate | Na | |
| 1313.4 | 1313.4 | 5 | 2 | 1 | 0 | $[M – H]^-$ |
| 1475.4 | 1475.5 | 6 | 2 | 1 | 0 | $[M – H]^-$ |
| 1637.5 | 1637.6 | 7 | 2 | 1 | 0 | $[M – H]^-$ |
| 1799.5 | 1799.6 | 8 | 2 | 1 | 0 | $[M – H]^-$ |
| 1961.6 | 1961.6 | 9 | 2 | 1 | 0 | $[M – H]^-$ |
| 2041.6 | 2041.6 | 9 | 2 | 2 | 0 | $[M – H]^-$ |
| 2063.5 | 2063.6 | 9 | 2 | 2 | 1 | $[M – H + Na]^-$ |
| 2123.6 | 2123.7 | 10 | 2 | 1 | 0 | $[M – H]^-$ |
| 2203.6 | 2203.7 | 10 | 2 | 2 | 0 | $[M – H]^-$ |
| 2225.6 | 2225.7 | 10 | 2 | 2 | 1 | $[M – H + Na]^-$ |
| 2285.7 | 2285.7 | 11 | 2 | 1 | 0 | $[M – H]^-$ |
| 2365.7 | 2365.7 | 11 | 2 | 2 | 0 | $[M – H]^-$ |
| 2387.7 | 2387.7 | 11 | 2 | 2 | 1 | $[M – H + Na]^-$ |
| 2447.8 | 2447.8 | 12 | 2 | 1 | 0 | $[M – H]^-$ |
| 2549.7 | 2549.8 | 12 | 2 | 2 | 1 | $[M – H + Na]^-$ |
| 2609.8 | 2609.8 | 13 | 2 | 1 | 0 | $[M – H]^-$ |
| 2711.8 | 2711.8 | 13 | 2 | 2 | 1 | $[M – H + Na]^-$ |
| 2771.8 | 2771.9 | 14 | 2 | 1 | 0 | $[M – H]^-$ |
| 2872.8 | 2873.8 | 14 | 2 | 2 | 1 | $[M – H + Na]^-$ |

**Supplementary Table. 2**: Abdala N-glycan compositions from ion mobility extracted singly charged ions with the identified molecular ion

| *m/z* | | Composition | | | | Ion |
|---|---|---|---|---|---|---|
| Found | Calc. | Man | GlcNAc | Phosphate | Na | |
| 1020.3 | 1020.3 | 9 | 2 | 2 | 0 | $[M - H_2]^{2-}$ |
| 1080.3 | 1080.3 | 9 | 2 | 2 | 1 | $[M - H + Na + H_2PO_4]^{2-}$ |
| 1101.4 | 1101.4 | 10 | 2 | 2 | 0 | $[M - H_2]^{2-}$ |
| 1161.4 | 1161.4 | 10 | 2 | 2 | 1 | $[M - H + Na + H_2PO_4]^{2-}$ |
| 1182.4 | 1182.4 | 11 | 2 | 2 | 0 | $[M - H_2]^{2-}$ |
| 1242.4 | 1242.3 | 11 | 2 | 2 | 1 | $[M - H + Na + H_2PO_4]^{2-}$ |
| 1263.4 | 1263.4 | 12 | 2 | 2 | 0 | $[M - H_2]^{2-}$ |
| 1323.4 | 1323.3 | 12 | 2 | 2 | 1 | $[M - H + Na + H_2PO_4]^{2-}$ |
| 1344.5 | 1344.4 | 13 | 2 | 2 | 0 | $[M - H_2]^{2-}$ |
| 1404.5 | 1404.4 | 13 | 2 | 2 | 1 | $[M - H + Na + H_2PO_4]^{2-}$ |
| 1425.5 | 1425.4 | 14 | 2 | 2 | 0 | $[M - H_2]^{2-}$ |
| 1485.5 | 1485.4 | 14 | 2 | 2 | 1 | $[M - H + Na + H_2PO_4]^{2-}$ |
| 1506.5 | 1506.4 | 15 | 2 | 2 | 0 | $[M - H_2]^{2-}$ |
| 1566.5 | 1566.5 | 15 | 2 | 2 | 1 | $[M - H + Na + H_2PO_4]^{2-}$ |
| 1587.5 | 1587.5 | 16 | 2 | 2 | 0 | $[M - H_2]^{2-}$ |
| 1647.5 | 1647.5 | 16 | 2 | 2 | 1 | $[M - H + Na + H_2PO_4]^{2-}$ |
| 1668.5 | 1668.5 | 17 | 2 | 2 | 0 | $[M - H_2]^{2-}$ |
| 1749.5 | 1749.5 | 18 | 2 | 2 | 0 | $[M - H_2]^{2-}$ |
| 1830.6 | 1830.6 | 19 | 2 | 2 | 0 | $[M - H_2]^{2-}$ |
| 1911.5 | 1911.6 | 20 | 2 | 2 | 0 | $[M - H_2]^{2-}$ |
| 1992.6 | 1992.6 | 21 | 2 | 2 | 0 | $[M - H_2]^{2-}$ |

**Supplementary Table. 3**: Abdala N-glycan compositions from ion mobility extracted doubly charged ions with the identified molecular ion.

| *m/z* | | Composition | | | | Ion |
|---|---|---|---|---|---|---|
| **Found** | **Calc.** | **Man** | **GlcNAc** | **Phosphate** | **Na** | |
| 760.9 | 760.5 | 10 | 2 | 3 | 0 | $[M - H_3]^{3-}$ |
| 814.6 | 814.5 | 11 | 2 | 3 | 0 | $[M - H_3]^{3-}$ |
| 820.6 | 820.5 | 11 | 2 | 2 | 0 | $[M - H_2 + H_2PO_4]^{3-}$ |
| 854.3 | 854.5 | 11 | 2 | 3 | 1 | $[M - H_2 + Na + H_2PO_4]^{3-}$ |
| 868.6 | 868.6 | 12 | 2 | 3 | 0 | $[M - H_3]^{3-}$ |
| 874.6 | 874.6 | 12 | 2 | 2 | 0 | $[M - H_2 + H_2PO_4]^{3-}$ |
| 908.6 | 908.5 | 12 | 2 | 3 | 1 | $[M - H_2 + Na + H_2PO_4]^{3-}$ |
| 922.6 | 922.6 | 13 | 2 | 3 | 0 | $[M - H_3]^{3-}$ |
| 928.6 | 928.6 | 13 | 2 | 2 | 0 | $[M - H_2 + H_2PO_4]^{3-}$ |
| 962.6 | 962.6 | 13 | 2 | 3 | 1 | $[M - H_2 + Na + H_2PO_4]^{3-}$ |
| 976.6 | 976.6 | 14 | 2 | 3 | 0 | $[M - H_3]^{3-}$ |
| 982.6 | 982.6 | 14 | 2 | 2 | 0 | $[M - H_2 + H_2PO_4]^{3-}$ |
| 1016.6 | 1016.6 | 14 | 2 | 3 | 1 | $[M - H_2 + Na + H_2PO_4]^{3-}$ |
| 1030.6 | 1030.6 | 15 | 2 | 3 | 0 | $[M - H_3]^{3-}$ |
| 1036.7 | 1036.6 | 15 | 2 | 2 | 0 | $[M - H_2 + H_2PO_4]^{3-}$ |
| 1070.7 | 1070.6 | 15 | 2 | 3 | 1 | $[M - H_2 + Na + H_2PO_4]^{3-}$ |
| 1084.7 | 1084.6 | 16 | 2 | 3 | 0 | $[M - H_3]^{3-}$ |
| 1090.7 | 1090.6 | 16 | 2 | 2 | 0 | $[M - H_2 + H_2PO_4]^{3-}$ |
| 1124.7 | 1124.6 | 16 | 2 | 3 | 1 | $[M - H_2 + Na + H_2PO_4]^{3-}$ |
| 1138.7 | 1138.6 | 17 | 2 | 3 | 0 | $[M - H_3]^{3-}$ |
| 1144.7 | 1144.7 | 17 | 2 | 2 | 0 | $[M - H_2 + H_2PO_4]^{3-}$ |
| 1178.7 | 1178.6 | 17 | 2 | 3 | 1 | $[M - H_2 + Na + H_2PO_4]^{3-}$ |
| 1192.7 | 1192.7 | 18 | 2 | 3 | 0 | $[M - H_3]^{3-}$ |
| 1198.7 | 1198.7 | 18 | 2 | 2 | 0 | $[M - H_2 + H_2PO_4]^{3-}$ |
| 1246.7 | 1246.7 | 19 | 2 | 3 | 0 | $[M - H_3]^{3-}$ |
| 1252.7 | 1252.7 | 19 | 2 | 2 | 0 | $[M - H_2 + H_2PO_4]^{3-}$ |
| 1300.7 | 1300.7 | 20 | 2 | 3 | 0 | $[M - H_3]^{3-}$ |
| 1354.8 | 1354.7 | 21 | 2 | 3 | 0 | $[M - H_3]^{3-}$ |
| 1409.1 | 1408.7 | 22 | 2 | 3 | 0 | $[M - H_3]^{3-}$ |

**Supplementary Table. 4**: Abdala N-glycan compositions from ion mobility extracted triply charged ions with the identified molecular ion.

| Mannose | GlcNAc | Phosphate |
|---|---|---|
| 1 | 2 | 1 |
| 2 | 2 | 1 |
| 3 | 2 | 1 |
| 4 | 2 | 1 |
| 5 | 2 | 1 |
| 6 | 2 | 1 |
| 7 | 2 | 1 |
| 8 | 2 | 1 |
| 9 | 2 | 1 |
| | | 2 |
| 10 | 2 | 1 |
| | | 2 |
| | | 3 |
| 11 | 2 | 1 |
| | | 2 |
| | | 3 |
| 12 | 2 | 1 |
| | | 2 |
| | | 3 |
| 13 | 2 | 1 |
| | | 2 |
| | | 3 |
| 14 | 2 | 1 |
| | | 2 |
| | | 3 |
| 15 | 2 | 2 |
| | | 3 |
| 16 | 2 | 2 |
| | | 3 |
| 17 | 2 | 2 |
| | | 3 |
| 18 | 2 | 2 |
| | | 3 |
| 19 | 2 | 2 |
| | | 3 |
| 20 | 2 | 2 |
| | | 3 |
| 21 | 2 | 2 |
| | | 3 |
| 22 | 2 | 3 |

**Supplementary Table. 5**: Summary of the major Abdala N-glycans identified by IM-MS.

| Dataset | HEK-derived RBD | Abdala |
|---|---|---|
| HDX reaction details | deuterated phosphate buffer saline (pH/Dread=7.3, $D_2O$ fraction=0.95) | |
| HDX time courses | 10 s (ice), 10 s (23 °C), 100 s (23 °C), 1,000 s (23 °C), 10,000 s (23 °C), 9 h (28 °C) | |
| HDX control sample | maximally labelled control (quadruplicates) | |
| Back-exchange (Mean/IQR) | 32.7% / 15.6% | |
| # peptides/ average peptide length | 81 / 13.95 | 83 / 14.33 |
| Sequence coverage | 82.30% | |
| Replicates (technical) | triplicates: 10 s (ice), 9 h (28 °C); duplicates: all other time points | |
| Repeatability (average STD) | 0.047095842 | 0.045653481 |
| Significant difference in HDX ($\Delta$HDX>X D) | 0.44 (99% CI) | |

**Supplementary Table. 6**: Summary of HDX experimental workflow, reaction details and time course information.

**Supplementary Table 7**. Individual identified peptide and their respective deuterium uptake at all time points.

| Start | End | Sequence | Glycan | Max Uptake | MHP | State | Exposure | Uptake | Uptake SD | RT | RT SD | Back exchange |
|---|---|---|---|---|---|---|---|---|---|---|---|---|
| 348 | 361 | ASVYAWNRKRISNC | | 13 | 1667.8435 | RBD HEK 293-F | 10 s (ice) | 1.955344 | 0.014694 | 2.876962 | 0.000868 | |
| 348 | 361 | ASVYAWNRKRISNC | | 13 | 1667.8435 | RBD HEK 293-F | 10 s (23 °C) | 2.928052 | 0.0003 | 2.869374 | 0.00325 | |
| 348 | 361 | ASVYAWNRKRISNC | | 13 | 1667.8435 | RBD HEK 293-F | 100 s (23 °C) | 3.408539 | 0.025032 | 2.870422 | 0.000675 | |
| 348 | 361 | ASVYAWNRKRISNC | | 13 | 1667.8435 | RBD HEK 293-F | 1,000 s (23 °C) | 4.455666 | 0.070636 | 2.871059 | 0.000999 | |
| 348 | 361 | ASVYAWNRKRISNC | | 13 | 1667.8435 | RBD HEK 293-F | 10,000 s (23 °C) | 4.880255 | 0.01894 | 2.874118 | 0.001508 | |
| 348 | 361 | ASVYAWNRKRISNC | | 13 | 1667.8435 | RBD HEK 293-F | 9 h (28 °C) | 5.748446 | 0.012495 | 2.869342 | 0.002087 | |
| 348 | 361 | ASVYAWNRKRISNC | | 13 | 1667.8435 | RBD Abdala | 10 s (ice) | 2.159235 | 0.062518 | 2.876959 | 0.00017 | |
| 348 | 361 | ASVYAWNRKRISNC | | 13 | 1667.8435 | RBD Abdala | 10 s (23 °C) | 2.992945 | 0.084159 | 2.870622 | 0.002075 | |
| 348 | 361 | ASVYAWNRKRISNC | | 13 | 1667.8435 | RBD Abdala | 100 s (23 °C) | 3.584116 | 0.065636 | 2.874172 | 0.001844 | |
| 348 | 361 | ASVYAWNRKRISNC | | 13 | 1667.8435 | RBD Abdala | 1,000 s (23 °C) | 4.423155 | 0.029391 | 2.872372 | 0.001531 | |
| 348 | 361 | ASVYAWNRKRISNC | | 13 | 1667.8435 | RBD Abdala | 10,000 s (23 °C) | 5.094983 | 0.01089 | 2.868221 | 0.000259 | |
| 348 | 361 | ASVYAWNRKRISNC | | 13 | 1667.8435 | RBD Abdala | 9 h (28 °C) | 6.946338 | 0.120869 | 2.868227 | 0.002517 | |
| 348 | 361 | ASVYAWNRKRISNC | | 13 | 1667.8435 | MaxD | Max | 9.304731 | 0.116122 | 2.865201 | 0.002122 | 0.246580486 |
| 351 | 361 | YAWNRKRISNC | | 10 | 1410.7059 | RBD HEK 293-F | 10 s (ice) | 1.655598 | 0.017105 | 2.436898 | 0.001807 | |
| 351 | 361 | YAWNRKRISNC | | 10 | 1410.7059 | RBD HEK 293-F | 10 s (23 °C) | 2.557193 | 0.015671 | 2.430521 | 0.00242 | |
| 351 | 361 | YAWNRKRISNC | | 10 | 1410.7059 | RBD HEK 293-F | 100 s (23 °C) | 3.047359 | 0.001151 | 2.431602 | 0.000893 | |
| 351 | 361 | YAWNRKRISNC | | 10 | 1410.7059 | RBD HEK 293-F | 1,000 s (23 °C) | 3.951641 | 0.018384 | 2.430131 | 0.006899 | |
| 351 | 361 | YAWNRKRISNC | | 10 | 1410.7059 | RBD HEK 293-F | 10,000 s (23 °C) | 4.350997 | 0.042902 | 2.434139 | 0.000211 | |
| 351 | 361 | YAWNRKRISNC | | 10 | 1410.7059 | RBD HEK 293-F | 9 h (28 °C) | 4.976075 | 0.069232 | 2.43178 | 0.001179 | |
| 351 | 361 | YAWNRKRISNC | | 10 | 1410.7059 | RBD Abdala | 10 s (ice) | 1.860755 | 0.031988 | 2.4377 | 0.003353 | |
| 351 | 361 | YAWNRKRISNC | | 10 | 1410.7059 | RBD Abdala | 10 s (23 °C) | 2.614399 | 0.113444 | 2.428969 | 0.004473 | |
| 351 | 361 | YAWNRKRISNC | | 10 | 1410.7059 | RBD Abdala | 100 s (23 °C) | 3.169938 | 0.046153 | 2.42807 | 0.006892 | |
| 351 | 361 | YAWNRKRISNC | | 10 | 1410.7059 | RBD Abdala | 1,000 s (23 °C) | 3.984625 | 0.009823 | 2.424236 | 0.0032 | |

| | | | | | | | | | | | | |
|---|---|---|---|---|---|---|---|---|---|---|---|---|
| 351 | 361 | YAWNRKRISNC | | 10 | 1410.7059 | RBD Abdala | 10,000 s (23 °C) | 4.55968 | 0.034245 | 2.429386 | 0.002971 | |
| 351 | 361 | YAWNRKRISNC | | 10 | 1410.7059 | RBD Abdala | 9 h (28 °C) | 5.54967 | 0.117952 | 2.427783 | 0.004683 | |
| 351 | 361 | YAWNRKRISNC | | 10 | 1410.7059 | MaxD | Max | 6.345327 | 0.032506 | 2.410959 | 0.008421 | 0.332070842 |
| 362 | 368 | VADYSVL | | 6 | 766.3981 | RBD HEK 293-F | 10 s (ice) | 0.468679 | 0.047573 | 3.912808 | 0.016536 | |
| 362 | 368 | VADYSVL | | 6 | 766.3981 | RBD HEK 293-F | 10 s (23 °C) | 2.101795 | 0.027184 | 3.919696 | 0.022544 | |
| 362 | 368 | VADYSVL | | 6 | 766.3981 | RBD HEK 293-F | 100 s (23 °C) | 3.270561 | 0.080521 | 3.919694 | 0.026797 | |
| 362 | 368 | VADYSVL | | 6 | 766.3981 | RBD HEK 293-F | 1,000 s (23 °C) | 3.767914 | 0.065584 | 3.914728 | 0.022786 | |
| 362 | 368 | VADYSVL | | 6 | 766.3981 | RBD HEK 293-F | 10,000 s (23 °C) | 3.804384 | 0.07617 | 3.915098 | 0.019058 | |
| 362 | 368 | VADYSVL | | 6 | 766.3981 | RBD HEK 293-F | 9 h (28 °C) | 3.825025 | 0.005961 | 3.907857 | 0.003601 | |
| 362 | 368 | VADYSVL | | 6 | 766.3981 | RBD Abdala | 10 s (ice) | 0.50067 | 0.055368 | 3.91979 | 0.016313 | |
| 362 | 368 | VADYSVL | | 6 | 766.3981 | RBD Abdala | 10 s (23 °C) | 2.084354 | 0.080724 | 3.943223 | 0.002378 | |
| 362 | 368 | VADYSVL | | 6 | 766.3981 | RBD Abdala | 100 s (23 °C) | 3.249689 | 0.107339 | 3.949947 | 0.002906 | |
| 362 | 368 | VADYSVL | | 6 | 766.3981 | RBD Abdala | 1,000 s (23 °C) | 3.871382 | 0.040688 | 3.94159 | 0.003539 | |
| 362 | 368 | VADYSVL | | 6 | 766.3981 | RBD Abdala | 10,000 s (23 °C) | 3.839054 | 0.009237 | 3.916128 | 0.016491 | |
| 362 | 368 | VADYSVL | | 6 | 766.3981 | RBD Abdala | 9 h (28 °C) | 3.851313 | 0.052443 | 3.933513 | 0.025109 | |
| 362 | 368 | VADYSVL | | 6 | 766.3981 | MaxD | Max | 3.909954 | 0.025003 | 3.906379 | 0.003663 | 0.314043158 |
| 369 | 374 | YNSASF | | 5 | 688.2937 | RBD HEK 293-F | 10 s (ice) | 2.598458 | 0.041156 | 2.850612 | 0.002479 | |
| 369 | 374 | YNSASF | | 5 | 688.2937 | RBD HEK 293-F | 10 s (23 °C) | 3.156226 | 0.005479 | 2.844405 | 0.003921 | |
| 369 | 374 | YNSASF | | 5 | 688.2937 | RBD HEK 293-F | 100 s (23 °C) | 3.196583 | 0.012978 | 2.84868 | 0.003141 | |
| 369 | 374 | YNSASF | | 5 | 688.2937 | RBD HEK 293-F | 1,000 s (23 °C) | 3.185078 | 0.024795 | 2.847533 | 0.004171 | |
| 369 | 374 | YNSASF | | 5 | 688.2937 | RBD HEK 293-F | 10,000 s (23 °C) | 3.202294 | 0.012513 | 2.855681 | 0.000975 | |
| 369 | 374 | YNSASF | | 5 | 688.2937 | RBD HEK 293-F | 9 h (28 °C) | 3.141404 | 0.003294 | 2.850508 | 0.00261 | |
| 369 | 374 | YNSASF | | 5 | 688.2937 | RBD Abdala | 10 s (ice) | 2.56986 | 0.036324 | 2.847898 | 0.002515 | |
| 369 | 374 | YNSASF | | 5 | 688.2937 | RBD Abdala | 10 s (23 °C) | 3.128337 | 0.033479 | 2.842593 | 0.00213 | |
| 369 | 374 | YNSASF | | 5 | 688.2937 | RBD Abdala | 100 s (23 °C) | 3.029571 | 0.117359 | 2.844656 | 0.007473 | |
| 369 | 374 | YNSASF | | 5 | 688.2937 | RBD Abdala | 1,000 s (23 °C) | 3.098338 | 0.064842 | 2.841392 | 0.003911 | |

| | | | | | | | | | | | | |
|---|---|---|---|---|---|---|---|---|---|---|---|---|
| 369 | 374 | YNSASF | | 5 | 688.2937 | RBD Abdala | 10,000 s (23 °C) | 3.180549 | 0.049221 | 2.842 | 0.003527 | |
| 369 | 374 | YNSASF | | 5 | 688.2937 | RBD Abdala | 9 h (28 °C) | 3.189511 | 0.025291 | 2.846722 | 0.005201 | |
| 369 | 374 | YNSASF | | 5 | 688.2937 | MaxD | Max | 3.132393 | 0.052762 | 2.844649 | 0.005754 | 0.340548842 |
| 369 | 376 | YNSASFST | | 7 | 876.3734 | RBD HEK 293-F | 10 s (ice) | 3.727628 | 0.056568 | 2.589058 | 0.004288 | |
| 369 | 376 | YNSASFST | | 7 | 876.3734 | RBD HEK 293-F | 10 s (23 °C) | 4.729309 | 0.003029 | 2.584708 | 0.003935 | |
| 369 | 376 | YNSASFST | | 7 | 876.3734 | RBD HEK 293-F | 100 s (23 °C) | 5.27503 | 0.01946 | 2.588203 | 0.000217 | |
| 369 | 376 | YNSASFST | | 7 | 876.3734 | RBD HEK 293-F | 1,000 s (23 °C) | 5.293316 | 0.023177 | 2.585377 | 0.004728 | |
| 369 | 376 | YNSASFST | | 7 | 876.3734 | RBD HEK 293-F | 10,000 s (23 °C) | 5.318597 | 0.013052 | 2.592184 | 0.001928 | |
| 369 | 376 | YNSASFST | | 7 | 876.3734 | RBD HEK 293-F | 9 h (28 °C) | 5.279288 | 0.014271 | 2.586734 | 0.002218 | |
| 369 | 376 | YNSASFST | | 7 | 876.3734 | RBD Abdala | 10 s (ice) | 3.677814 | 0.044109 | 2.587549 | 0.002169 | |
| 369 | 376 | YNSASFST | | 7 | 876.3734 | RBD Abdala | 10 s (23 °C) | 4.719846 | 0.0319 | 2.582982 | 0.003941 | |
| 369 | 376 | YNSASFST | | 7 | 876.3734 | RBD Abdala | 100 s (23 °C) | 5.179134 | 0.128509 | 2.584346 | 0.006214 | |
| 369 | 376 | YNSASFST | | 7 | 876.3734 | RBD Abdala | 1,000 s (23 °C) | 5.289368 | 0.04503 | 2.581874 | 0.003557 | |
| 369 | 376 | YNSASFST | | 7 | 876.3734 | RBD Abdala | 10,000 s (23 °C) | 5.286599 | 0.003972 | 2.584029 | 0.003747 | |
| 369 | 376 | YNSASFST | | 7 | 876.3734 | RBD Abdala | 9 h (28 °C) | 5.322424 | 0.040836 | 2.583996 | 0.005932 | |
| 369 | 376 | YNSASFST | | 7 | 876.3734 | MaxD | Max | 5.193956 | 0.002373 | 2.589304 | 0.002416 | 0.218953985 |
| 374 | 387 | FSTFKCYGVSPTKL | | 12 | 1577.8032 | RBD HEK 293-F | 10 s (ice) | 3.905532 | 0.072309 | 4.359804 | 0.017696 | |
| 374 | 387 | FSTFKCYGVSPTKL | | 12 | 1577.8032 | RBD HEK 293-F | 10 s (23 °C) | 5.236888 | 0.036381 | 4.348968 | 0.023948 | |
| 374 | 387 | FSTFKCYGVSPTKL | | 12 | 1577.8032 | RBD HEK 293-F | 100 s (23 °C) | 6.627466 | 0.067538 | 4.359494 | 0.026149 | |
| 374 | 387 | FSTFKCYGVSPTKL | | 12 | 1577.8032 | RBD HEK 293-F | 1,000 s (23 °C) | 7.140544 | 0.022749 | 4.358641 | 0.028775 | |
| 374 | 387 | FSTFKCYGVSPTKL | | 12 | 1577.8032 | RBD HEK 293-F | 10,000 s (23 °C) | 7.218034 | 0.011342 | 4.357843 | 0.016498 | |
| 374 | 387 | FSTFKCYGVSPTKL | | 12 | 1577.8032 | RBD HEK 293-F | 9 h (28 °C) | 7.763164 | 0.041202 | 4.344195 | 0.003452 | |
| 374 | 387 | FSTFKCYGVSPTKL | | 12 | 1577.8032 | RBD Abdala | 10 s (ice) | 3.938465 | 0.061296 | 4.361613 | 0.018367 | |
| 374 | 387 | FSTFKCYGVSPTKL | | 12 | 1577.8032 | RBD Abdala | 10 s (23 °C) | 5.354001 | 0.024453 | 4.38241 | 0.004378 | |
| 374 | 387 | FSTFKCYGVSPTKL | | 12 | 1577.8032 | RBD Abdala | 100 s (23 °C) | 6.545096 | 0.114759 | 4.394435 | 0.002866 | |
| 374 | 387 | FSTFKCYGVSPTKL | | 12 | 1577.8032 | RBD Abdala | 1,000 s (23 °C) | 7.112187 | 0.028867 | 4.384888 | 0.00572 | |

| | | | | | | | | | | | | |
|---|---|---|---|---|---|---|---|---|---|---|---|---|
| 374 | 387 | FSTFKCYGVSPTKL | | 12 | 1577.8032 | RBD Abdala | 10,000 s (23 °C) | 7.275354 | 0.021935 | 4.364466 | 0.016453 | |
| 374 | 387 | FSTFKCYGVSPTKL | | 12 | 1577.8032 | RBD Abdala | 9 h (28 °C) | 7.847006 | 0.078378 | 4.372009 | 0.026917 | |
| 374 | 387 | FSTFKCYGVSPTKL | | 12 | 1577.8032 | MaxD | Max | 8.303963 | 0.033058 | 4.346688 | 0.002948 | 0.271582193 |
| 375 | 387 | STFKCYGVSPTKL | | 11 | 1430.7348 | RBD HEK 293-F | 10 s (ice) | 3.984443 | 0.067336 | 3.726677 | 0.020115 | |
| 375 | 387 | STFKCYGVSPTKL | | 11 | 1430.7348 | RBD HEK 293-F | 10 s (23 °C) | 5.099582 | 0.026487 | 3.730495 | 0.029257 | |
| 375 | 387 | STFKCYGVSPTKL | | 11 | 1430.7348 | RBD HEK 293-F | 100 s (23 °C) | 6.122784 | 0.071481 | 3.735519 | 0.037965 | |
| 375 | 387 | STFKCYGVSPTKL | | 11 | 1430.7348 | RBD HEK 293-F | 1,000 s (23 °C) | 6.648377 | 0.039508 | 3.730019 | 0.034644 | |
| 375 | 387 | STFKCYGVSPTKL | | 11 | 1430.7348 | RBD HEK 293-F | 10,000 s (23 °C) | 6.819142 | 0.009174 | 3.734353 | 0.023232 | |
| 375 | 387 | STFKCYGVSPTKL | | 11 | 1430.7348 | RBD HEK 293-F | 9 h (28 °C) | 7.394933 | 0.050473 | 3.717139 | 0.005616 | |
| 375 | 387 | STFKCYGVSPTKL | | 11 | 1430.7348 | RBD Abdala | 10 s (ice) | 3.967924 | 0.075141 | 3.735215 | 0.020324 | |
| 375 | 387 | STFKCYGVSPTKL | | 11 | 1430.7348 | RBD Abdala | 10 s (23 °C) | 5.079892 | 0.063158 | 3.766812 | 0.004787 | |
| 375 | 387 | STFKCYGVSPTKL | | 11 | 1430.7348 | RBD Abdala | 100 s (23 °C) | 5.924687 | 0.212903 | 3.770962 | 0.005066 | |
| 375 | 387 | STFKCYGVSPTKL | | 11 | 1430.7348 | RBD Abdala | 1,000 s (23 °C) | 6.679963 | 0.005493 | 3.764575 | 0.007774 | |
| 375 | 387 | STFKCYGVSPTKL | | 11 | 1430.7348 | RBD Abdala | 10,000 s (23 °C) | 6.836456 | 0.038482 | 3.735786 | 0.019261 | |
| 375 | 387 | STFKCYGVSPTKL | | 11 | 1430.7348 | RBD Abdala | 9 h (28 °C) | 7.467527 | 0.00782 | 3.74527 | 0.031889 | |
| 375 | 387 | STFKCYGVSPTKL | | 11 | 1430.7348 | MaxD | Max | 8.106143 | 0.049475 | 3.712605 | 0.003741 | 0.224292536 |
| 378 | 387 | KCYGVSPTKL | | 8 | 1095.5867 | RBD HEK 293-F | 10 s (ice) | 2.895588 | 0.056125 | 2.595128 | 0.000559 | |
| 378 | 387 | KCYGVSPTKL | | 8 | 1095.5867 | RBD HEK 293-F | 10 s (23 °C) | 3.577862 | 0.006661 | 2.587439 | 0.001132 | |
| 378 | 387 | KCYGVSPTKL | | 8 | 1095.5867 | RBD HEK 293-F | 100 s (23 °C) | 4.515872 | 0.081657 | 2.586981 | 0.002111 | |
| 378 | 387 | KCYGVSPTKL | | 8 | 1095.5867 | RBD HEK 293-F | 1,000 s (23 °C) | 5.022153 | 0.038402 | 2.587222 | 0.0051 | |
| 378 | 387 | KCYGVSPTKL | | 8 | 1095.5867 | RBD HEK 293-F | 10,000 s (23 °C) | 4.982661 | 0.052111 | 2.592708 | 0.001982 | |
| 378 | 387 | KCYGVSPTKL | | 8 | 1095.5867 | RBD HEK 293-F | 9 h (28 °C) | 5.1975 | 0.050533 | 2.591914 | 0.001817 | |
| 378 | 387 | KCYGVSPTKL | | 8 | 1095.5867 | RBD Abdala | 10 s (ice) | 2.95597 | 0.02427 | 2.593359 | 0.002627 | |
| 378 | 387 | KCYGVSPTKL | | 8 | 1095.5867 | RBD Abdala | 10 s (23 °C) | 3.672471 | 0.000608 | 2.586953 | 0.002609 | |
| 378 | 387 | KCYGVSPTKL | | 8 | 1095.5867 | RBD Abdala | 100 s (23 °C) | 4.367399 | 0.065796 | 2.588861 | 0.005357 | |
| 378 | 387 | KCYGVSPTKL | | 8 | 1095.5867 | RBD Abdala | 1,000 s (23 °C) | 4.945594 | 0.055854 | 2.586182 | 0.002705 | |

| | | | | | | | | | | | | |
|---|---|---|---|---|---|---|---|---|---|---|---|---|
| 378 | 387 | KCYGVSPTKL | | 8 | 1095.5867 | RBD Abdala | 10,000 s (23 °C) | 5.030441 | 0.035404 | 2.58779 | 0.002084 | |
| 378 | 387 | KCYGVSPTKL | | 8 | 1095.5867 | RBD Abdala | 9 h (28 °C) | 5.213894 | 0.025731 | 2.589025 | 0.004257 | |
| 378 | 387 | KCYGVSPTKL | | 8 | 1095.5867 | MaxD | Max | 5.782683 | 0.033234 | 2.578675 | 0.005799 | 0.239120658 |
| 388 | 392 | NDLCF | | 4 | 611.2494 | RBD HEK 293-F | 10 s (ice) | 0.495635 | 0.026007 | 4.562098 | 0.009868 | |
| 388 | 392 | NDLCF | | 4 | 611.2494 | RBD HEK 293-F | 10 s (23 °C) | 1.12286 | 0.010829 | 4.552414 | 0.021588 | |
| 388 | 392 | NDLCF | | 4 | 611.2494 | RBD HEK 293-F | 100 s (23 °C) | 1.479233 | 0.004004 | 4.561122 | 0.023963 | |
| 388 | 392 | NDLCF | | 4 | 611.2494 | RBD HEK 293-F | 1,000 s (23 °C) | 2.042021 | 0.015038 | 4.561582 | 0.02297 | |
| 388 | 392 | NDLCF | | 4 | 611.2494 | RBD HEK 293-F | 10,000 s (23 °C) | 2.122263 | 0.012625 | 4.55967 | 0.012495 | |
| 388 | 392 | NDLCF | | 4 | 611.2494 | RBD HEK 293-F | 9 h (28 °C) | 2.092668 | 0.022312 | 4.551996 | 0.004071 | |
| 388 | 392 | NDLCF | | 4 | 611.2494 | RBD Abdala | 10 s (ice) | 0.579687 | 0.008098 | 4.563807 | 0.011373 | |
| 388 | 392 | NDLCF | | 4 | 611.2494 | RBD Abdala | 10 s (23 °C) | 1.186744 | 0.015143 | 4.577992 | 0.003756 | |
| 388 | 392 | NDLCF | | 4 | 611.2494 | RBD Abdala | 100 s (23 °C) | 1.465922 | 0.053143 | 4.586792 | 0.003135 | |
| 388 | 392 | NDLCF | | 4 | 611.2494 | RBD Abdala | 1,000 s (23 °C) | 2.030122 | 0.031392 | 4.579807 | 0.004664 | |
| 388 | 392 | NDLCF | | 4 | 611.2494 | RBD Abdala | 10,000 s (23 °C) | 2.117236 | 0.00438 | 4.567212 | 0.01131 | |
| 388 | 392 | NDLCF | | 4 | 611.2494 | RBD Abdala | 9 h (28 °C) | 2.074448 | 0.016649 | 4.572075 | 0.018775 | |
| 388 | 392 | NDLCF | | 4 | 611.2494 | MaxD | Max | 2.102064 | 0.026234 | 4.55279 | 0.001727 | 0.446825263 |
| 390 | 395 | LCFTNV | | 5 | 696.3385 | RBD HEK 293-F | 10 s (ice) | 0.478917 | 0.017807 | 4.358682 | 0.016789 | |
| 390 | 395 | LCFTNV | | 5 | 696.3385 | RBD HEK 293-F | 10 s (23 °C) | 1.514798 | 0.026342 | 4.353193 | 0.022143 | |
| 390 | 395 | LCFTNV | | 5 | 696.3385 | RBD HEK 293-F | 100 s (23 °C) | 2.49526 | 0.013125 | 4.358511 | 0.027143 | |
| 390 | 395 | LCFTNV | | 5 | 696.3385 | RBD HEK 293-F | 1,000 s (23 °C) | 2.999602 | 0.052307 | 4.357428 | 0.029001 | |
| 390 | 395 | LCFTNV | | 5 | 696.3385 | RBD HEK 293-F | 10,000 s (23 °C) | 3.25653 | 0.056535 | 4.354627 | 0.017748 | |
| 390 | 395 | LCFTNV | | 5 | 696.3385 | RBD HEK 293-F | 9 h (28 °C) | 3.216235 | 0.019251 | 4.341868 | 0.006255 | |
| 390 | 395 | LCFTNV | | 5 | 696.3385 | RBD Abdala | 10 s (ice) | 0.652333 | 0.022771 | 4.36005 | 0.017501 | |
| 390 | 395 | LCFTNV | | 5 | 696.3385 | RBD Abdala | 10 s (23 °C) | 1.708314 | 0.014241 | 4.381644 | 0.004557 | |
| 390 | 395 | LCFTNV | | 5 | 696.3385 | RBD Abdala | 100 s (23 °C) | 2.532744 | 0.074076 | 4.388281 | 0.001278 | |
| 390 | 395 | LCFTNV | | 5 | 696.3385 | RBD Abdala | 1,000 s (23 °C) | 3.102612 | 0.049845 | 4.379482 | 0.004535 | |

| | | | | | | | | | | | | |
|---|---|---|---|---|---|---|---|---|---|---|---|---|
| 390 | 395 | LCFTNV | | 5 | 696.3385 | RBD Abdala | 10,000 s (23 °C) | 3.377593 | 0.052742 | 4.361903 | 0.014487 | |
| 390 | 395 | LCFTNV | | 5 | 696.3385 | RBD Abdala | 9 h (28 °C) | 3.257047 | 0.007947 | 4.368611 | 0.023931 | |
| 390 | 395 | LCFTNV | | 5 | 696.3385 | MaxD | Max | 3.250281 | 0.012199 | 4.34128 | 0.003284 | 0.315730316 |
| 392 | 399 | FTNVYADS | | 7 | 916.4047 | RBD HEK 293-F | 10 s (ice) | 0.103604 | 0.048423 | 2.829007 | 0.002895 | |
| 392 | 399 | FTNVYADS | | 7 | 916.4047 | RBD HEK 293-F | 10 s (23 °C) | 0.229664 | 0.015612 | 2.827636 | 0.002232 | |
| 392 | 399 | FTNVYADS | | 7 | 916.4047 | RBD HEK 293-F | 100 s (23 °C) | 0.613731 | 0.006072 | 2.829914 | 0.000187 | |
| 392 | 399 | FTNVYADS | | 7 | 916.4047 | RBD HEK 293-F | 1,000 s (23 °C) | 1.175617 | 0.011907 | 2.826289 | 0.003783 | |
| 392 | 399 | FTNVYADS | | 7 | 916.4047 | RBD HEK 293-F | 10,000 s (23 °C) | 1.675755 | 0.009404 | 2.830308 | 0.00086 | |
| 392 | 399 | FTNVYADS | | 7 | 916.4047 | RBD HEK 293-F | 9 h (28 °C) | 2.174512 | 0.022088 | 2.825205 | 0.00137 | |
| 392 | 399 | FTNVYADS | | 7 | 916.4047 | RBD Abdala | 10 s (ice) | 0.08561 | 0.01735 | 2.829923 | 0.003129 | |
| 392 | 399 | FTNVYADS | | 7 | 916.4047 | RBD Abdala | 10 s (23 °C) | 0.263706 | 0.039415 | 2.824712 | 0.003086 | |
| 392 | 399 | FTNVYADS | | 7 | 916.4047 | RBD Abdala | 100 s (23 °C) | 0.67956 | 0.003285 | 2.82345 | 0.00508 | |
| 392 | 399 | FTNVYADS | | 7 | 916.4047 | RBD Abdala | 1,000 s (23 °C) | 1.349414 | 0.022401 | 2.823178 | 0.002469 | |
| 392 | 399 | FTNVYADS | | 7 | 916.4047 | RBD Abdala | 10,000 s (23 °C) | 1.676198 | 0.02647 | 2.823542 | 0.002569 | |
| 392 | 399 | FTNVYADS | | 7 | 916.4047 | RBD Abdala | 9 h (28 °C) | 2.270209 | 0.016574 | 2.823356 | 0.005521 | |
| 392 | 399 | FTNVYADS | | 7 | 916.4047 | MaxD | Max | 4.69072 | 0.03039 | 2.81977 | 0.003046 | 0.294628571 |
| 392 | 400 | FTNVYADSF | | 8 | 1063.4731 | RBD HEK 293-F | 10 s (ice) | 0.095953 | 0.01338 | 4.741066 | 0.006189 | |
| 392 | 400 | FTNVYADSF | | 8 | 1063.4731 | RBD HEK 293-F | 10 s (23 °C) | 0.298877 | 0.013381 | 4.739235 | 0.017398 | |
| 392 | 400 | FTNVYADSF | | 8 | 1063.4731 | RBD HEK 293-F | 100 s (23 °C) | 0.545277 | 0.03741 | 4.749099 | 0.022024 | |
| 392 | 400 | FTNVYADSF | | 8 | 1063.4731 | RBD HEK 293-F | 1,000 s (23 °C) | 0.972575 | 0.049051 | 4.741118 | 0.016518 | |
| 392 | 400 | FTNVYADSF | | 8 | 1063.4731 | RBD HEK 293-F | 10,000 s (23 °C) | 1.583096 | 0.018169 | 4.746289 | 0.012782 | |
| 392 | 400 | FTNVYADSF | | 8 | 1063.4731 | RBD HEK 293-F | 9 h (28 °C) | 1.989306 | 0.017525 | 4.739132 | 0.00452 | |
| 392 | 400 | FTNVYADSF | | 8 | 1063.4731 | RBD Abdala | 10 s (ice) | 0.077009 | 0.017958 | 4.750057 | 0.008525 | |
| 392 | 400 | FTNVYADSF | | 8 | 1063.4731 | RBD Abdala | 10 s (23 °C) | 0.32379 | 0 | 4.756229 | 0 | |
| 392 | 400 | FTNVYADSF | | 8 | 1063.4731 | RBD Abdala | 100 s (23 °C) | 0.618651 | 0.019073 | 4.770667 | 0.000794 | |
| 392 | 400 | FTNVYADSF | | 8 | 1063.4731 | RBD Abdala | 1,000 s (23 °C) | 1.118754 | 0.033421 | 4.763586 | 0.004991 | |

| | | | | | | | | | | | | |
|---|---|---|---|---|---|---|---|---|---|---|---|---|
| 392 | 400 | FTNVYADSF | | 8 | 1063.4731 | RBD Abdala | 10,000 s (23 °C) | 1.60663 | 0.03491 | 4.749341 | 0.005789 | |
| 392 | 400 | FTNVYADSF | | 8 | 1063.4731 | RBD Abdala | 9 h (28 °C) | 2.063415 | 0.045304 | 4.755186 | 0.017356 | |
| 392 | 400 | FTNVYADSF | | 8 | 1063.4731 | MaxD | Max | 5.393107 | 0.038589 | 4.73454 | 0.003655 | 0.290380658 |
| 393 | 399 | TNVYADS | | 6 | 769.3363 | RBD HEK 293-F | 10 s (ice) | 0.020211 | 0.006425 | 1.509445 | 0.032775 | |
| 393 | 399 | TNVYADS | | 6 | 769.3363 | RBD HEK 293-F | 10 s (23 °C) | 0.021949 | 0.002883 | 1.519619 | 0.001786 | |
| 393 | 399 | TNVYADS | | 6 | 769.3363 | RBD HEK 293-F | 100 s (23 °C) | 0.124933 | 0.007034 | 1.525048 | 0.002921 | |
| 393 | 399 | TNVYADS | | 6 | 769.3363 | RBD HEK 293-F | 1,000 s (23 °C) | 0.635269 | 0.025436 | 1.491988 | 0.040077 | |
| 393 | 399 | TNVYADS | | 6 | 769.3363 | RBD HEK 293-F | 10,000 s (23 °C) | 1.101846 | 0.013675 | 1.523547 | 0.000752 | |
| 393 | 399 | TNVYADS | | 6 | 769.3363 | RBD HEK 293-F | 9 h (28 °C) | 1.576224 | 0.029897 | 1.531418 | 0.014429 | |
| 393 | 399 | TNVYADS | | 6 | 769.3363 | RBD Abdala | 10 s (ice) | 0.006811 | 0.010853 | 1.498683 | 0.024415 | |
| 393 | 399 | TNVYADS | | 6 | 769.3363 | RBD Abdala | 10 s (23 °C) | 0.026332 | 0.034604 | 1.489498 | 0.013478 | |
| 393 | 399 | TNVYADS | | 6 | 769.3363 | RBD Abdala | 100 s (23 °C) | 0.1738 | 0.023374 | 1.479063 | 0.05064 | |
| 393 | 399 | TNVYADS | | 6 | 769.3363 | RBD Abdala | 1,000 s (23 °C) | 0.750841 | 0.016877 | 1.433828 | 0.015158 | |
| 393 | 399 | TNVYADS | | 6 | 769.3363 | RBD Abdala | 10,000 s (23 °C) | 1.118424 | 0.001805 | 1.494336 | 0.037655 | |
| 393 | 399 | TNVYADS | | 6 | 769.3363 | RBD Abdala | 9 h (28 °C) | 1.659553 | 0.014773 | 1.491784 | 0.035646 | |
| 393 | 399 | TNVYADS | | 6 | 769.3363 | MaxD | Max | 4.023473 | 0.036536 | 1.495154 | 0.099618 | 0.294127544 |
| 393 | 400 | TNVYADSF | | 7 | 916.4047 | RBD HEK 293-F | 10 s (ice) | 0.054225 | 0.020023 | 3.87379 | 0.017111 | |
| 393 | 400 | TNVYADSF | | 7 | 916.4047 | RBD HEK 293-F | 10 s (23 °C) | 0.056774 | 0.011964 | 3.877387 | 0.027033 | |
| 393 | 400 | TNVYADSF | | 7 | 916.4047 | RBD HEK 293-F | 100 s (23 °C) | 0.139986 | 0.024502 | 3.882765 | 0.029759 | |
| 393 | 400 | TNVYADSF | | 7 | 916.4047 | RBD HEK 293-F | 1,000 s (23 °C) | 0.56856 | 0.034384 | 3.881052 | 0.029357 | |
| 393 | 400 | TNVYADSF | | 7 | 916.4047 | RBD HEK 293-F | 10,000 s (23 °C) | 1.059302 | 0.000126 | 3.883043 | 0.01992 | |
| 393 | 400 | TNVYADSF | | 7 | 916.4047 | RBD HEK 293-F | 9 h (28 °C) | 1.519049 | 0.00993 | 3.868013 | 0.003731 | |
| 393 | 400 | TNVYADSF | | 7 | 916.4047 | RBD Abdala | 10 s (ice) | 0.008417 | 0.045644 | 3.883161 | 0.019144 | |
| 393 | 400 | TNVYADSF | | 7 | 916.4047 | RBD Abdala | 10 s (23 °C) | 0.079938 | 0.002126 | 3.906305 | 0.000361 | |
| 393 | 400 | TNVYADSF | | 7 | 916.4047 | RBD Abdala | 100 s (23 °C) | 0.142886 | 0.014266 | 3.91207 | 0.003072 | |
| 393 | 400 | TNVYADSF | | 7 | 916.4047 | RBD Abdala | 1,000 s (23 °C) | 0.758422 | 0.029368 | 3.907023 | 0.005019 | |

| | | | | | | | | | | | | |
|---|---|---|---|---|---|---|---|---|---|---|---|---|
| 393 | 400 | TNVYADSF | | 7 | 916.4047 | RBD Abdala | 10,000 s (23 °C) | 1.039097 | 0.008169 | 3.885316 | 0.016452 | |
| 393 | 400 | TNVYADSF | | 7 | 916.4047 | RBD Abdala | 9 h (28 °C) | 1.596837 | 0.006752 | 3.891839 | 0.026167 | |
| 393 | 400 | TNVYADSF | | 7 | 916.4047 | MaxD | Max | 4.844359 | 0.043912 | 3.864016 | 0.001028 | 0.271524962 |
| 396 | 400 | YADSF | | 4 | 602.2457 | RBD HEK 293-F | 10 s (ice) | 0.085711 | 0.03206 | 3.129387 | 0.000946 | |
| 396 | 400 | YADSF | | 4 | 602.2457 | RBD HEK 293-F | 10 s (23 °C) | 0.119437 | 0.011762 | 3.123184 | 0.003257 | |
| 396 | 400 | YADSF | | 4 | 602.2457 | RBD HEK 293-F | 100 s (23 °C) | 0.136909 | 0.01093 | 3.127921 | 0.001843 | |
| 396 | 400 | YADSF | | 4 | 602.2457 | RBD HEK 293-F | 1,000 s (23 °C) | 0.137423 | 0.025799 | 3.129964 | 0.00343 | |
| 396 | 400 | YADSF | | 4 | 602.2457 | RBD HEK 293-F | 10,000 s (23 °C) | 0.243716 | 0.007018 | 3.131994 | 3.12E-05 | |
| 396 | 400 | YADSF | | 4 | 602.2457 | RBD HEK 293-F | 9 h (28 °C) | 0.597439 | 0.0208 | 3.127014 | 0.001536 | |
| 396 | 400 | YADSF | | 4 | 602.2457 | RBD Abdala | 10 s (ice) | 0.074619 | 0.029477 | 3.128223 | 0.001802 | |
| 396 | 400 | YADSF | | 4 | 602.2457 | RBD Abdala | 10 s (23 °C) | 0.093704 | 0.015397 | 3.124087 | 0.006915 | |
| 396 | 400 | YADSF | | 4 | 602.2457 | RBD Abdala | 100 s (23 °C) | 0.085649 | 0.004063 | 3.125174 | 0.008574 | |
| 396 | 400 | YADSF | | 4 | 602.2457 | RBD Abdala | 1,000 s (23 °C) | 0.107346 | 0.023143 | 3.124595 | 0.000868 | |
| 396 | 400 | YADSF | | 4 | 602.2457 | RBD Abdala | 10,000 s (23 °C) | 0.221339 | 0.014971 | 3.12627 | 0.004993 | |
| 396 | 400 | YADSF | | 4 | 602.2457 | RBD Abdala | 9 h (28 °C) | 0.661979 | 0.021326 | 3.124415 | 0.004685 | |
| 396 | 400 | YADSF | | 4 | 602.2457 | MaxD | Max | 2.145237 | 0.017988 | 3.127809 | 0.003593 | 0.435463947 |
| 400 | 406 | FVIRGDE | | 6 | 835.4308 | RBD HEK 293-F | 10 s (ice) | 0.148761 | 0.030339 | 2.449269 | 0.004467 | |
| 400 | 406 | FVIRGDE | | 6 | 835.4308 | RBD HEK 293-F | 10 s (23 °C) | 0.246217 | 0.000969 | 2.445627 | 0.001862 | |
| 400 | 406 | FVIRGDE | | 6 | 835.4308 | RBD HEK 293-F | 100 s (23 °C) | 0.374432 | 0.011635 | 2.446659 | 0.001348 | |
| 400 | 406 | FVIRGDE | | 6 | 835.4308 | RBD HEK 293-F | 1,000 s (23 °C) | 0.385186 | 0.009859 | 2.444807 | 0.008503 | |
| 400 | 406 | FVIRGDE | | 6 | 835.4308 | RBD HEK 293-F | 10,000 s (23 °C) | 0.434394 | 0.013445 | 2.45332 | 0.001668 | |
| 400 | 406 | FVIRGDE | | 6 | 835.4308 | RBD HEK 293-F | 9 h (28 °C) | 0.921648 | 0.018216 | 2.451646 | 0.001414 | |
| 400 | 406 | FVIRGDE | | 6 | 835.4308 | RBD Abdala | 10 s (ice) | 0.148738 | 0.035264 | 2.449002 | 0.004594 | |
| 400 | 406 | FVIRGDE | | 6 | 835.4308 | RBD Abdala | 10 s (23 °C) | 0.200051 | 0.0035 | 2.44144 | 0.004276 | |
| 400 | 406 | FVIRGDE | | 6 | 835.4308 | RBD Abdala | 100 s (23 °C) | 0.361904 | 0.012197 | 2.44336 | 0.008595 | |
| 400 | 406 | FVIRGDE | | 6 | 835.4308 | RBD Abdala | 1,000 s (23 °C) | 0.416061 | 0.000161 | 2.437248 | 0.003552 | |

| | | | | | | | | | | | | |
|---|---|---|---|---|---|---|---|---|---|---|---|---|
| 400 | 406 | FVIRGDE | | 6 | 835.4308 | RBD Abdala | 10,000 s (23 °C) | 0.476345 | 0.011038 | 2.444222 | 0.00608 | |
| 400 | 406 | FVIRGDE | | 6 | 835.4308 | RBD Abdala | 9 h (28 °C) | 1.076515 | 0.019647 | 2.445162 | 0.007797 | |
| 400 | 406 | FVIRGDE | | 6 | 835.4308 | MaxD | Max | 3.122901 | 0.035188 | 2.431926 | 0.019159 | 0.452122632 |
| 400 | 421 | FVIRGDEVRQIAPGQTGKIADY | | 20 | 2433.2885 | RBD HEK 293-F | 10 s (ice) | 2.094157 | 0.019897 | 3.773402 | 0.023379 | |
| 400 | 421 | FVIRGDEVRQIAPGQTGKIADY | | 20 | 2433.2885 | RBD HEK 293-F | 10 s (23 °C) | 3.379392 | 0.004593 | 3.771965 | 0.029294 | |
| 400 | 421 | FVIRGDEVRQIAPGQTGKIADY | | 20 | 2433.2885 | RBD HEK 293-F | 100 s (23 °C) | 4.03272 | 0.018328 | 3.785228 | 0.038554 | |
| 400 | 421 | FVIRGDEVRQIAPGQTGKIADY | | 20 | 2433.2885 | RBD HEK 293-F | 1,000 s (23 °C) | 5.210932 | 0.037156 | 3.775451 | 0.039869 | |
| 400 | 421 | FVIRGDEVRQIAPGQTGKIADY | | 20 | 2433.2885 | RBD HEK 293-F | 10,000 s (23 °C) | 6.210799 | 0.026321 | 3.774006 | 0.024124 | |
| 400 | 421 | FVIRGDEVRQIAPGQTGKIADY | | 20 | 2433.2885 | RBD HEK 293-F | 9 h (28 °C) | 8.186135 | 0.07079 | 3.755687 | 0.006945 | |
| 400 | 421 | FVIRGDEVRQIAPGQTGKIADY | | 20 | 2433.2885 | RBD Abdala | 10 s (ice) | 2.093524 | 0.034368 | 3.782074 | 0.024016 | |
| 400 | 421 | FVIRGDEVRQIAPGQTGKIADY | | 20 | 2433.2885 | RBD Abdala | 10 s (23 °C) | 3.352521 | 0.011998 | 3.813407 | 0.004622 | |
| 400 | 421 | FVIRGDEVRQIAPGQTGKIADY | | 20 | 2433.2885 | RBD Abdala | 100 s (23 °C) | 3.879107 | 0.12872 | 3.822948 | 0.003592 | |
| 400 | 421 | FVIRGDEVRQIAPGQTGKIADY | | 20 | 2433.2885 | RBD Abdala | 1,000 s (23 °C) | 5.241901 | 0.003944 | 3.814009 | 0.007623 | |
| 400 | 421 | FVIRGDEVRQIAPGQTGKIADY | | 20 | 2433.2885 | RBD Abdala | 10,000 s (23 °C) | 6.252526 | 0.040541 | 3.779867 | 0.020997 | |
| 400 | 421 | FVIRGDEVRQIAPGQTGKIADY | | 20 | 2433.2885 | RBD Abdala | 9 h (28 °C) | 8.928997 | 0.050878 | 3.789435 | 0.031446 | |
| 400 | 421 | FVIRGDEVRQIAPGQTGKIADY | | 20 | 2433.2885 | MaxD | Max | 13.795442 | 0.077626 | 3.7512 | 0.005436 | 0.273924105 |
| 400 | 422 | FVIRGDEVRQIAPGQTGKIADYN | | 21 | 2547.3314 | RBD HEK 293-F | 10 s (ice) | 2.082513 | 0.02359 | 3.523822 | 0.029492 | |
| 400 | 422 | FVIRGDEVRQIAPGQTGKIADYN | | 21 | 2547.3314 | RBD HEK 293-F | 10 s (23 °C) | 3.363817 | 0.033407 | 3.529079 | 0.038961 | |
| 400 | 422 | FVIRGDEVRQIAPGQTGKIADYN | | 21 | 2547.3314 | RBD HEK 293-F | 100 s (23 °C) | 3.913651 | 0.00396 | 3.537773 | 0.047559 | |
| 400 | 422 | FVIRGDEVRQIAPGQTGKIADYN | | 21 | 2547.3314 | RBD HEK 293-F | 1,000 s (23 °C) | 5.21057 | 0.027267 | 3.529164 | 0.047478 | |
| 400 | 422 | FVIRGDEVRQIAPGQTGKIADYN | | 21 | 2547.3314 | RBD HEK 293-F | 10,000 s (23 °C) | 6.181146 | 0.001513 | 3.522954 | 0.032809 | |
| 400 | 422 | FVIRGDEVRQIAPGQTGKIADYN | | 21 | 2547.3314 | RBD HEK 293-F | 9 h (28 °C) | 8.307179 | 0.042453 | 3.50415 | 0.007375 | |
| 400 | 422 | FVIRGDEVRQIAPGQTGKIADYN | | 21 | 2547.3314 | RBD Abdala | 10 s (ice) | 2.077521 | 0.022098 | 3.529177 | 0.027568 | |
| 400 | 422 | FVIRGDEVRQIAPGQTGKIADYN | | 21 | 2547.3314 | RBD Abdala | 10 s (23 °C) | 3.312624 | 0.000212 | 3.57454 | 0.002604 | |
| 400 | 422 | FVIRGDEVRQIAPGQTGKIADYN | | 21 | 2547.3314 | RBD Abdala | 100 s (23 °C) | 3.853369 | 0.090384 | 3.574582 | 0.002285 | |
| 400 | 422 | FVIRGDEVRQIAPGQTGKIADYN | | 21 | 2547.3314 | RBD Abdala | 1,000 s (23 °C) | 5.233451 | 0.031469 | 3.566781 | 0.005557 | |

| | | | | | | | | | | | | |
|---|---|---|---|---|---|---|---|---|---|---|---|---|
| 400 | 422 | FVIRGDEVRQIAPGQTGKIADYN | | 21 | 2547.3314 | RBD Abdala | 10,000 s (23 °C) | 6.292626 | 0.019946 | 3.533154 | 0.027248 | |
| 400 | 422 | FVIRGDEVRQIAPGQTGKIADYN | | 21 | 2547.3314 | RBD Abdala | 9 h (28 °C) | 9.163418 | 0.073245 | 3.536042 | 0.038792 | |
| 400 | 422 | FVIRGDEVRQIAPGQTGKIADYN | | 21 | 2547.3314 | MaxD | Max | 14.553671 | 0.073405 | 3.499387 | 0.009104 | 0.270492682 |
| 400 | 431 | FVIRGDEVRQIAPGQTGKIADYNYKLPDDFTG | | 29 | 3583.818 | RBD HEK 293-F | 10 s (ice) | 2.493084 | 0.01258 | 4.528917 | 0.007808 | |
| 400 | 431 | FVIRGDEVRQIAPGQTGKIADYNYKLPDDFTG | | 29 | 3583.818 | RBD HEK 293-F | 10 s (23 °C) | 4.065504 | 0.054981 | 4.518447 | 0.014309 | |
| 400 | 431 | FVIRGDEVRQIAPGQTGKIADYNYKLPDDFTG | | 29 | 3583.818 | RBD HEK 293-F | 100 s (23 °C) | 5.420012 | 0.01924 | 4.529816 | 0.016293 | |
| 400 | 431 | FVIRGDEVRQIAPGQTGKIADYNYKLPDDFTG | | 29 | 3583.818 | RBD HEK 293-F | 1,000 s (23 °C) | 6.717104 | 0.053058 | 4.528176 | 0.014911 | |
| 400 | 431 | FVIRGDEVRQIAPGQTGKIADYNYKLPDDFTG | | 29 | 3583.818 | RBD HEK 293-F | 10,000 s (23 °C) | 7.554831 | 0.053411 | 4.526022 | 0.014043 | |
| 400 | 431 | FVIRGDEVRQIAPGQTGKIADYNYKLPDDFTG | | 29 | 3583.818 | RBD HEK 293-F | 9 h (28 °C) | 10.01108 | 0.127816 | 4.520861 | 0.006809 | |
| 400 | 431 | FVIRGDEVRQIAPGQTGKIADYNYKLPDDFTG | | 29 | 3583.818 | RBD Abdala | 10 s (ice) | 2.565184 | 0.005111 | 4.533815 | 0.00971 | |
| 400 | 431 | FVIRGDEVRQIAPGQTGKIADYNYKLPDDFTG | | 29 | 3583.818 | RBD Abdala | 10 s (23 °C) | 4.015579 | 0.03601 | 4.547788 | 0.007373 | |
| 400 | 431 | FVIRGDEVRQIAPGQTGKIADYNYKLPDDFTG | | 29 | 3583.818 | RBD Abdala | 100 s (23 °C) | 5.311108 | 0.099343 | 4.552397 | 0.001033 | |
| 400 | 431 | FVIRGDEVRQIAPGQTGKIADYNYKLPDDFTG | | 29 | 3583.818 | RBD Abdala | 1,000 s (23 °C) | 6.735539 | 0.002549 | 4.555107 | 0.003598 | |
| 400 | 431 | FVIRGDEVRQIAPGQTGKIADYNYKLPDDFTG | | 29 | 3583.818 | RBD Abdala | 10,000 s (23 °C) | 7.752687 | 0.003723 | 4.537711 | 0.010184 | |
| 400 | 431 | FVIRGDEVRQIAPGQTGKIADYNYKLPDDFTG | | 29 | 3583.818 | RBD Abdala | 9 h (28 °C) | 11.078178 | 0.078108 | 4.540302 | 0.015004 | |
| 400 | 431 | FVIRGDEVRQIAPGQTGKIADYNYKLPDDFTG | | 29 | 3583.818 | MaxD | Max | 18.769545 | 0.106915 | 4.515722 | 0.004255 | 0.3187098 |
| 401 | 406 | VIRGDE | | 5 | 688.3624 | RBD HEK 293-F | 10 s (ice) | 0.202294 | 0.021325 | 0.770368 | 0.005275 | |
| 401 | 406 | VIRGDE | | 5 | 688.3624 | RBD HEK 293-F | 10 s (23 °C) | 0.364811 | 0.017828 | 0.770869 | 0.004717 | |
| 401 | 406 | VIRGDE | | 5 | 688.3624 | RBD HEK 293-F | 100 s (23 °C) | 0.590229 | 0.002512 | 0.770965 | 0.003893 | |
| 401 | 406 | VIRGDE | | 5 | 688.3624 | RBD HEK 293-F | 1,000 s (23 °C) | 0.631615 | 0.019376 | 0.770512 | 0.00504 | |
| 401 | 406 | VIRGDE | | 5 | 688.3624 | RBD HEK 293-F | 10,000 s (23 °C) | 0.672082 | 0.000676 | 0.771698 | 0.005086 | |
| 401 | 406 | VIRGDE | | 5 | 688.3624 | RBD HEK 293-F | 9 h (28 °C) | 1.051137 | 0.02242 | 0.773009 | 0.005011 | |
| 401 | 406 | VIRGDE | | 5 | 688.3624 | RBD Abdala | 10 s (ice) | 0.249541 | 0.02411 | 0.769391 | 0.004587 | |
| 401 | 406 | VIRGDE | | 5 | 688.3624 | RBD Abdala | 10 s (23 °C) | 0.405243 | 0.013988 | 0.768605 | 0.004838 | |
| 401 | 406 | VIRGDE | | 5 | 688.3624 | RBD Abdala | 100 s (23 °C) | 0.578337 | 0.041389 | 0.770282 | 0.004426 | |
| 401 | 406 | VIRGDE | | 5 | 688.3624 | RBD Abdala | 1,000 s (23 °C) | 0.6228 | 0.007491 | 0.768572 | 0.004792 | |

| | | | | | | | | | | | | |
|---|---|---|---|---|---|---|---|---|---|---|---|---|
| 401 | 406 | VIRGDE | | 5 | 688.3624 | RBD Abdala | 10,000 s (23 °C) | 0.703778 | 0.02602 | 0.770469 | 0.005355 | |
| 401 | 406 | VIRGDE | | 5 | 688.3624 | RBD Abdala | 9 h (28 °C) | 1.239816 | 0.024283 | 0.770339 | 0.005164 | |
| 401 | 406 | VIRGDE | | 5 | 688.3624 | MaxD | Max | 2.271034 | 0.060995 | 0.771631 | 0.003649 | 0.521887579 |
| 401 | 420 | VIRGDEVRQIAPGQTGKIAD | | 18 | 2123.1567 | RBD HEK 293-F | 10 s (ice) | 2.144281 | 0.044926 | 2.506733 | 0.00197 | |
| 401 | 420 | VIRGDEVRQIAPGQTGKIAD | | 18 | 2123.1567 | RBD HEK 293-F | 10 s (23 °C) | 3.446729 | 0.00424 | 2.500938 | 0.002216 | |
| 401 | 420 | VIRGDEVRQIAPGQTGKIAD | | 18 | 2123.1567 | RBD HEK 293-F | 100 s (23 °C) | 4.17539 | 0.021262 | 2.50237 | 0.000933 | |
| 401 | 420 | VIRGDEVRQIAPGQTGKIAD | | 18 | 2123.1567 | RBD HEK 293-F | 1,000 s (23 °C) | 5.419232 | 0.058525 | 2.500722 | 0.004187 | |
| 401 | 420 | VIRGDEVRQIAPGQTGKIAD | | 18 | 2123.1567 | RBD HEK 293-F | 10,000 s (23 °C) | 6.416592 | 0.01366 | 2.502326 | 0.000362 | |
| 401 | 420 | VIRGDEVRQIAPGQTGKIAD | | 18 | 2123.1567 | RBD HEK 293-F | 9 h (28 °C) | 8.313486 | 0.085019 | 2.495197 | 0.000646 | |
| 401 | 420 | VIRGDEVRQIAPGQTGKIAD | | 18 | 2123.1567 | RBD Abdala | 10 s (ice) | 2.156198 | 0.046681 | 2.506596 | 0.001606 | |
| 401 | 420 | VIRGDEVRQIAPGQTGKIAD | | 18 | 2123.1567 | RBD Abdala | 10 s (23 °C) | 3.398381 | 0.022817 | 2.499875 | 0.002687 | |
| 401 | 420 | VIRGDEVRQIAPGQTGKIAD | | 18 | 2123.1567 | RBD Abdala | 100 s (23 °C) | 4.0536 | 0.099467 | 2.502069 | 0.002356 | |
| 401 | 420 | VIRGDEVRQIAPGQTGKIAD | | 18 | 2123.1567 | RBD Abdala | 1,000 s (23 °C) | 5.459766 | 0.010577 | 2.502726 | 0.000854 | |
| 401 | 420 | VIRGDEVRQIAPGQTGKIAD | | 18 | 2123.1567 | RBD Abdala | 10,000 s (23 °C) | 6.473293 | 0.028057 | 2.497351 | 0.001097 | |
| 401 | 420 | VIRGDEVRQIAPGQTGKIAD | | 18 | 2123.1567 | RBD Abdala | 9 h (28 °C) | 8.955755 | 0.04165 | 2.493665 | 0.002078 | |
| 401 | 420 | VIRGDEVRQIAPGQTGKIAD | | 18 | 2123.1567 | MaxD | Max | 12.227177 | 0.066638 | 2.489871 | 0.001036 | 0.284960409 |
| 401 | 421 | VIRGDEVRQIAPGQTGKIADY | | 19 | 2286.2201 | RBD HEK 293-F | 10 s (ice) | 2.188425 | 0.017274 | 3.099428 | 0.000316 | |
| 401 | 421 | VIRGDEVRQIAPGQTGKIADY | | 19 | 2286.2201 | RBD HEK 293-F | 10 s (23 °C) | 3.493562 | 0.00406 | 3.093311 | 0.004071 | |
| 401 | 421 | VIRGDEVRQIAPGQTGKIADY | | 19 | 2286.2201 | RBD HEK 293-F | 100 s (23 °C) | 4.171158 | 0.044597 | 3.093505 | 0.001178 | |
| 401 | 421 | VIRGDEVRQIAPGQTGKIADY | | 19 | 2286.2201 | RBD HEK 293-F | 1,000 s (23 °C) | 5.422873 | 0.000919 | 3.093333 | 0.0004 | |
| 401 | 421 | VIRGDEVRQIAPGQTGKIADY | | 19 | 2286.2201 | RBD HEK 293-F | 10,000 s (23 °C) | 6.398046 | 0.007712 | 3.093964 | 0.000196 | |
| 401 | 421 | VIRGDEVRQIAPGQTGKIADY | | 19 | 2286.2201 | RBD HEK 293-F | 9 h (28 °C) | 8.369746 | 0.04235 | 3.086225 | 0.002296 | |
| 401 | 421 | VIRGDEVRQIAPGQTGKIADY | | 19 | 2286.2201 | RBD Abdala | 10 s (ice) | 2.191565 | 0.0063 | 3.100946 | 0.001553 | |
| 401 | 421 | VIRGDEVRQIAPGQTGKIADY | | 19 | 2286.2201 | RBD Abdala | 10 s (23 °C) | 3.451284 | 0.008616 | 3.092807 | 0.00186 | |
| 401 | 421 | VIRGDEVRQIAPGQTGKIADY | | 19 | 2286.2201 | RBD Abdala | 100 s (23 °C) | 4.035393 | 0.158767 | 3.096941 | 0.001054 | |
| 401 | 421 | VIRGDEVRQIAPGQTGKIADY | | 19 | 2286.2201 | RBD Abdala | 1,000 s (23 °C) | 5.469581 | 0.005769 | 3.097051 | 0.002526 | |

| | | | | | | | | | | | | |
|---|---|---|---|---|---|---|---|---|---|---|---|---|
| 401 | 421 | VIRGDEVRQIAPGQTGKIADY | | 19 | 2286.2201 | RBD Abdala | 10,000 s (23 °C) | 6.481341 | 0.019937 | 3.09113 | 0.001298 | |
| 401 | 421 | VIRGDEVRQIAPGQTGKIADY | | 19 | 2286.2201 | RBD Abdala | 9 h (28 °C) | 9.120266 | 0.047135 | 3.087295 | 0.003314 | |
| 401 | 421 | VIRGDEVRQIAPGQTGKIADY | | 19 | 2286.2201 | MaxD | Max | 12.994112 | 0.058023 | 3.079449 | 0.001381 | 0.280104598 |
| 401 | 422 | VIRGDEVRQIAPGQTGKIADYN | | 20 | 2400.263 | RBD HEK 293-F | 10 s (ice) | 2.215794 | 0.036407 | 2.857831 | 0.000683 | |
| 401 | 422 | VIRGDEVRQIAPGQTGKIADYN | | 20 | 2400.263 | RBD HEK 293-F | 10 s (23 °C) | 3.538431 | 0.006503 | 2.849831 | 0.002121 | |
| 401 | 422 | VIRGDEVRQIAPGQTGKIADYN | | 20 | 2400.263 | RBD HEK 293-F | 100 s (23 °C) | 4.193874 | 0.016983 | 2.850828 | 0.001068 | |
| 401 | 422 | VIRGDEVRQIAPGQTGKIADYN | | 20 | 2400.263 | RBD HEK 293-F | 1,000 s (23 °C) | 5.464666 | 0.004658 | 2.85043 | 0.001441 | |
| 401 | 422 | VIRGDEVRQIAPGQTGKIADYN | | 20 | 2400.263 | RBD HEK 293-F | 10,000 s (23 °C) | 6.490306 | 0.022356 | 2.851009 | 0.000529 | |
| 401 | 422 | VIRGDEVRQIAPGQTGKIADYN | | 20 | 2400.263 | RBD HEK 293-F | 9 h (28 °C) | 8.511889 | 0.020911 | 2.843996 | 0.002091 | |
| 401 | 422 | VIRGDEVRQIAPGQTGKIADYN | | 20 | 2400.263 | RBD Abdala | 10 s (ice) | 2.238896 | 0.010134 | 2.856433 | 0.000759 | |
| 401 | 422 | VIRGDEVRQIAPGQTGKIADYN | | 20 | 2400.263 | RBD Abdala | 10 s (23 °C) | 3.493449 | 0.015459 | 2.852041 | 0.001841 | |
| 401 | 422 | VIRGDEVRQIAPGQTGKIADYN | | 20 | 2400.263 | RBD Abdala | 100 s (23 °C) | 4.075646 | 0.095049 | 2.85383 | 0.001505 | |
| 401 | 422 | VIRGDEVRQIAPGQTGKIADYN | | 20 | 2400.263 | RBD Abdala | 1,000 s (23 °C) | 5.507027 | 0.02338 | 2.853708 | 0.000249 | |
| 401 | 422 | VIRGDEVRQIAPGQTGKIADYN | | 20 | 2400.263 | RBD Abdala | 10,000 s (23 °C) | 6.553772 | 0.026613 | 2.847565 | 2.18E-05 | |
| 401 | 422 | VIRGDEVRQIAPGQTGKIADYN | | 20 | 2400.263 | RBD Abdala | 9 h (28 °C) | 9.422671 | 0.087455 | 2.843017 | 0.001928 | |
| 401 | 422 | VIRGDEVRQIAPGQTGKIADYN | | 20 | 2400.263 | MaxD | Max | 13.869428 | 0.05615 | 2.839458 | 0.001476 | 0.270030105 |
| 407 | 420 | VRQIAPGQTGKIAD | | 12 | 1453.8122 | RBD HEK 293-F | 10 s (ice) | 1.887897 | 0.016057 | 1.79341 | 0.025053 | |
| 407 | 420 | VRQIAPGQTGKIAD | | 12 | 1453.8122 | RBD HEK 293-F | 10 s (23 °C) | 2.518782 | 0.008663 | 1.801901 | 0.001357 | |
| 407 | 420 | VRQIAPGQTGKIAD | | 12 | 1453.8122 | RBD HEK 293-F | 100 s (23 °C) | 3.038964 | 0.007295 | 1.8044 | 0.002088 | |
| 407 | 420 | VRQIAPGQTGKIAD | | 12 | 1453.8122 | RBD HEK 293-F | 1,000 s (23 °C) | 4.242086 | 0.004852 | 1.777908 | 0.030937 | |
| 407 | 420 | VRQIAPGQTGKIAD | | 12 | 1453.8122 | RBD HEK 293-F | 10,000 s (23 °C) | 4.878952 | 0.00092 | 1.804889 | 0.000744 | |
| 407 | 420 | VRQIAPGQTGKIAD | | 12 | 1453.8122 | RBD HEK 293-F | 9 h (28 °C) | 5.694962 | 0.045271 | 1.803691 | 0.003911 | |
| 407 | 420 | VRQIAPGQTGKIAD | | 12 | 1453.8122 | RBD Abdala | 10 s (ice) | 1.899992 | 0.017159 | 1.793497 | 0.014708 | |
| 407 | 420 | VRQIAPGQTGKIAD | | 12 | 1453.8122 | RBD Abdala | 10 s (23 °C) | 2.496294 | 0.002742 | 1.783941 | 0.010259 | |
| 407 | 420 | VRQIAPGQTGKIAD | | 12 | 1453.8122 | RBD Abdala | 100 s (23 °C) | 2.989428 | 0.11503 | 1.769257 | 0.044525 | |
| 407 | 420 | VRQIAPGQTGKIAD | | 12 | 1453.8122 | RBD Abdala | 1,000 s (23 °C) | 4.233143 | 0.039721 | 1.724616 | 0.019897 | |

| | | | | | | | | | | | | |
|---|---|---|---|---|---|---|---|---|---|---|---|---|
| 407 | 420 | VRQIAPGQTGKIAD | | 12 | 1453.8122 | RBD Abdala | 10,000 s (23 °C) | 4.938581 | 0.043733 | 1.774747 | 0.029867 | |
| 407 | 420 | VRQIAPGQTGKIAD | | 12 | 1453.8122 | RBD Abdala | 9 h (28 °C) | 6.195225 | 0.049918 | 1.774475 | 0.033663 | |
| 407 | 420 | VRQIAPGQTGKIAD | | 12 | 1453.8122 | MaxD | Max | 8.501238 | 0.042529 | 1.797083 | 0.056134 | 0.254277368 |
| 407 | 421 | VRQIAPGQTGKIADY | | 13 | 1616.8755 | RBD HEK 293-F | 10 s (ice) | 1.800787 | 0.028182 | 2.695169 | 0.0015 | |
| 407 | 421 | VRQIAPGQTGKIADY | | 13 | 1616.8755 | RBD HEK 293-F | 10 s (23 °C) | 2.390622 | 0.00431 | 2.691332 | 0.002754 | |
| 407 | 421 | VRQIAPGQTGKIADY | | 13 | 1616.8755 | RBD HEK 293-F | 100 s (23 °C) | 2.92474 | 0.006034 | 2.692363 | 0.001349 | |
| 407 | 421 | VRQIAPGQTGKIADY | | 13 | 1616.8755 | RBD HEK 293-F | 1,000 s (23 °C) | 4.114391 | 0.027581 | 2.69126 | 0.003509 | |
| 407 | 421 | VRQIAPGQTGKIADY | | 13 | 1616.8755 | RBD HEK 293-F | 10,000 s (23 °C) | 4.735007 | 0.001578 | 2.693992 | 0.001125 | |
| 407 | 421 | VRQIAPGQTGKIADY | | 13 | 1616.8755 | RBD HEK 293-F | 9 h (28 °C) | 5.717292 | 0.047882 | 2.689606 | 0.001204 | |
| 407 | 421 | VRQIAPGQTGKIADY | | 13 | 1616.8755 | RBD Abdala | 10 s (ice) | 1.827322 | 0.03163 | 2.696006 | 0.002363 | |
| 407 | 421 | VRQIAPGQTGKIADY | | 13 | 1616.8755 | RBD Abdala | 10 s (23 °C) | 2.409381 | 0.007331 | 2.690876 | 0.002892 | |
| 407 | 421 | VRQIAPGQTGKIADY | | 13 | 1616.8755 | RBD Abdala | 100 s (23 °C) | 2.886205 | 0.087999 | 2.693397 | 0.002588 | |
| 407 | 421 | VRQIAPGQTGKIADY | | 13 | 1616.8755 | RBD Abdala | 1,000 s (23 °C) | 4.164127 | 0.022178 | 2.691147 | 0.001338 | |
| 407 | 421 | VRQIAPGQTGKIADY | | 13 | 1616.8755 | RBD Abdala | 10,000 s (23 °C) | 4.857413 | 0.008686 | 2.687034 | 0.001717 | |
| 407 | 421 | VRQIAPGQTGKIADY | | 13 | 1616.8755 | RBD Abdala | 9 h (28 °C) | 6.338881 | 0.031103 | 2.689693 | 0.002836 | |
| 407 | 421 | VRQIAPGQTGKIADY | | 13 | 1616.8755 | MaxD | Max | 9.099928 | 0.027645 | 2.688275 | 0.003193 | 0.263163725 |
| 407 | 422 | VRQIAPGQTGKIADYN | | 14 | 1730.9184 | RBD HEK 293-F | 10 s (ice) | 1.978598 | 0.035681 | 2.372099 | 0.003464 | |
| 407 | 422 | VRQIAPGQTGKIADYN | | 14 | 1730.9184 | RBD HEK 293-F | 10 s (23 °C) | 2.596336 | 0.006366 | 2.366513 | 0.002451 | |
| 407 | 422 | VRQIAPGQTGKIADYN | | 14 | 1730.9184 | RBD HEK 293-F | 100 s (23 °C) | 3.181135 | 0.024593 | 2.368701 | 0.000508 | |
| 407 | 422 | VRQIAPGQTGKIADYN | | 14 | 1730.9184 | RBD HEK 293-F | 1,000 s (23 °C) | 4.3901 | 0.035141 | 2.364075 | 0.006394 | |
| 407 | 422 | VRQIAPGQTGKIADYN | | 14 | 1730.9184 | RBD HEK 293-F | 10,000 s (23 °C) | 5.075403 | 0.007897 | 2.371107 | 0.001483 | |
| 407 | 422 | VRQIAPGQTGKIADYN | | 14 | 1730.9184 | RBD HEK 293-F | 9 h (28 °C) | 6.087077 | 0.043952 | 2.366453 | 0.000933 | |
| 407 | 422 | VRQIAPGQTGKIADYN | | 14 | 1730.9184 | RBD Abdala | 10 s (ice) | 1.982746 | 0.019507 | 2.370183 | 0.003002 | |
| 407 | 422 | VRQIAPGQTGKIADYN | | 14 | 1730.9184 | RBD Abdala | 10 s (23 °C) | 2.620819 | 0.012833 | 2.365069 | 0.003045 | |
| 407 | 422 | VRQIAPGQTGKIADYN | | 14 | 1730.9184 | RBD Abdala | 100 s (23 °C) | 3.072925 | 0.074434 | 2.366862 | 0.006197 | |
| 407 | 422 | VRQIAPGQTGKIADYN | | 14 | 1730.9184 | RBD Abdala | 1,000 s (23 °C) | 4.418862 | 0.001196 | 2.363213 | 0.002797 | |

| | | | | | | | | | | | | |
|---|---|---|---|---|---|---|---|---|---|---|---|---|
| 407 | 422 | VRQIAPGQTGKIADYN | | 14 | 1730.9184 | RBD Abdala | 10,000 s (23 °C) | 5.130252 | 0.000839 | 2.363123 | 0.003841 | |
| 407 | 422 | VRQIAPGQTGKIADYN | | 14 | 1730.9184 | RBD Abdala | 9 h (28 °C) | 6.84173 | 0.094952 | 2.362545 | 0.004004 | |
| 407 | 422 | VRQIAPGQTGKIADYN | | 14 | 1730.9184 | MaxD | Max | 10.277705 | 0.043156 | 2.358794 | 0.00533 | 0.227240226 |
| 407 | 431 | VRQIAPGQTGKIADYNYKLPDDFTG | | 22 | 2767.405 | RBD HEK 293-F | 10 s (ice) | 2.155318 | 0.044246 | 4.17717 | 0.01806 | |
| 407 | 431 | VRQIAPGQTGKIADYNYKLPDDFTG | | 22 | 2767.405 | RBD HEK 293-F | 10 s (23 °C) | 3.065878 | 0.043374 | 4.171029 | 0.023436 | |
| 407 | 431 | VRQIAPGQTGKIADYNYKLPDDFTG | | 22 | 2767.405 | RBD HEK 293-F | 100 s (23 °C) | 3.939229 | 0.004075 | 4.185023 | 0.028994 | |
| 407 | 431 | VRQIAPGQTGKIADYNYKLPDDFTG | | 22 | 2767.405 | RBD HEK 293-F | 1,000 s (23 °C) | 5.301229 | 0.012354 | 4.182681 | 0.027499 | |
| 407 | 431 | VRQIAPGQTGKIADYNYKLPDDFTG | | 22 | 2767.405 | RBD HEK 293-F | 10,000 s (23 °C) | 5.958169 | 0.111482 | 4.178089 | 0.021535 | |
| 407 | 431 | VRQIAPGQTGKIADYNYKLPDDFTG | | 22 | 2767.405 | RBD HEK 293-F | 9 h (28 °C) | 7.226693 | 0.101033 | 4.168223 | 0.008007 | |
| 407 | 431 | VRQIAPGQTGKIADYNYKLPDDFTG | | 22 | 2767.405 | RBD Abdala | 10 s (ice) | 2.144152 | 0.09517 | 4.179567 | 0.019655 | |
| 407 | 431 | VRQIAPGQTGKIADYNYKLPDDFTG | | 22 | 2767.405 | RBD Abdala | 10 s (23 °C) | 3.003871 | 0.075162 | 4.204801 | 0.007022 | |
| 407 | 431 | VRQIAPGQTGKIADYNYKLPDDFTG | | 22 | 2767.405 | RBD Abdala | 100 s (23 °C) | 3.866527 | 0.139968 | 4.217707 | 0.003511 | |
| 407 | 431 | VRQIAPGQTGKIADYNYKLPDDFTG | | 22 | 2767.405 | RBD Abdala | 1,000 s (23 °C) | 5.29078 | 0.041003 | 4.206942 | 0.006251 | |
| 407 | 431 | VRQIAPGQTGKIADYNYKLPDDFTG | | 22 | 2767.405 | RBD Abdala | 10,000 s (23 °C) | 5.975559 | 0.017932 | 4.186706 | 0.01431 | |
| 407 | 431 | VRQIAPGQTGKIADYNYKLPDDFTG | | 22 | 2767.405 | RBD Abdala | 9 h (28 °C) | 7.984953 | 0.218088 | 4.193658 | 0.025415 | |
| 407 | 431 | VRQIAPGQTGKIADYNYKLPDDFTG | | 22 | 2767.405 | MaxD | Max | 12.725122 | 0.042705 | 4.162304 | 0.003466 | 0.391142488 |
| 408 | 422 | RQIAPGQTGKIADYN | | 13 | 1631.85 | RBD HEK 293-F | 10 s (ice) | 1.818565 | 0.084759 | 2.202188 | 0.005399 | |
| 408 | 422 | RQIAPGQTGKIADYN | | 13 | 1631.85 | RBD HEK 293-F | 10 s (23 °C) | 2.368434 | 0.018006 | 2.199918 | 0.002744 | |
| 408 | 422 | RQIAPGQTGKIADYN | | 13 | 1631.85 | RBD HEK 293-F | 100 s (23 °C) | 2.689106 | 0.009719 | 2.203402 | 0.001881 | |
| 408 | 422 | RQIAPGQTGKIADYN | | 13 | 1631.85 | RBD HEK 293-F | 1,000 s (23 °C) | 4.043163 | 0.044523 | 2.194163 | 0.007044 | |
| 408 | 422 | RQIAPGQTGKIADYN | | 13 | 1631.85 | RBD HEK 293-F | 10,000 s (23 °C) | 4.330839 | 0.025898 | 2.204238 | 0.001463 | |
| 408 | 422 | RQIAPGQTGKIADYN | | 13 | 1631.85 | RBD HEK 293-F | 9 h (28 °C) | 5.002456 | 0.032856 | 2.199467 | 0.001395 | |
| 408 | 422 | RQIAPGQTGKIADYN | | 13 | 1631.85 | RBD Abdala | 10 s (ice) | 1.789282 | 0.038495 | 2.200106 | 0.006227 | |
| 408 | 422 | RQIAPGQTGKIADYN | | 13 | 1631.85 | RBD Abdala | 10 s (23 °C) | 2.308485 | 0.004518 | 2.192849 | 0.002646 | |
| 408 | 422 | RQIAPGQTGKIADYN | | 13 | 1631.85 | RBD Abdala | 100 s (23 °C) | 2.622576 | 0.106798 | 2.196524 | 0.012294 | |
| 408 | 422 | RQIAPGQTGKIADYN | | 13 | 1631.85 | RBD Abdala | 1,000 s (23 °C) | 4.028918 | 0.013479 | 2.186625 | 0.003741 | |

| | | | | | | | | | | | | |
|---|---|---|---|---|---|---|---|---|---|---|---|---|
| 408 | 422 | RQIAPGQTGKIADYN | | 13 | 1631.85 | RBD Abdala | 10,000 s (23 °C) | 4.45063 | 0.057282 | 2.19209 | 0.007122 | |
| 408 | 422 | RQIAPGQTGKIADYN | | 13 | 1631.85 | RBD Abdala | 9 h (28 °C) | 5.363047 | 0.095811 | 2.191163 | 0.006824 | |
| 408 | 422 | RQIAPGQTGKIADYN | | 13 | 1631.85 | MaxD | Max | 8.489562 | 0.232143 | 2.185727 | 0.016631 | 0.312586073 |
| 412 | 420 | PGQTGKIAD | | 8 | 886.4629 | RBD HEK 293-F | 10 s (ice) | 1.390199 | 0.006691 | 1.795454 | 0.022214 | |
| 412 | 420 | PGQTGKIAD | | 8 | 886.4629 | RBD HEK 293-F | 10 s (23 °C) | 1.795922 | 0.032252 | 1.801665 | 0.000943 | |
| 412 | 420 | PGQTGKIAD | | 8 | 886.4629 | RBD HEK 293-F | 100 s (23 °C) | 2.149211 | 0.017506 | 1.802866 | 0.001151 | |
| 412 | 420 | PGQTGKIAD | | 8 | 886.4629 | RBD HEK 293-F | 1,000 s (23 °C) | 3.135561 | 0.018169 | 1.77659 | 0.031368 | |
| 412 | 420 | PGQTGKIAD | | 8 | 886.4629 | RBD HEK 293-F | 10,000 s (23 °C) | 3.452514 | 0.067229 | 1.804465 | 3.15E-05 | |
| 412 | 420 | PGQTGKIAD | | 8 | 886.4629 | RBD HEK 293-F | 9 h (28 °C) | 3.807884 | 0.112988 | 1.802348 | 0.004165 | |
| 412 | 420 | PGQTGKIAD | | 8 | 886.4629 | RBD Abdala | 10 s (ice) | 1.422843 | 0.010576 | 1.793194 | 0.014154 | |
| 412 | 420 | PGQTGKIAD | | 8 | 886.4629 | RBD Abdala | 10 s (23 °C) | 1.789739 | 0.013746 | 1.785263 | 0.010527 | |
| 412 | 420 | PGQTGKIAD | | 8 | 886.4629 | RBD Abdala | 100 s (23 °C) | 2.091622 | 0.047688 | 1.77002 | 0.04212 | |
| 412 | 420 | PGQTGKIAD | | 8 | 886.4629 | RBD Abdala | 1,000 s (23 °C) | 3.138585 | 0.006544 | 1.727647 | 0.01926 | |
| 412 | 420 | PGQTGKIAD | | 8 | 886.4629 | RBD Abdala | 10,000 s (23 °C) | 3.53859 | 0.031955 | 1.776535 | 0.030094 | |
| 412 | 420 | PGQTGKIAD | | 8 | 886.4629 | RBD Abdala | 9 h (28 °C) | 4.169321 | 0.129193 | 1.776157 | 0.033539 | |
| 412 | 420 | PGQTGKIAD | | 8 | 886.4629 | MaxD | Max | 5.201754 | 0.011383 | 1.801882 | 0.001017 | 0.315558684 |
| 412 | 422 | PGQTGKIADYN | | 10 | 1163.5691 | RBD HEK 293-F | 10 s (ice) | 1.539304 | 0.015861 | 2.373485 | 0.002046 | |
| 412 | 422 | PGQTGKIADYN | | 10 | 1163.5691 | RBD HEK 293-F | 10 s (23 °C) | 1.993708 | 0.02507 | 2.367379 | 0.001667 | |
| 412 | 422 | PGQTGKIADYN | | 10 | 1163.5691 | RBD HEK 293-F | 100 s (23 °C) | 2.462679 | 0.026824 | 2.36768 | 0.000937 | |
| 412 | 422 | PGQTGKIADYN | | 10 | 1163.5691 | RBD HEK 293-F | 1,000 s (23 °C) | 3.489693 | 0.01493 | 2.366326 | 0.00335 | |
| 412 | 422 | PGQTGKIADYN | | 10 | 1163.5691 | RBD HEK 293-F | 10,000 s (23 °C) | 3.884677 | 0.035057 | 2.369708 | 0.000763 | |
| 412 | 422 | PGQTGKIADYN | | 10 | 1163.5691 | RBD HEK 293-F | 9 h (28 °C) | 4.366617 | 0.063203 | 2.367154 | 0.002367 | |
| 412 | 422 | PGQTGKIADYN | | 10 | 1163.5691 | RBD Abdala | 10 s (ice) | 1.556769 | 0.012086 | 2.372163 | 0.001805 | |
| 412 | 422 | PGQTGKIADYN | | 10 | 1163.5691 | RBD Abdala | 10 s (23 °C) | 2.007191 | 0.004941 | 2.365765 | 0.002079 | |
| 412 | 422 | PGQTGKIADYN | | 10 | 1163.5691 | RBD Abdala | 100 s (23 °C) | 2.404698 | 0.013391 | 2.368644 | 0.003577 | |
| 412 | 422 | PGQTGKIADYN | | 10 | 1163.5691 | RBD Abdala | 1,000 s (23 °C) | 3.546287 | 0.002213 | 2.362561 | 0.00247 | |

| | | | | | | | | | | | | |
|---|---|---|---|---|---|---|---|---|---|---|---|---|
| 412 | 422 | PGQTGKIADYN | | 10 | 1163.5691 | RBD Abdala | 10,000 s (23 °C) | 3.949617 | 0.003294 | 2.362426 | 0.003249 | |
| 412 | 422 | PGQTGKIADYN | | 10 | 1163.5691 | RBD Abdala | 9 h (28 °C) | 4.927005 | 0.079794 | 2.366194 | 0.005022 | |
| 412 | 422 | PGQTGKIADYN | | 10 | 1163.5691 | MaxD | Max | 7.130536 | 0.059733 | 2.360764 | 0.005039 | 0.249417263 |
| 415 | 420 | TGKIAD | | 5 | 604.3301 | RBD HEK 293-F | 10 s (ice) | 0.918601 | 0.026529 | 3.259633 | 0.027356 | |
| 415 | 420 | TGKIAD | | 5 | 604.3301 | RBD HEK 293-F | 10 s (23 °C) | 1.07787 | 0.049042 | 3.278243 | 0.038393 | |
| 415 | 420 | TGKIAD | | 5 | 604.3301 | RBD HEK 293-F | 100 s (23 °C) | 1.321259 | 0 | 3.313904 | 0.035463 | |
| 415 | 420 | TGKIAD | | 5 | 604.3301 | RBD HEK 293-F | 1,000 s (23 °C) | 2.035874 | 0.096257 | 3.272996 | 0.040427 | |
| 415 | 420 | TGKIAD | | 5 | 604.3301 | RBD HEK 293-F | 10,000 s (23 °C) | 2.275387 | 0.036708 | 3.272524 | 0.036492 | |
| 415 | 420 | TGKIAD | | 5 | 604.3301 | RBD HEK 293-F | 9 h (28 °C) | 2.255955 | 0.082079 | 3.246386 | 0.003882 | |
| 415 | 420 | TGKIAD | | 5 | 604.3301 | RBD Abdala | 10 s (ice) | 0.954451 | 0.019513 | 3.275891 | 0.032179 | |
| 415 | 420 | TGKIAD | | 5 | 604.3301 | RBD Abdala | 10 s (23 °C) | 1.074681 | 0.034065 | 3.322744 | 0.002302 | |
| 415 | 420 | TGKIAD | | 5 | 604.3301 | RBD Abdala | 100 s (23 °C) | 1.21821 | 0 | 3.327932 | 0.00258 | |
| 415 | 420 | TGKIAD | | 5 | 604.3301 | RBD Abdala | 1,000 s (23 °C) | 2.095939 | 0.019482 | 3.323495 | 0.006342 | |
| 415 | 420 | TGKIAD | | 5 | 604.3301 | RBD Abdala | 10,000 s (23 °C) | 2.250599 | 0.023712 | 3.282752 | 0.033128 | |
| 415 | 420 | TGKIAD | | 5 | 604.3301 | RBD Abdala | 9 h (28 °C) | 2.382745 | 0.070657 | 3.289781 | 0.03912 | |
| 415 | 420 | TGKIAD | | 5 | 604.3301 | MaxD | Max | 3.219901 | 0 | 3.246275 | 0.000554 | 0.322126105 |
| 415 | 421 | TGKIADY | | 6 | 767.3934 | RBD HEK 293-F | 10 s (ice) | 0.819453 | 0.049548 | 3.099588 | 0.002879 | |
| 415 | 421 | TGKIADY | | 6 | 767.3934 | RBD HEK 293-F | 10 s (23 °C) | 1.114297 | 0.028027 | 3.091106 | 0.00256 | |
| 415 | 421 | TGKIADY | | 6 | 767.3934 | RBD HEK 293-F | 100 s (23 °C) | 1.284966 | 0.009385 | 3.093414 | 0.001641 | |
| 415 | 421 | TGKIADY | | 6 | 767.3934 | RBD HEK 293-F | 1,000 s (23 °C) | 2.032307 | 0.014441 | 3.092061 | 0.000133 | |
| 415 | 421 | TGKIADY | | 6 | 767.3934 | RBD HEK 293-F | 10,000 s (23 °C) | 2.155093 | 0.018588 | 3.092711 | 2.58E-05 | |
| 415 | 421 | TGKIADY | | 6 | 767.3934 | RBD HEK 293-F | 9 h (28 °C) | 2.545088 | 0.034296 | 3.085568 | 0.001737 | |
| 415 | 421 | TGKIADY | | 6 | 767.3934 | RBD Abdala | 10 s (ice) | 0.873829 | 0.047448 | 3.097851 | 0.001704 | |
| 415 | 421 | TGKIADY | | 6 | 767.3934 | RBD Abdala | 10 s (23 °C) | 1.014875 | 0.078724 | 3.092907 | 0.000668 | |
| 415 | 421 | TGKIADY | | 6 | 767.3934 | RBD Abdala | 100 s (23 °C) | 1.222238 | 0.017799 | 3.096005 | 1.33E-05 | |
| 415 | 421 | TGKIADY | | 6 | 767.3934 | RBD Abdala | 1,000 s (23 °C) | 2.055227 | 0.04231 | 3.093975 | 0.001567 | |

| | | | | | | | | | | | | |
|---|---|---|---|---|---|---|---|---|---|---|---|---|
| 415 | 421 | TGKIADY | | 6 | 767.3934 | RBD Abdala | 10,000 s (23 °C) | 2.158491 | 0.093423 | 3.090591 | 0.000662 | |
| 415 | 421 | TGKIADY | | 6 | 767.3934 | RBD Abdala | 9 h (28 °C) | 2.642997 | 0.087358 | 3.087689 | 0.00095 | |
| 415 | 421 | TGKIADY | | 6 | 767.3934 | MaxD | Max | 3.656303 | 0.03102 | 3.086977 | 0.000341 | 0.358543333 |
| 415 | 422 | TGKIADYN | | 7 | 881.4363 | RBD HEK 293-F | 10 s (ice) | 1.072015 | 0.043437 | 3.136951 | 0.320535 | |
| 415 | 422 | TGKIADYN | | 7 | 881.4363 | RBD HEK 293-F | 10 s (23 °C) | 1.380763 | 0 | 2.848035 | 0 | |
| 415 | 422 | TGKIADYN | | 7 | 881.4363 | RBD HEK 293-F | 100 s (23 °C) | 1.713603 | 0.009346 | 3.195487 | 0.314781 | |
| 415 | 422 | TGKIADYN | | 7 | 881.4363 | RBD HEK 293-F | 1,000 s (23 °C) | 2.593857 | 0.075184 | 3.190341 | 0.314208 | |
| 415 | 422 | TGKIADYN | | 7 | 881.4363 | RBD HEK 293-F | 10,000 s (23 °C) | 2.793839 | 0.112517 | 3.203239 | 0.318301 | |
| 415 | 422 | TGKIADYN | | 7 | 881.4363 | RBD HEK 293-F | 9 h (28 °C) | 3.102923 | 0.102163 | 3.503251 | 0.005574 | |
| 415 | 422 | TGKIADYN | | 7 | 881.4363 | RBD Abdala | 10 s (ice) | 1.001536 | 0.027682 | 2.854703 | 0.002016 | |
| 415 | 422 | TGKIADYN | | 7 | 881.4363 | RBD Abdala | 10 s (23 °C) | 1.362676 | 0.053167 | 2.84997 | 0.003087 | |
| 415 | 422 | TGKIADYN | | 7 | 881.4363 | RBD Abdala | 100 s (23 °C) | 1.492665 | 0.00939 | 2.853415 | 0.004187 | |
| 415 | 422 | TGKIADYN | | 7 | 881.4363 | RBD Abdala | 1,000 s (23 °C) | 2.50388 | 0.094723 | 2.852175 | 0.000927 | |
| 415 | 422 | TGKIADYN | | 7 | 881.4363 | RBD Abdala | 10,000 s (23 °C) | 2.6587 | 0.055402 | 2.845565 | 0.001007 | |
| 415 | 422 | TGKIADYN | | 7 | 881.4363 | RBD Abdala | 9 h (28 °C) | 3.063185 | 0.131427 | 2.845341 | 0.003707 | |
| 415 | 422 | TGKIADYN | | 7 | 881.4363 | MaxD | Max | 4.841616 | 0.007462 | 3.502691 | 0.008851 | 0.271937444 |
| 418 | 441 | IADYNYKLPDDFTGCVIAWNSNNL | | 22 | 2746.2817 | RBD HEK 293-F | 10 s (ice) | 4.634202 | 0.026766 | 4.944053 | 0.007759 | |
| 418 | 441 | IADYNYKLPDDFTGCVIAWNSNNL | | 22 | 2746.2817 | RBD HEK 293-F | 10 s (23 °C) | 6.39214 | 0.029376 | 4.929247 | 0.010126 | |
| 418 | 441 | IADYNYKLPDDFTGCVIAWNSNNL | | 22 | 2746.2817 | RBD HEK 293-F | 100 s (23 °C) | 8.532433 | 0.057055 | 4.933654 | 0.016075 | |
| 418 | 441 | IADYNYKLPDDFTGCVIAWNSNNL | | 22 | 2746.2817 | RBD HEK 293-F | 1,000 s (23 °C) | 9.93343 | 0.056439 | 4.935819 | 0.010015 | |
| 418 | 441 | IADYNYKLPDDFTGCVIAWNSNNL | | 22 | 2746.2817 | RBD HEK 293-F | 10,000 s (23 °C) | 10.224462 | 0.030397 | 4.93406 | 0.00686 | |
| 418 | 441 | IADYNYKLPDDFTGCVIAWNSNNL | | 22 | 2746.2817 | RBD HEK 293-F | 9 h (28 °C) | 10.918201 | 0.15883 | 4.935837 | 0.006395 | |
| 418 | 441 | IADYNYKLPDDFTGCVIAWNSNNL | | 22 | 2746.2817 | RBD Abdala | 10 s (ice) | 4.597693 | 0.05649 | 4.942976 | 0.001376 | |
| 418 | 441 | IADYNYKLPDDFTGCVIAWNSNNL | | 22 | 2746.2817 | RBD Abdala | 10 s (23 °C) | 6.536417 | 0.058755 | 4.948117 | 0.002959 | |
| 418 | 441 | IADYNYKLPDDFTGCVIAWNSNNL | | 22 | 2746.2817 | RBD Abdala | 100 s (23 °C) | 8.561852 | 0.166786 | 4.950325 | 0.001056 | |
| 418 | 441 | IADYNYKLPDDFTGCVIAWNSNNL | | 22 | 2746.2817 | RBD Abdala | 1,000 s (23 °C) | 10.00323 | 0.0724 | 4.942631 | 0.002465 | |

| | | | | | | | | | | | | |
|---|---|---|---|---|---|---|---|---|---|---|---|---|
| 418 | 441 | IADYNYKLPDDFTGCVIAWNSNNL | | 22 | 2746.2817 | RBD Abdala | 10,000 s (23 °C) | 10.409673 | 0.057389 | 4.947248 | 0.001827 | |
| 418 | 441 | IADYNYKLPDDFTGCVIAWNSNNL | | 22 | 2746.2817 | RBD Abdala | 9 h (28 °C) | 11.079361 | 0.087316 | 4.94077 | 0.008556 | |
| 421 | 428 | YNYKLPDD | | 6 | 1027.4731 | RBD HEK 293-F | 10 s (ice) | 0.731771 | 0.028736 | 3.096129 | 0.001277 | |
| 421 | 428 | YNYKLPDD | | 6 | 1027.4731 | RBD HEK 293-F | 10 s (23 °C) | 0.793393 | 0.009963 | 3.090544 | 0.003705 | |
| 421 | 428 | YNYKLPDD | | 6 | 1027.4731 | RBD HEK 293-F | 100 s (23 °C) | 0.783138 | 0.012593 | 3.091228 | 0.001979 | |
| 421 | 428 | YNYKLPDD | | 6 | 1027.4731 | RBD HEK 293-F | 1,000 s (23 °C) | 0.851643 | 0.001589 | 3.094897 | 0.000533 | |
| 421 | 428 | YNYKLPDD | | 6 | 1027.4731 | RBD HEK 293-F | 10,000 s (23 °C) | 0.808819 | 0.012272 | 3.09818 | 0.000176 | |
| 421 | 428 | YNYKLPDD | | 6 | 1027.4731 | RBD HEK 293-F | 9 h (28 °C) | 0.948435 | 0.038512 | 3.09317 | 0.002582 | |
| 421 | 428 | YNYKLPDD | | 6 | 1027.4731 | RBD Abdala | 10 s (ice) | 0.7272 | 0.011792 | 3.095883 | 0.00243 | |
| 421 | 428 | YNYKLPDD | | 6 | 1027.4731 | RBD Abdala | 10 s (23 °C) | 0.774977 | 0.004859 | 3.090805 | 0.001685 | |
| 421 | 428 | YNYKLPDD | | 6 | 1027.4731 | RBD Abdala | 100 s (23 °C) | 0.711617 | 0.042723 | 3.095056 | 0.00193 | |
| 421 | 428 | YNYKLPDD | | 6 | 1027.4731 | RBD Abdala | 1,000 s (23 °C) | 0.753138 | 0.085808 | 3.096903 | 0.00206 | |
| 421 | 428 | YNYKLPDD | | 6 | 1027.4731 | RBD Abdala | 10,000 s (23 °C) | 0.823551 | 0.025578 | 3.092485 | 0.000437 | |
| 421 | 428 | YNYKLPDD | | 6 | 1027.4731 | RBD Abdala | 9 h (28 °C) | 1.126655 | 0.032714 | 3.094753 | 0.003585 | |
| 421 | 428 | YNYKLPDD | | 6 | 1027.4731 | MaxD | Max | 2.712595 | 0.053279 | 3.095512 | 0.001046 | 0.52410614 |
| 421 | 429 | YNYKLPDDF | | 7 | 1174.5415 | RBD HEK 293-F | 10 s (ice) | 0.637951 | 0.017041 | 4.673263 | 0.00885 | |
| 421 | 429 | YNYKLPDDF | | 7 | 1174.5415 | RBD HEK 293-F | 10 s (23 °C) | 0.846033 | 0.013734 | 4.661553 | 0.014979 | |
| 421 | 429 | YNYKLPDDF | | 7 | 1174.5415 | RBD HEK 293-F | 100 s (23 °C) | 1.368016 | 0.025393 | 4.672155 | 0.019075 | |
| 421 | 429 | YNYKLPDDF | | 7 | 1174.5415 | RBD HEK 293-F | 1,000 s (23 °C) | 1.425592 | 0.012358 | 4.672963 | 0.019502 | |
| 421 | 429 | YNYKLPDDF | | 7 | 1174.5415 | RBD HEK 293-F | 10,000 s (23 °C) | 1.425634 | 0.022013 | 4.673163 | 0.010052 | |
| 421 | 429 | YNYKLPDDF | | 7 | 1174.5415 | RBD HEK 293-F | 9 h (28 °C) | 1.627543 | 0.021311 | 4.66705 | 0.006406 | |
| 421 | 429 | YNYKLPDDF | | 7 | 1174.5415 | RBD Abdala | 10 s (ice) | 0.637195 | 0.012309 | 4.674008 | 0.00814 | |
| 421 | 429 | YNYKLPDDF | | 7 | 1174.5415 | RBD Abdala | 10 s (23 °C) | 0.81483 | 0.017857 | 4.687779 | 0.003592 | |
| 421 | 429 | YNYKLPDDF | | 7 | 1174.5415 | RBD Abdala | 100 s (23 °C) | 1.32786 | 0.060404 | 4.695822 | 0.002706 | |
| 421 | 429 | YNYKLPDDF | | 7 | 1174.5415 | RBD Abdala | 1,000 s (23 °C) | 1.398166 | 0.052677 | 4.693647 | 0.003208 | |
| 421 | 429 | YNYKLPDDF | | 7 | 1174.5415 | RBD Abdala | 10,000 s (23 °C) | 1.472974 | 0.02573 | 4.682338 | 0.009209 | |

| | | | | | | | | | | | | |
|---|---|---|---|---|---|---|---|---|---|---|---|---|
| 421 | 429 | YNYKLPDDF | | 7 | 1174.5415 | RBD Abdala | 9 h (28 °C) | 1.857578 | 0.024961 | 4.680874 | 0.014936 | |
| 421 | 429 | YNYKLPDDF | | 7 | 1174.5415 | MaxD | Max | 3.687242 | 0.018284 | 4.674216 | 0.002658 | 0.445527519 |
| 421 | 431 | YNYKLPDDFTG | | 9 | 1332.6107 | RBD HEK 293-F | 10 s (ice) | 0.539997 | 0.017184 | 4.328537 | 0.019098 | |
| 421 | 431 | YNYKLPDDFTG | | 9 | 1332.6107 | RBD HEK 293-F | 10 s (23 °C) | 1.032526 | 0.020992 | 4.318395 | 0.024375 | |
| 421 | 431 | YNYKLPDDFTG | | 9 | 1332.6107 | RBD HEK 293-F | 100 s (23 °C) | 1.871984 | 0.006656 | 4.332608 | 0.02667 | |
| 421 | 431 | YNYKLPDDFTG | | 9 | 1332.6107 | RBD HEK 293-F | 1,000 s (23 °C) | 1.938332 | 0.013523 | 4.327562 | 0.029591 | |
| 421 | 431 | YNYKLPDDFTG | | 9 | 1332.6107 | RBD HEK 293-F | 10,000 s (23 °C) | 1.983104 | 0.010118 | 4.330423 | 0.015825 | |
| 421 | 431 | YNYKLPDDFTG | | 9 | 1332.6107 | RBD HEK 293-F | 9 h (28 °C) | 2.35174 | 0.029273 | 4.314023 | 0.006557 | |
| 421 | 431 | YNYKLPDDFTG | | 9 | 1332.6107 | RBD Abdala | 10 s (ice) | 0.497117 | 0.036529 | 4.331409 | 0.019668 | |
| 421 | 431 | YNYKLPDDFTG | | 9 | 1332.6107 | RBD Abdala | 10 s (23 °C) | 0.915116 | 0.000623 | 4.354726 | 0.004268 | |
| 421 | 431 | YNYKLPDDFTG | | 9 | 1332.6107 | RBD Abdala | 100 s (23 °C) | 1.727315 | 0.084903 | 4.366518 | 0.003071 | |
| 421 | 431 | YNYKLPDDFTG | | 9 | 1332.6107 | RBD Abdala | 1,000 s (23 °C) | 1.838168 | 0.076294 | 4.358961 | 0.006013 | |
| 421 | 431 | YNYKLPDDFTG | | 9 | 1332.6107 | RBD Abdala | 10,000 s (23 °C) | 1.973218 | 0.021699 | 4.336806 | 0.01588 | |
| 421 | 431 | YNYKLPDDFTG | | 9 | 1332.6107 | RBD Abdala | 9 h (28 °C) | 2.588831 | 0.03765 | 4.345098 | 0.026716 | |
| 421 | 431 | YNYKLPDDFTG | | 9 | 1332.6107 | MaxD | Max | 5.09778 | 0.031398 | 4.315318 | 0.00301 | 0.403768421 |
| 422 | 431 | NYKLPDDFTG | | 8 | 1169.5473 | RBD HEK 293-F | 10 s (ice) | 0.565031 | 0.014117 | 3.969735 | 0.018688 | |
| 422 | 431 | NYKLPDDFTG | | 8 | 1169.5473 | RBD HEK 293-F | 10 s (23 °C) | 1.031341 | 0.000377 | 3.971735 | 0.024332 | |
| 422 | 431 | NYKLPDDFTG | | 8 | 1169.5473 | RBD HEK 293-F | 100 s (23 °C) | 1.804983 | 0.010381 | 3.978593 | 0.032482 | |
| 422 | 431 | NYKLPDDFTG | | 8 | 1169.5473 | RBD HEK 293-F | 1,000 s (23 °C) | 1.91431 | 0.020199 | 3.974583 | 0.030946 | |
| 422 | 431 | NYKLPDDFTG | | 8 | 1169.5473 | RBD HEK 293-F | 10,000 s (23 °C) | 1.901062 | 0.016304 | 3.97486 | 0.019495 | |
| 422 | 431 | NYKLPDDFTG | | 8 | 1169.5473 | RBD HEK 293-F | 9 h (28 °C) | 2.248258 | 0.027306 | 3.961256 | 0.004043 | |
| 422 | 431 | NYKLPDDFTG | | 8 | 1169.5473 | RBD Abdala | 10 s (ice) | 0.618193 | 0.032954 | 3.971804 | 0.019807 | |
| 422 | 431 | NYKLPDDFTG | | 8 | 1169.5473 | RBD Abdala | 10 s (23 °C) | 1.001176 | 1.61278E-05 | 3.998492 | 0.005731 | |
| 422 | 431 | NYKLPDDFTG | | 8 | 1169.5473 | RBD Abdala | 100 s (23 °C) | 1.751717 | 0.092458 | 4.007128 | 0.004168 | |
| 422 | 431 | NYKLPDDFTG | | 8 | 1169.5473 | RBD Abdala | 1,000 s (23 °C) | 1.906865 | 0.097127 | 4.001061 | 0.00694 | |
| 422 | 431 | NYKLPDDFTG | | 8 | 1169.5473 | RBD Abdala | 10,000 s (23 °C) | 2.0197 | 0.003062 | 3.977396 | 0.01674 | |

| | | | | | | | | | | | | |
|---|---|---|---|---|---|---|---|---|---|---|---|---|
| 422 | 431 | NYKLPDDFTG | | 8 | 1169.5473 | RBD Abdala | 9 h (28 °C) | 2.457239 | 0.042789 | 3.988413 | 0.027096 | |
| 422 | 431 | NYKLPDDFTG | | 8 | 1169.5473 | MaxD | Max | 4.228453 | 0.04029 | 3.960024 | 0.003428 | 0.443624605 |
| 423 | 429 | YKLPDDF | | 5 | 897.4353 | RBD HEK 293-F | 10 s (ice) | 0.7368 | 0.019391 | 4.355237 | 0.016791 | |
| 423 | 429 | YKLPDDF | | 5 | 897.4353 | RBD HEK 293-F | 10 s (23 °C) | 1.071046 | 0.00573 | 4.352917 | 0.024113 | |
| 423 | 429 | YKLPDDF | | 5 | 897.4353 | RBD HEK 293-F | 100 s (23 °C) | 1.560813 | 0.023227 | 4.361992 | 0.026483 | |
| 423 | 429 | YKLPDDF | | 5 | 897.4353 | RBD HEK 293-F | 1,000 s (23 °C) | 1.642898 | 0.016303 | 4.359088 | 0.029757 | |
| 423 | 429 | YKLPDDF | | 5 | 897.4353 | RBD HEK 293-F | 10,000 s (23 °C) | 1.572527 | 0.013317 | 4.361314 | 0.017176 | |
| 423 | 429 | YKLPDDF | | 5 | 897.4353 | RBD HEK 293-F | 9 h (28 °C) | 1.730612 | 0.040489 | 4.348428 | 0.005776 | |
| 423 | 429 | YKLPDDF | | 5 | 897.4353 | RBD Abdala | 10 s (ice) | 0.741808 | 0.022953 | 4.361692 | 0.018904 | |
| 423 | 429 | YKLPDDF | | 5 | 897.4353 | RBD Abdala | 10 s (23 °C) | 1.009724 | 0.024781 | 4.383223 | 0.004671 | |
| 423 | 429 | YKLPDDF | | 5 | 897.4353 | RBD Abdala | 100 s (23 °C) | 1.485225 | 0.060292 | 4.396118 | 0.003214 | |
| 423 | 429 | YKLPDDF | | 5 | 897.4353 | RBD Abdala | 1,000 s (23 °C) | 1.555573 | 0.045237 | 4.389107 | 0.00721 | |
| 423 | 429 | YKLPDDF | | 5 | 897.4353 | RBD Abdala | 10,000 s (23 °C) | 1.655478 | 0.021802 | 4.362928 | 0.016043 | |
| 423 | 429 | YKLPDDF | | 5 | 897.4353 | RBD Abdala | 9 h (28 °C) | 1.806562 | 0.048145 | 4.375929 | 0.02488 | |
| 423 | 429 | YKLPDDF | | 5 | 897.4353 | MaxD | Max | 2.600485 | 0.036634 | 4.344124 | 0.004801 | 0.452529474 |
| 423 | 431 | YKLPDDFTG | | 7 | 1055.5044 | RBD HEK 293-F | 10 s (ice) | 0.665649 | 0.015664 | 3.929431 | 0.017837 | |
| 423 | 431 | YKLPDDFTG | | 7 | 1055.5044 | RBD HEK 293-F | 10 s (23 °C) | 1.162677 | 0.027173 | 3.932936 | 0.02749 | |
| 423 | 431 | YKLPDDFTG | | 7 | 1055.5044 | RBD HEK 293-F | 100 s (23 °C) | 2.095248 | 0.015195 | 3.941472 | 0.034253 | |
| 423 | 431 | YKLPDDFTG | | 7 | 1055.5044 | RBD HEK 293-F | 1,000 s (23 °C) | 2.097998 | 0.024998 | 3.936417 | 0.033515 | |
| 423 | 431 | YKLPDDFTG | | 7 | 1055.5044 | RBD HEK 293-F | 10,000 s (23 °C) | 2.174045 | 0.012699 | 3.936188 | 0.020713 | |
| 423 | 431 | YKLPDDFTG | | 7 | 1055.5044 | RBD HEK 293-F | 9 h (28 °C) | 2.470814 | 0.026252 | 3.922513 | 0.006265 | |
| 423 | 431 | YKLPDDFTG | | 7 | 1055.5044 | RBD Abdala | 10 s (ice) | 0.66102 | 0.009019 | 3.936211 | 0.020957 | |
| 423 | 431 | YKLPDDFTG | | 7 | 1055.5044 | RBD Abdala | 10 s (23 °C) | 1.092072 | 0.003444 | 3.964256 | 0.003854 | |
| 423 | 431 | YKLPDDFTG | | 7 | 1055.5044 | RBD Abdala | 100 s (23 °C) | 2.080689 | 0 | 3.974635 | 0 | |
| 423 | 431 | YKLPDDFTG | | 7 | 1055.5044 | RBD Abdala | 1,000 s (23 °C) | 1.995453 | 0.075853 | 3.966894 | 0.006286 | |
| 423 | 431 | YKLPDDFTG | | 7 | 1055.5044 | RBD Abdala | 10,000 s (23 °C) | 2.206123 | 0.006404 | 3.939337 | 0.018821 | |

| | | | | | | | | | | | | |
|---|---|---|---|---|---|---|---|---|---|---|---|---|
| 423 | 431 | YKLPDDFTG | | 7 | 1055.5044 | RBD Abdala | 9 h (28 °C) | 2.580666 | 0.010427 | 3.953986 | 0.027817 | |
| 423 | 431 | YKLPDDFTG | | 7 | 1055.5044 | MaxD | Max | 4.016252 | 0.024139 | 3.920561 | 0.002793 | 0.396052331 |
| 425 | 431 | LPDDFTG | | 5 | 764.3461 | RBD HEK 293-F | 10 s (ice) | 0.447905 | 0.056617 | 3.929681 | 0.016592 | |
| 425 | 431 | LPDDFTG | | 5 | 764.3461 | RBD HEK 293-F | 10 s (23 °C) | 0.76041 | 0.010001 | 3.933758 | 0.026833 | |
| 425 | 431 | LPDDFTG | | 5 | 764.3461 | RBD HEK 293-F | 100 s (23 °C) | 1.497378 | 0.013728 | 3.944862 | 0.032216 | |
| 425 | 431 | LPDDFTG | | 5 | 764.3461 | RBD HEK 293-F | 1,000 s (23 °C) | 1.513275 | 0.044162 | 3.93875 | 0.033632 | |
| 425 | 431 | LPDDFTG | | 5 | 764.3461 | RBD HEK 293-F | 10,000 s (23 °C) | 1.424198 | 0 | 3.920638 | 0 | |
| 425 | 431 | LPDDFTG | | 5 | 764.3461 | RBD HEK 293-F | 9 h (28 °C) | 1.611232 | 0.088204 | 3.928156 | 0.003775 | |
| 425 | 431 | LPDDFTG | | 5 | 764.3461 | RBD Abdala | 10 s (ice) | 0.478735 | 0.038336 | 3.943411 | 0.017973 | |
| 425 | 431 | LPDDFTG | | 5 | 764.3461 | RBD Abdala | 10 s (23 °C) | 0.893794 | 0 | 3.97065 | 0 | |
| 425 | 431 | LPDDFTG | | 5 | 764.3461 | RBD Abdala | 100 s (23 °C) | 1.47752 | 0 | 3.978025 | 0 | |
| 425 | 431 | LPDDFTG | | 5 | 764.3461 | RBD Abdala | 1,000 s (23 °C) | 1.542763 | 0.051424 | 3.970707 | 0.008608 | |
| 425 | 431 | LPDDFTG | | 5 | 764.3461 | RBD Abdala | 10,000 s (23 °C) | 1.513255 | 0.065261 | 3.942783 | 0.016626 | |
| 425 | 431 | LPDDFTG | | 5 | 764.3461 | RBD Abdala | 9 h (28 °C) | 1.780876 | 0.11064 | 3.960123 | 0.026219 | |
| 425 | 431 | LPDDFTG | | 5 | 764.3461 | MaxD | Max | 2.650155 | 0.018455 | 3.922977 | 0.003218 | 0.442072632 |
| 426 | 431 | PDDFTG | | 5 | 651.262 | RBD HEK 293-F | 10 s (ice) | 0.418776 | 0.023669 | 3.935281 | 0.02009 | |
| 426 | 431 | PDDFTG | | 5 | 651.262 | RBD HEK 293-F | 10 s (23 °C) | 0.643013 | 0.00684 | 3.935338 | 0.028462 | |
| 426 | 431 | PDDFTG | | 5 | 651.262 | RBD HEK 293-F | 100 s (23 °C) | 1.368478 | 0.014155 | 3.946867 | 0.029613 | |
| 426 | 431 | PDDFTG | | 5 | 651.262 | RBD HEK 293-F | 1,000 s (23 °C) | 1.400267 | 0.024215 | 3.942719 | 0.034209 | |
| 426 | 431 | PDDFTG | | 5 | 651.262 | RBD HEK 293-F | 10,000 s (23 °C) | 1.228376 | 0.017404 | 3.943395 | 0.019673 | |
| 426 | 431 | PDDFTG | | 5 | 651.262 | RBD HEK 293-F | 9 h (28 °C) | 1.59181 | 0.039742 | 3.927198 | 0.00809 | |
| 426 | 431 | PDDFTG | | 5 | 651.262 | RBD Abdala | 10 s (ice) | 0.392023 | 0.022194 | 3.941765 | 0.020902 | |
| 426 | 431 | PDDFTG | | 5 | 651.262 | RBD Abdala | 10 s (23 °C) | 0.593727 | 0.000165 | 3.970387 | 0.00522 | |
| 426 | 431 | PDDFTG | | 5 | 651.262 | RBD Abdala | 100 s (23 °C) | 1.401608 | 0 | 3.981303 | 0 | |
| 426 | 431 | PDDFTG | | 5 | 651.262 | RBD Abdala | 1,000 s (23 °C) | 1.481124 | 0 | 3.980711 | 0 | |
| 426 | 431 | PDDFTG | | 5 | 651.262 | RBD Abdala | 10,000 s (23 °C) | 1.338601 | 0.00779 | 3.950061 | 0.01729 | |

| | | | | | | | | | | | | |
|---|---|---|---|---|---|---|---|---|---|---|---|---|
| 426 | 431 | PDDFTG | | 5 | 651.262 | RBD Abdala | 9 h (28 °C) | 1.600853 | 0.060648 | 3.960348 | 0.026127 | |
| 426 | 431 | PDDFTG | | 5 | 651.262 | MaxD | Max | 2.383415 | 0.05113 | 3.925723 | 0.002972 | 0.498228421 |
| 432 | 441 | CVIAWNSNNL | | 9 | 1133.5408 | RBD HEK 293-F | 10 s (ice) | 0.764316 | 0.003294 | 4.884845 | 0.009069 | |
| 432 | 441 | CVIAWNSNNL | | 9 | 1133.5408 | RBD HEK 293-F | 10 s (23 °C) | 1.365055 | 0.016409 | 4.871117 | 0.015431 | |
| 432 | 441 | CVIAWNSNNL | | 9 | 1133.5408 | RBD HEK 293-F | 100 s (23 °C) | 2.197684 | 0.066752 | 4.877164 | 0.017748 | |
| 432 | 441 | CVIAWNSNNL | | 9 | 1133.5408 | RBD HEK 293-F | 1,000 s (23 °C) | 2.767286 | 0.016042 | 4.879612 | 0.017027 | |
| 432 | 441 | CVIAWNSNNL | | 9 | 1133.5408 | RBD HEK 293-F | 10,000 s (23 °C) | 2.93302 | 0.037517 | 4.880512 | 0.01099 | |
| 432 | 441 | CVIAWNSNNL | | 9 | 1133.5408 | RBD HEK 293-F | 9 h (28 °C) | 2.961369 | 0.021968 | 4.878706 | 0.004308 | |
| 432 | 441 | CVIAWNSNNL | | 9 | 1133.5408 | RBD Abdala | 10 s (ice) | 0.790292 | 0.049883 | 4.885394 | 0.004144 | |
| 432 | 441 | CVIAWNSNNL | | 9 | 1133.5408 | RBD Abdala | 10 s (23 °C) | 1.270366 | 0.001916 | 4.895453 | 0.006015 | |
| 432 | 441 | CVIAWNSNNL | | 9 | 1133.5408 | RBD Abdala | 100 s (23 °C) | 2.106868 | 0.076656 | 4.903282 | 0.000959 | |
| 432 | 441 | CVIAWNSNNL | | 9 | 1133.5408 | RBD Abdala | 1,000 s (23 °C) | 2.79399 | 0.014807 | 4.894262 | 0.006882 | |
| 432 | 441 | CVIAWNSNNL | | 9 | 1133.5408 | RBD Abdala | 10,000 s (23 °C) | 2.929967 | 0.002428 | 4.888351 | 0.001457 | |
| 432 | 441 | CVIAWNSNNL | | 9 | 1133.5408 | RBD Abdala | 9 h (28 °C) | 3.009386 | 0.051167 | 4.88909 | 0.013288 | |
| 432 | 441 | CVIAWNSNNL | | 9 | 1133.5408 | MaxD | Max | 4.715836 | 0.085952 | 4.882531 | 0.0052 | 0.448440234 |
| 434 | 441 | IAWNSNNL | | 7 | 931.4632 | RBD HEK 293-F | 10 s (ice) | 1.058491 | 0.011271 | 4.227948 | 0.016928 | |
| 434 | 441 | IAWNSNNL | | 7 | 931.4632 | RBD HEK 293-F | 10 s (23 °C) | 1.818309 | 0.007708 | 4.224606 | 0.024382 | |
| 434 | 441 | IAWNSNNL | | 7 | 931.4632 | RBD HEK 293-F | 100 s (23 °C) | 3.071826 | 0.022317 | 4.233601 | 0.028579 | |
| 434 | 441 | IAWNSNNL | | 7 | 931.4632 | RBD HEK 293-F | 1,000 s (23 °C) | 3.966653 | 0.013693 | 4.229607 | 0.028972 | |
| 434 | 441 | IAWNSNNL | | 7 | 931.4632 | RBD HEK 293-F | 10,000 s (23 °C) | 4.156336 | 0.00133 | 4.23028 | 0.020152 | |
| 434 | 441 | IAWNSNNL | | 7 | 931.4632 | RBD HEK 293-F | 9 h (28 °C) | 4.161366 | 0.054559 | 4.218241 | 0.00628 | |
| 434 | 441 | IAWNSNNL | | 7 | 931.4632 | RBD Abdala | 10 s (ice) | 1.060913 | 0.02332 | 4.232626 | 0.01803 | |
| 434 | 441 | IAWNSNNL | | 7 | 931.4632 | RBD Abdala | 10 s (23 °C) | 1.781747 | 0.000537 | 4.256558 | 0.004497 | |
| 434 | 441 | IAWNSNNL | | 7 | 931.4632 | RBD Abdala | 100 s (23 °C) | 2.976724 | 0.105253 | 4.264445 | 0.003656 | |
| 434 | 441 | IAWNSNNL | | 7 | 931.4632 | RBD Abdala | 1,000 s (23 °C) | 3.921566 | 0.071251 | 4.256713 | 0.00579 | |
| 434 | 441 | IAWNSNNL | | 7 | 931.4632 | RBD Abdala | 10,000 s (23 °C) | 4.197594 | 0.042242 | 4.237426 | 0.014558 | |

| | | | | | | | | | | | | |
|---|---|---|---|---|---|---|---|---|---|---|---|---|
| 434 | 441 | IAWNSNNL | | 7 | 931.4632 | RBD Abdala | 9 h (28 °C) | 4.198388 | 0.060346 | 4.24678 | 0.024279 | |
| 434 | 441 | IAWNSNNL | | 7 | 931.4632 | MaxD | Max | 4.98996 | 0.043579 | 4.214094 | 0.002074 | 0.249630075 |
| 436 | 441 | WNSNNL | | 5 | 747.342 | RBD HEK 293-F | 10 s (ice) | 1.136065 | 0.012135 | 3.257279 | 0.005631 | |
| 436 | 441 | WNSNNL | | 5 | 747.342 | RBD HEK 293-F | 10 s (23 °C) | 1.847613 | 0.021538 | 3.258331 | 0.01219 | |
| 436 | 441 | WNSNNL | | 5 | 747.342 | RBD HEK 293-F | 100 s (23 °C) | 2.64316 | 0.010898 | 3.258205 | 0.012607 | |
| 436 | 441 | WNSNNL | | 5 | 747.342 | RBD HEK 293-F | 1,000 s (23 °C) | 3.271191 | 0.02168 | 3.255534 | 0.007925 | |
| 436 | 441 | WNSNNL | | 5 | 747.342 | RBD HEK 293-F | 10,000 s (23 °C) | 3.459666 | 0.002717 | 3.26087 | 0.0103 | |
| 436 | 441 | WNSNNL | | 5 | 747.342 | RBD HEK 293-F | 9 h (28 °C) | 3.483577 | 0.03069 | 3.249305 | 0.001473 | |
| 436 | 441 | WNSNNL | | 5 | 747.342 | RBD Abdala | 10 s (ice) | 1.111295 | 0.012406 | 3.259706 | 0.009037 | |
| 436 | 441 | WNSNNL | | 5 | 747.342 | RBD Abdala | 10 s (23 °C) | 1.760778 | 0.011115 | 3.268778 | 0.003339 | |
| 436 | 441 | WNSNNL | | 5 | 747.342 | RBD Abdala | 100 s (23 °C) | 2.540984 | 0.057045 | 3.269008 | 0.003958 | |
| 436 | 441 | WNSNNL | | 5 | 747.342 | RBD Abdala | 1,000 s (23 °C) | 3.234583 | 0.067168 | 3.268638 | 0.002037 | |
| 436 | 441 | WNSNNL | | 5 | 747.342 | RBD Abdala | 10,000 s (23 °C) | 3.423771 | 0.006483 | 3.254874 | 0.005529 | |
| 436 | 441 | WNSNNL | | 5 | 747.342 | RBD Abdala | 9 h (28 °C) | 3.472796 | 0.013875 | 3.261217 | 0.010034 | |
| 436 | 441 | WNSNNL | | 5 | 747.342 | MaxD | Max | 3.413285 | 0.083658 | 3.249005 | 0.004113 | 0.281413684 |
| 442 | 449 | DSKVGGNY | | 7 | 839.3894 | RBD HEK 293-F | 10 s (ice) | 4.110053 | 0.045594 | 1.321866 | 0.057594 | |
| 442 | 449 | DSKVGGNY | | 7 | 839.3894 | RBD HEK 293-F | 10 s (23 °C) | 4.626954 | 0.008588 | 1.336049 | 0.001424 | |
| 442 | 449 | DSKVGGNY | | 7 | 839.3894 | RBD HEK 293-F | 100 s (23 °C) | 4.639254 | 0.006526 | 1.340816 | 0.004014 | |
| 442 | 449 | DSKVGGNY | | 7 | 839.3894 | RBD HEK 293-F | 1,000 s (23 °C) | 4.673179 | 0.01478 | 1.28537 | 0.079373 | |
| 442 | 449 | DSKVGGNY | | 7 | 839.3894 | RBD HEK 293-F | 10,000 s (23 °C) | 4.624374 | 0.012877 | 1.339013 | 0.002688 | |
| 442 | 449 | DSKVGGNY | | 7 | 839.3894 | RBD HEK 293-F | 9 h (28 °C) | 4.645162 | 0.010104 | 1.352102 | 0.015705 | |
| 442 | 449 | DSKVGGNY | | 7 | 839.3894 | RBD Abdala | 10 s (ice) | 4.120812 | 0.025688 | 1.302457 | 0.042694 | |
| 442 | 449 | DSKVGGNY | | 7 | 839.3894 | RBD Abdala | 10 s (23 °C) | 4.61776 | 0.01853 | 1.285364 | 0.02959 | |
| 442 | 449 | DSKVGGNY | | 7 | 839.3894 | RBD Abdala | 100 s (23 °C) | 4.553151 | 0.059459 | 1.284146 | 0.090965 | |
| 442 | 449 | DSKVGGNY | | 7 | 839.3894 | RBD Abdala | 1,000 s (23 °C) | 4.622856 | 0.139838 | 1.146048 | 0.054375 | |
| 442 | 449 | DSKVGGNY | | 7 | 839.3894 | RBD Abdala | 10,000 s (23 °C) | 4.639429 | 0.011743 | 1.293547 | 0.075929 | |

| | | | | | | | | | | | | |
|---|---|---|---|---|---|---|---|---|---|---|---|---|
| 442 | 449 | DSKVGGNY | | 7 | 839.3894 | RBD Abdala | 9 h (28 °C) | 4.71627 | 0.007098 | 1.29149 | 0.071707 | |
| 442 | 449 | DSKVGGNY | | 7 | 839.3894 | MaxD | Max | 4.550611 | 0.025397 | 1.316336 | 0.078349 | 0.315697594 |
| 442 | 452 | DSKVGGNYNYL | | 10 | 1229.5797 | RBD HEK 293-F | 10 s (ice) | 4.971878 | 0.011713 | 3.589368 | 0.015498 | |
| 442 | 452 | DSKVGGNYNYL | | 10 | 1229.5797 | RBD HEK 293-F | 10 s (23 °C) | 5.849216 | 0.022791 | 3.598483 | 0.024709 | |
| 442 | 452 | DSKVGGNYNYL | | 10 | 1229.5797 | RBD HEK 293-F | 100 s (23 °C) | 6.30204 | 0.037418 | 3.603823 | 0.031253 | |
| 442 | 452 | DSKVGGNYNYL | | 10 | 1229.5797 | RBD HEK 293-F | 1,000 s (23 °C) | 6.666144 | 0.008182 | 3.597365 | 0.027883 | |
| 442 | 452 | DSKVGGNYNYL | | 10 | 1229.5797 | RBD HEK 293-F | 10,000 s (23 °C) | 7.133509 | 0.009047 | 3.59699 | 0.021658 | |
| 442 | 452 | DSKVGGNYNYL | | 10 | 1229.5797 | RBD HEK 293-F | 9 h (28 °C) | 7.121149 | 0.028615 | 3.5853 | 0.0049 | |
| 442 | 452 | DSKVGGNYNYL | | 10 | 1229.5797 | RBD Abdala | 10 s (ice) | 4.991452 | 0.024795 | 3.598389 | 0.018143 | |
| 442 | 452 | DSKVGGNYNYL | | 10 | 1229.5797 | RBD Abdala | 10 s (23 °C) | 5.852826 | 0.014864 | 3.624344 | 0.001745 | |
| 442 | 452 | DSKVGGNYNYL | | 10 | 1229.5797 | RBD Abdala | 100 s (23 °C) | 6.08624 | 0.203525 | 3.628447 | 0.00362 | |
| 442 | 452 | DSKVGGNYNYL | | 10 | 1229.5797 | RBD Abdala | 1,000 s (23 °C) | 6.490088 | 0.166762 | 3.623371 | 0.006867 | |
| 442 | 452 | DSKVGGNYNYL | | 10 | 1229.5797 | RBD Abdala | 10,000 s (23 °C) | 7.16555 | 0.007472 | 3.597969 | 0.01713 | |
| 442 | 452 | DSKVGGNYNYL | | 10 | 1229.5797 | RBD Abdala | 9 h (28 °C) | 7.168645 | 0.021362 | 3.61079 | 0.026397 | |
| 442 | 452 | DSKVGGNYNYL | | 10 | 1229.5797 | MaxD | Max | 6.979835 | 0.034741 | 3.592786 | 0.003869 | 0.265280526 |
| 453 | 467 | YRLFRKSNLKPFERD | | 13 | 1969.0766 | RBD HEK 293-F | 10 s (ice) | 0.877572 | 0.040091 | 2.967838 | 0.002644 | |
| 453 | 467 | YRLFRKSNLKPFERD | | 13 | 1969.0766 | RBD HEK 293-F | 10 s (23 °C) | 1.238467 | 0.016298 | 2.957629 | 0.005191 | |
| 453 | 467 | YRLFRKSNLKPFERD | | 13 | 1969.0766 | RBD HEK 293-F | 100 s (23 °C) | 1.618112 | 0.071285 | 2.961724 | 0.002457 | |
| 453 | 467 | YRLFRKSNLKPFERD | | 13 | 1969.0766 | RBD HEK 293-F | 1,000 s (23 °C) | 2.226288 | 0.00068 | 2.963579 | 0.001788 | |
| 453 | 467 | YRLFRKSNLKPFERD | | 13 | 1969.0766 | RBD HEK 293-F | 10,000 s (23 °C) | 3.198295 | 0.12479 | 2.96726 | 0.001264 | |
| 453 | 467 | YRLFRKSNLKPFERD | | 13 | 1969.0766 | RBD HEK 293-F | 9 h (28 °C) | 4.743062 | 0.171977 | 2.961298 | 0.001347 | |
| 453 | 467 | YRLFRKSNLKPFERD | | 13 | 1969.0766 | RBD Abdala | 10 s (ice) | 0.949881 | 0.019275 | 2.966801 | 0.002378 | |
| 453 | 467 | YRLFRKSNLKPFERD | | 13 | 1969.0766 | RBD Abdala | 10 s (23 °C) | 1.202084 | 0.006473 | 2.958741 | 0.002081 | |
| 453 | 467 | YRLFRKSNLKPFERD | | 13 | 1969.0766 | RBD Abdala | 100 s (23 °C) | 1.569816 | 0.008485 | 2.967913 | 0.001452 | |
| 453 | 467 | YRLFRKSNLKPFERD | | 13 | 1969.0766 | RBD Abdala | 1,000 s (23 °C) | 2.372938 | 0.01662 | 2.964037 | 0.001228 | |
| 453 | 467 | YRLFRKSNLKPFERD | | 13 | 1969.0766 | RBD Abdala | 10,000 s (23 °C) | 3.252522 | 0.050109 | 2.959844 | 0.001517 | |

| | | | | | | | | | | | |
|---|---|---|---|---|---|---|---|---|---|---|---|
| 453 | 467 | YRLFRKSNLKPFERD | | 13 | 1969.0766 | RBD Abdala | 9 h (28 °C) | 5.23768 | 0.130374 | 2.962451 | 0.003186 | |
| 453 | 467 | YRLFRKSNLKPFERD | | 13 | 1969.0766 | MaxD | Max | 6.403073 | 0.045135 | 2.960041 | 0.003354 | 0.481532551 |
| 453 | 469 | YRLFRKSNLKPFERDIS | | 15 | 2169.1927 | RBD HEK 293-F | 10 s (ice) | 1.293621 | 0.023785 | 3.268367 | 0.038546 | |
| 453 | 469 | YRLFRKSNLKPFERDIS | | 15 | 2169.1927 | RBD HEK 293-F | 10 s (23 °C) | 2.286645 | 0.005302 | 3.282046 | 0.044051 | |
| 453 | 469 | YRLFRKSNLKPFERDIS | | 15 | 2169.1927 | RBD HEK 293-F | 100 s (23 °C) | 2.845777 | 0.022856 | 3.289526 | 0.049304 | |
| 453 | 469 | YRLFRKSNLKPFERDIS | | 15 | 2169.1927 | RBD HEK 293-F | 1,000 s (23 °C) | 3.958454 | 0.023226 | 3.27811 | 0.043476 | |
| 453 | 469 | YRLFRKSNLKPFERDIS | | 15 | 2169.1927 | RBD HEK 293-F | 10,000 s (23 °C) | 5.002007 | 0.131466 | 3.280266 | 0.045109 | |
| 453 | 469 | YRLFRKSNLKPFERDIS | | 15 | 2169.1927 | RBD HEK 293-F | 9 h (28 °C) | 6.801471 | 0.151037 | 3.244604 | 0.006415 | |
| 453 | 469 | YRLFRKSNLKPFERDIS | | 15 | 2169.1927 | RBD Abdala | 10 s (ice) | 1.312316 | 0.031198 | 3.278904 | 0.037908 | |
| 453 | 469 | YRLFRKSNLKPFERDIS | | 15 | 2169.1927 | RBD Abdala | 10 s (23 °C) | 2.284278 | 0.006539 | 3.331046 | 0.000397 | |
| 453 | 469 | YRLFRKSNLKPFERDIS | | 15 | 2169.1927 | RBD Abdala | 100 s (23 °C) | 2.839975 | 0.029858 | 3.334787 | 0.001243 | |
| 453 | 469 | YRLFRKSNLKPFERDIS | | 15 | 2169.1927 | RBD Abdala | 1,000 s (23 °C) | 4.09997 | 0.02441 | 3.328996 | 0.004905 | |
| 453 | 469 | YRLFRKSNLKPFERDIS | | 15 | 2169.1927 | RBD Abdala | 10,000 s (23 °C) | 5.190359 | 0.059076 | 3.288583 | 0.034719 | |
| 453 | 469 | YRLFRKSNLKPFERDIS | | 15 | 2169.1927 | RBD Abdala | 9 h (28 °C) | 7.27224 | 0.076939 | 3.291289 | 0.042856 | |
| 453 | 469 | YRLFRKSNLKPFERDIS | | 15 | 2169.1927 | MaxD | Max | 8.424452 | 0.051729 | 3.237277 | 0.005273 | 0.408810386 |
| 453 | 471 | YRLFRKSNLKPFERDISTE | | 17 | 2399.283 | RBD HEK 293-F | 10 s (ice) | 1.910289 | 0.054911 | 3.363364 | 0.042675 | |
| 453 | 471 | YRLFRKSNLKPFERDISTE | | 17 | 2399.283 | RBD HEK 293-F | 10 s (23 °C) | 3.280928 | 0.015027 | 3.374855 | 0.05284 | |
| 453 | 471 | YRLFRKSNLKPFERDISTE | | 17 | 2399.283 | RBD HEK 293-F | 100 s (23 °C) | 4.366944 | 0.075572 | 3.373363 | 0.063022 | |
| 453 | 471 | YRLFRKSNLKPFERDISTE | | 17 | 2399.283 | RBD HEK 293-F | 1,000 s (23 °C) | 5.669844 | 0.014032 | 3.370056 | 0.058217 | |
| 453 | 471 | YRLFRKSNLKPFERDISTE | | 17 | 2399.283 | RBD HEK 293-F | 10,000 s (23 °C) | 6.903283 | 0.103616 | 3.374791 | 0.052293 | |
| 453 | 471 | YRLFRKSNLKPFERDISTE | | 17 | 2399.283 | RBD HEK 293-F | 9 h (28 °C) | 8.834889 | 0.099731 | 3.335278 | 0.011882 | |
| 453 | 471 | YRLFRKSNLKPFERDISTE | | 17 | 2399.283 | RBD Abdala | 10 s (ice) | 1.952041 | 0.013692 | 3.381015 | 0.041623 | |
| 453 | 471 | YRLFRKSNLKPFERDISTE | | 17 | 2399.283 | RBD Abdala | 10 s (23 °C) | 3.279571 | 0.032987 | 3.427641 | 0.003027 | |
| 453 | 471 | YRLFRKSNLKPFERDISTE | | 17 | 2399.283 | RBD Abdala | 100 s (23 °C) | 4.397616 | 0.04001 | 3.433367 | 0.000557 | |
| 453 | 471 | YRLFRKSNLKPFERDISTE | | 17 | 2399.283 | RBD Abdala | 1,000 s (23 °C) | 5.890628 | 0.069053 | 3.426039 | 0.005864 | |
| 453 | 471 | YRLFRKSNLKPFERDISTE | | 17 | 2399.283 | RBD Abdala | 10,000 s (23 °C) | 7.108413 | 0.041582 | 3.381325 | 0.041619 | |

| | | | | | | | | | | | | |
|---|---|---|---|---|---|---|---|---|---|---|---|---|
| 453 | 471 | YRLFRKSNLKPFERDISTE | | 17 | 2399.283 | RBD Abdala | 9 h (28 °C) | 9.390514 | 0.046499 | 3.388197 | 0.053868 | |
| 453 | 471 | YRLFRKSNLKPFERDISTE | | 17 | 2399.283 | MaxD | Max | 10.565684 | 0.065738 | 3.327474 | 0.010071 | 0.34577808 |
| 453 | 471 | YRLFRKSNLKPFERDISTE | Man1 (S/T 18/19) | 17 | 2399.283 | RBD Abdala | 10 s (ice) | 1.717500 | 0.068300 | 3.381015 | 0.041623 | |
| 453 | 471 | YRLFRKSNLKPFERDISTE | Man1 (S/T 18/19) | 17 | 2399.283 | RBD Abdala | 10 s (23 °C) | 3.274800 | 0.014600 | 3.427641 | 0.003027 | |
| 453 | 471 | YRLFRKSNLKPFERDISTE | Man1 (S/T 18/19) | 17 | 2399.283 | RBD Abdala | 100 s (23 °C) | 4.317200 | 0.104900 | 3.433367 | 0.000557 | |
| 453 | 471 | YRLFRKSNLKPFERDISTE | Man1 (S/T 18/19) | 17 | 2399.283 | RBD Abdala | 1,000 s (23 °C) | 5.934200 | 0.038100 | 3.426039 | 0.005864 | |
| 453 | 471 | YRLFRKSNLKPFERDISTE | Man1 (S/T 18/19) | 17 | 2399.283 | RBD Abdala | 10,000 s (23 °C) | 7.268700 | 0.067200 | 3.381325 | 0.041619 | |
| 453 | 471 | YRLFRKSNLKPFERDISTE | Man1 (S/T 18/19) | 17 | 2399.283 | RBD Abdala | 9 h (28 °C) | 9.539300 | 0.093300 | 3.388197 | 0.053868 | |
| 453 | 471 | YRLFRKSNLKPFERDISTE | Man1 (S/T 18/19) | 17 | 2399.283 | MaxD | Max | 10.831100 | 0.112300 | 3.327474 | 0.010071 | 0.329343653 |
| 456 | 467 | FRKSNLKPFERD | | 10 | 1536.8281 | RBD HEK 293-F | 10 s (ice) | 0.883385 | 0.033091 | 1.99357 | 0.007848 | |
| 456 | 467 | FRKSNLKPFERD | | 10 | 1536.8281 | RBD HEK 293-F | 10 s (23 °C) | 1.2827 | 0.024027 | 1.990855 | 0.001654 | |
| 456 | 467 | FRKSNLKPFERD | | 10 | 1536.8281 | RBD HEK 293-F | 100 s (23 °C) | 1.500545 | 0.047392 | 1.990003 | 0.000993 | |
| 456 | 467 | FRKSNLKPFERD | | 10 | 1536.8281 | RBD HEK 293-F | 1,000 s (23 °C) | 1.715966 | 0.028263 | 1.984915 | 0.015557 | |
| 456 | 467 | FRKSNLKPFERD | | 10 | 1536.8281 | RBD HEK 293-F | 10,000 s (23 °C) | 2.629215 | 0.039946 | 1.996455 | 0.002586 | |
| 456 | 467 | FRKSNLKPFERD | | 10 | 1536.8281 | RBD HEK 293-F | 9 h (28 °C) | 3.814481 | 0.108916 | 1.996128 | 0.002285 | |
| 456 | 467 | FRKSNLKPFERD | | 10 | 1536.8281 | RBD Abdala | 10 s (ice) | 0.928997 | 0.01669 | 1.991577 | 0.007729 | |
| 456 | 467 | FRKSNLKPFERD | | 10 | 1536.8281 | RBD Abdala | 10 s (23 °C) | 1.255963 | 0.003707 | 1.98434 | 0.007167 | |
| 456 | 467 | FRKSNLKPFERD | | 10 | 1536.8281 | RBD Abdala | 100 s (23 °C) | 1.476556 | 0.001345 | 1.978344 | 0.019261 | |
| 456 | 467 | FRKSNLKPFERD | | 10 | 1536.8281 | RBD Abdala | 1,000 s (23 °C) | 1.82733 | 0.060721 | 1.968403 | 0.005195 | |
| 456 | 467 | FRKSNLKPFERD | | 10 | 1536.8281 | RBD Abdala | 10,000 s (23 °C) | 2.664376 | 0.042408 | 1.983594 | 0.012392 | |
| 456 | 467 | FRKSNLKPFERD | | 10 | 1536.8281 | RBD Abdala | 9 h (28 °C) | 4.10376 | 0.098203 | 1.983888 | 0.013837 | |
| 456 | 467 | FRKSNLKPFERD | | 10 | 1536.8281 | MaxD | Max | 4.684852 | 0.121268 | 1.991195 | 0.002277 | 0.506857684 |
| 456 | 470 | FRKSNLKPFERDIST | | 13 | 1837.9919 | RBD HEK 293-F | 10 s (ice) | 1.826794 | 0.054995 | 2.662104 | 0.001203 | |
| 456 | 470 | FRKSNLKPFERDIST | | 13 | 1837.9919 | RBD HEK 293-F | 10 s (23 °C) | 3.036387 | 0.023248 | 2.653219 | 0.003344 | |
| 456 | 470 | FRKSNLKPFERDIST | | 13 | 1837.9919 | RBD HEK 293-F | 100 s (23 °C) | 3.548792 | 0.042088 | 2.654954 | 0.002115 | |
| 456 | 470 | FRKSNLKPFERDIST | | 13 | 1837.9919 | RBD HEK 293-F | 1,000 s (23 °C) | 4.3367 | 0.02269 | 2.655632 | 0.002739 | |

| | | | | | | | | | | | | |
|---|---|---|---|---|---|---|---|---|---|---|---|---|
| 456 | 470 | FRKSNLKPFERDIST | | 13 | 1837.9919 | RBD HEK 293-F | 10,000 s (23 °C) | 5.403723 | 0.065599 | 2.657155 | 0.000102 | |
| 456 | 470 | FRKSNLKPFERDIST | | 13 | 1837.9919 | RBD HEK 293-F | 9 h (28 °C) | 6.822642 | 0.108504 | 2.6496 | 0.001836 | |
| 456 | 470 | FRKSNLKPFERDIST | | 13 | 1837.9919 | RBD Abdala | 10 s (ice) | 1.869118 | 0.006088 | 2.66273 | 0.002074 | |
| 456 | 470 | FRKSNLKPFERDIST | | 13 | 1837.9919 | RBD Abdala | 10 s (23 °C) | 3.048414 | 0.011826 | 2.654341 | 0.001832 | |
| 456 | 470 | FRKSNLKPFERDIST | | 13 | 1837.9919 | RBD Abdala | 100 s (23 °C) | 3.471227 | 0.027313 | 2.657491 | 0.00185 | |
| 456 | 470 | FRKSNLKPFERDIST | | 13 | 1837.9919 | RBD Abdala | 1,000 s (23 °C) | 4.445876 | 0.070714 | 2.655517 | 0.001304 | |
| 456 | 470 | FRKSNLKPFERDIST | | 13 | 1837.9919 | RBD Abdala | 10,000 s (23 °C) | 5.482448 | 0.083755 | 2.649606 | 0.000108 | |
| 456 | 470 | FRKSNLKPFERDIST | | 13 | 1837.9919 | RBD Abdala | 9 h (28 °C) | 7.225819 | 0.08665 | 2.649512 | 0.001708 | |
| 456 | 470 | FRKSNLKPFERDIST | | 13 | 1837.9919 | MaxD | Max | 7.525982 | 0.038359 | 2.646208 | 0.000932 | 0.390608745 |
| 456 | 471 | FRKSNLKPFERDISTE | | 14 | 1967.0345 | RBD HEK 293-F | 10 s (ice) | 1.978207 | 0.051823 | 2.741445 | 0.001427 | |
| 456 | 471 | FRKSNLKPFERDISTE | | 14 | 1967.0345 | RBD HEK 293-F | 10 s (23 °C) | 3.443614 | 0.008552 | 2.730787 | 0.002544 | |
| 456 | 471 | FRKSNLKPFERDISTE | | 14 | 1967.0345 | RBD HEK 293-F | 100 s (23 °C) | 4.351415 | 0.051761 | 2.72985 | 0.000709 | |
| 456 | 471 | FRKSNLKPFERDISTE | | 14 | 1967.0345 | RBD HEK 293-F | 1,000 s (23 °C) | 5.188562 | 0.036776 | 2.73214 | 0.001332 | |
| 456 | 471 | FRKSNLKPFERDISTE | | 14 | 1967.0345 | RBD HEK 293-F | 10,000 s (23 °C) | 6.372901 | 0.014159 | 2.732169 | 0.001301 | |
| 456 | 471 | FRKSNLKPFERDISTE | | 14 | 1967.0345 | RBD HEK 293-F | 9 h (28 °C) | 7.845064 | 0.077264 | 2.724658 | 0.001701 | |
| 456 | 471 | FRKSNLKPFERDISTE | | 14 | 1967.0345 | RBD Abdala | 10 s (ice) | 1.992441 | 0.029581 | 2.740771 | 0.001431 | |
| 456 | 471 | FRKSNLKPFERDISTE | | 14 | 1967.0345 | RBD Abdala | 10 s (23 °C) | 3.421906 | 0.011173 | 2.733515 | 0.001507 | |
| 456 | 471 | FRKSNLKPFERDISTE | | 14 | 1967.0345 | RBD Abdala | 100 s (23 °C) | 4.330276 | 0.04502 | 2.732878 | 0.002962 | |
| 456 | 471 | FRKSNLKPFERDISTE | | 14 | 1967.0345 | RBD Abdala | 1,000 s (23 °C) | 5.306804 | 0.078379 | 2.734625 | 0.002002 | |
| 456 | 471 | FRKSNLKPFERDISTE | | 14 | 1967.0345 | RBD Abdala | 10,000 s (23 °C) | 6.450277 | 0.05815 | 2.72918 | 0.001112 | |
| 456 | 471 | FRKSNLKPFERDISTE | | 14 | 1967.0345 | RBD Abdala | 9 h (28 °C) | 8.217673 | 0.051816 | 2.724166 | 0.003313 | |
| 456 | 471 | FRKSNLKPFERDISTE | | 14 | 1967.0345 | MaxD | Max | 8.684339 | 0.071226 | 2.723723 | 0.002123 | 0.34704218 |
| 456 | 471 | FRKSNLKPFERDISTE | Man1 (S/T 18/19) | 14 | 1967.0345 | RBD Abdala | 10 s (ice) | 1.86120 | 0.03430 | 2.740771 | 0.001431 | |
| 456 | 471 | FRKSNLKPFERDISTE | Man1 (S/T 18/19) | 14 | 1967.0345 | RBD Abdala | 10 s (23 °C) | 3.43480 | 0.06760 | 2.733515 | 0.001507 | |
| 456 | 471 | FRKSNLKPFERDISTE | Man1 (S/T 18/19) | 14 | 1967.0345 | RBD Abdala | 100 s (23 °C) | 4.27100 | 0.09940 | 2.732878 | 0.002962 | |
| 456 | 471 | FRKSNLKPFERDISTE | Man1 (S/T 18/19) | 14 | 1967.0345 | RBD Abdala | 1,000 s (23 °C) | 5.45260 | 0.03510 | 2.734625 | 0.002002 | |

| | | | | | | | | | | | | |
|---|---|---|---|---|---|---|---|---|---|---|---|---|
| 456 | 471 | FRKSNLKPFERDISTE | Man1 (S/T 18/19) | 14 | 1967.0345 | RBD Abdala | 10,000 s (23 °C) | 6.71910 | 0.08030 | 2.72918 | 0.001112 | |
| 456 | 471 | FRKSNLKPFERDISTE | Man1 (S/T 18/19) | 14 | 1967.0345 | RBD Abdala | 9 h (28 °C) | 8.49360 | 0.06810 | 2.724166 | 0.003313 | |
| 456 | 471 | FRKSNLKPFERDISTE | Man1 (S/T 18/19) | 14 | 1967.0345 | MaxD | Max | 8.84880 | 0.17120 | 2.723723 | 0.002123 | 0.334676692 |
| 471 | 486 | EIYQAGSTPCNGVEGF | | 14 | 1671.7319 | RBD HEK 293-F | 10 s (ice) | 3.593442 | 0.057027 | 4.479215 | 0.014095 | |
| 471 | 486 | EIYQAGSTPCNGVEGF | | 14 | 1671.7319 | RBD HEK 293-F | 10 s (23 °C) | 5.751241 | 0.051633 | 4.471911 | 0.020773 | |
| 471 | 486 | EIYQAGSTPCNGVEGF | | 14 | 1671.7319 | RBD HEK 293-F | 100 s (23 °C) | 7.200566 | 0.210326 | 4.480156 | 0.023774 | |
| 471 | 486 | EIYQAGSTPCNGVEGF | | 14 | 1671.7319 | RBD HEK 293-F | 1,000 s (23 °C) | 7.686998 | 0.056881 | 4.477177 | 0.024053 | |
| 471 | 486 | EIYQAGSTPCNGVEGF | | 14 | 1671.7319 | RBD HEK 293-F | 10,000 s (23 °C) | 7.608309 | 0.076716 | 4.478588 | 0.016754 | |
| 471 | 486 | EIYQAGSTPCNGVEGF | | 14 | 1671.7319 | RBD HEK 293-F | 9 h (28 °C) | 7.601228 | 0.079535 | 4.467252 | 0.005239 | |
| 471 | 486 | EIYQAGSTPCNGVEGF | | 14 | 1671.7319 | RBD Abdala | 10 s (ice) | 3.588633 | 0.09908 | 4.482485 | 0.014746 | |
| 471 | 486 | EIYQAGSTPCNGVEGF | | 14 | 1671.7319 | RBD Abdala | 10 s (23 °C) | 5.32291 | 0.168853 | 4.502427 | 0.005092 | |
| 471 | 486 | EIYQAGSTPCNGVEGF | | 14 | 1671.7319 | RBD Abdala | 100 s (23 °C) | 6.941061 | 0.187583 | 4.513134 | 0.004689 | |
| 471 | 486 | EIYQAGSTPCNGVEGF | | 14 | 1671.7319 | RBD Abdala | 1,000 s (23 °C) | 7.399103 | 0.114274 | 4.500131 | 0.004344 | |
| 471 | 486 | EIYQAGSTPCNGVEGF | | 14 | 1671.7319 | RBD Abdala | 10,000 s (23 °C) | 7.617576 | 0.007179 | 4.48528 | 0.014227 | |
| 471 | 486 | EIYQAGSTPCNGVEGF | | 14 | 1671.7319 | RBD Abdala | 9 h (28 °C) | 7.674063 | 0.094835 | 4.491979 | 0.022734 | |
| 471 | 486 | EIYQAGSTPCNGVEGF | | 14 | 1671.7319 | MaxD | Max | 7.580706 | 0.053113 | 4.465164 | 0.001065 | 0.430022105 |
| 472 | 486 | IYQAGSTPCNGVEGF | | 13 | 1542.6893 | RBD HEK 293-F | 10 s (ice) | 3.626309 | 0.056428 | 4.298975 | 0.016717 | |
| 472 | 486 | IYQAGSTPCNGVEGF | | 13 | 1542.6893 | RBD HEK 293-F | 10 s (23 °C) | 5.875863 | 0.029348 | 4.295111 | 0.025453 | |
| 472 | 486 | IYQAGSTPCNGVEGF | | 13 | 1542.6893 | RBD HEK 293-F | 100 s (23 °C) | 7.323222 | 0.133225 | 4.305181 | 0.028752 | |
| 472 | 486 | IYQAGSTPCNGVEGF | | 13 | 1542.6893 | RBD HEK 293-F | 1,000 s (23 °C) | 7.380232 | 0.043198 | 4.298678 | 0.030653 | |
| 472 | 486 | IYQAGSTPCNGVEGF | | 13 | 1542.6893 | RBD HEK 293-F | 10,000 s (23 °C) | 7.334023 | 0.098373 | 4.302945 | 0.019402 | |
| 472 | 486 | IYQAGSTPCNGVEGF | | 13 | 1542.6893 | RBD HEK 293-F | 9 h (28 °C) | 7.295944 | 0.10562 | 4.287142 | 0.004778 | |
| 472 | 486 | IYQAGSTPCNGVEGF | | 13 | 1542.6893 | RBD Abdala | 10 s (ice) | 3.569438 | 0.041637 | 4.300971 | 0.020292 | |
| 472 | 486 | IYQAGSTPCNGVEGF | | 13 | 1542.6893 | RBD Abdala | 10 s (23 °C) | 5.431758 | 0.022645 | 4.32439 | 0.004448 | |
| 472 | 486 | IYQAGSTPCNGVEGF | | 13 | 1542.6893 | RBD Abdala | 100 s (23 °C) | 6.841541 | 0.134846 | 4.335335 | 0.002283 | |
| 472 | 486 | IYQAGSTPCNGVEGF | | 13 | 1542.6893 | RBD Abdala | 1,000 s (23 °C) | 6.971719 | 0.151194 | 4.324467 | 0.005536 | |

| | | | | | | | | | | | | |
|---|---|---|---|---|---|---|---|---|---|---|---|---|
| 472 | 486 | IYQAGSTPCNGVEGF | | 13 | 1542.6893 | RBD Abdala | 10,000 s (23 °C) | 7.189254 | 0.051147 | 4.302468 | 0.015873 | |
| 472 | 486 | IYQAGSTPCNGVEGF | | 13 | 1542.6893 | RBD Abdala | 9 h (28 °C) | 7.160975 | 0.084508 | 4.314373 | 0.026525 | |
| 472 | 486 | IYQAGSTPCNGVEGF | | 13 | 1542.6893 | MaxD | Max | 7.136469 | 0.082785 | 4.285384 | 0.001471 | 0.422148259 |
| 472 | 487 | IYQAGSTPCNGVEGFN | | 14 | 1656.7322 | RBD HEK 293-F | 10 s (ice) | 4.469162 | 0.133834 | 3.726359 | 0.018397 | |
| 472 | 487 | IYQAGSTPCNGVEGFN | | 14 | 1656.7322 | RBD HEK 293-F | 10 s (23 °C) | 6.881502 | 0.017585 | 3.730247 | 0.028942 | |
| 472 | 487 | IYQAGSTPCNGVEGFN | | 14 | 1656.7322 | RBD HEK 293-F | 100 s (23 °C) | 8.327331 | 0.137257 | 3.737523 | 0.036713 | |
| 472 | 487 | IYQAGSTPCNGVEGFN | | 14 | 1656.7322 | RBD HEK 293-F | 1,000 s (23 °C) | 8.419681 | 0.057634 | 3.731048 | 0.03369 | |
| 472 | 487 | IYQAGSTPCNGVEGFN | | 14 | 1656.7322 | RBD HEK 293-F | 10,000 s (23 °C) | 8.377857 | 0.144887 | 3.734252 | 0.024039 | |
| 472 | 487 | IYQAGSTPCNGVEGFN | | 14 | 1656.7322 | RBD HEK 293-F | 9 h (28 °C) | 8.291512 | 0.106439 | 3.719145 | 0.005331 | |
| 472 | 487 | IYQAGSTPCNGVEGFN | | 14 | 1656.7322 | RBD Abdala | 10 s (ice) | 4.426544 | 0.037465 | 3.727316 | 0.017936 | |
| 472 | 487 | IYQAGSTPCNGVEGFN | | 14 | 1656.7322 | RBD Abdala | 10 s (23 °C) | 6.579906 | 0.173386 | 3.757629 | 0.006945 | |
| 472 | 487 | IYQAGSTPCNGVEGFN | | 14 | 1656.7322 | RBD Abdala | 100 s (23 °C) | 7.8799 | 0.068978 | 3.76096 | 0.004007 | |
| 472 | 487 | IYQAGSTPCNGVEGFN | | 14 | 1656.7322 | RBD Abdala | 1,000 s (23 °C) | 7.978519 | 0.184622 | 3.75396 | 0.00491 | |
| 472 | 487 | IYQAGSTPCNGVEGFN | | 14 | 1656.7322 | RBD Abdala | 10,000 s (23 °C) | 8.359418 | 0.018932 | 3.727162 | 0.021117 | |
| 472 | 487 | IYQAGSTPCNGVEGFN | | 14 | 1656.7322 | RBD Abdala | 9 h (28 °C) | 8.353989 | 0.109475 | 3.741475 | 0.030183 | |
| 472 | 487 | IYQAGSTPCNGVEGFN | | 14 | 1656.7322 | MaxD | Max | 8.176603 | 0.04145 | 3.716311 | 0.001387 | 0.38521782 |
| 487 | 494 | NCYFPLQS | | 6 | 971.4291 | RBD HEK 293-F | 10 s (ice) | 0.533054 | 0.015083 | 4.649706 | 0.0057 | |
| 487 | 494 | NCYFPLQS | | 6 | 971.4291 | RBD HEK 293-F | 10 s (23 °C) | 1.280101 | 0.021868 | 4.647365 | 0.016724 | |
| 487 | 494 | NCYFPLQS | | 6 | 971.4291 | RBD HEK 293-F | 100 s (23 °C) | 1.783084 | 0.061641 | 4.654334 | 0.019274 | |
| 487 | 494 | NCYFPLQS | | 6 | 971.4291 | RBD HEK 293-F | 1,000 s (23 °C) | 2.485487 | 0.027412 | 4.652244 | 0.01936 | |
| 487 | 494 | NCYFPLQS | | 6 | 971.4291 | RBD HEK 293-F | 10,000 s (23 °C) | 2.937834 | 0.010391 | 4.651352 | 0.012114 | |
| 487 | 494 | NCYFPLQS | | 6 | 971.4291 | RBD HEK 293-F | 9 h (28 °C) | 3.517313 | 0.05493 | 4.639064 | 0.003066 | |
| 487 | 494 | NCYFPLQS | | 6 | 971.4291 | RBD Abdala | 10 s (ice) | 0.557291 | 0.089918 | 4.643331 | 0.006782 | |
| 487 | 494 | NCYFPLQS | | 6 | 971.4291 | RBD Abdala | 10 s (23 °C) | 1.252729 | 0.025963 | 4.663843 | 0.005991 | |
| 487 | 494 | NCYFPLQS | | 6 | 971.4291 | RBD Abdala | 100 s (23 °C) | 1.857994 | 0.009207 | 4.671905 | 0.005422 | |
| 487 | 494 | NCYFPLQS | | 6 | 971.4291 | RBD Abdala | 1,000 s (23 °C) | 2.512433 | 0.035936 | 4.671618 | 0.002255 | |

| | | | | | | | | | | | | |
|---|---|---|---|---|---|---|---|---|---|---|---|---|
| 487 | 494 | NCYFPLQS | | 6 | 971.4291 | RBD Abdala | 10,000 s (23 °C) | 3.018245 | 0.005289 | 4.653665 | 0.011959 | |
| 487 | 494 | NCYFPLQS | | 6 | 971.4291 | RBD Abdala | 9 h (28 °C) | 3.505377 | 0.028944 | 4.658721 | 0.0175 | |
| 487 | 494 | NCYFPLQS | | 6 | 971.4291 | MaxD | Max | 4.175145 | 0.076519 | 4.637385 | 0.002232 | 0.267518421 |
| 487 | 510 | NCYFPLQSYGFQPTNGVGYQPYRV | | 20 | 2798.3031 | RBD HEK 293-F | 10 s (ice) | 4.603099 | 0.156961 | 6.094067 | 0.006567 | |
| 487 | 510 | NCYFPLQSYGFQPTNGVGYQPYRV | | 20 | 2798.3031 | RBD HEK 293-F | 10 s (23 °C) | 6.227415 | 0.009447 | 6.080053 | 0.003841 | |
| 487 | 510 | NCYFPLQSYGFQPTNGVGYQPYRV | | 20 | 2798.3031 | RBD HEK 293-F | 100 s (23 °C) | 7.657719 | 0.093174 | 6.094479 | 0.005981 | |
| 487 | 510 | NCYFPLQSYGFQPTNGVGYQPYRV | | 20 | 2798.3031 | RBD HEK 293-F | 1,000 s (23 °C) | 9.058118 | 0.082108 | 6.10065 | 0.001596 | |
| 487 | 510 | NCYFPLQSYGFQPTNGVGYQPYRV | | 20 | 2798.3031 | RBD HEK 293-F | 10,000 s (23 °C) | 9.886552 | 0.018904 | 6.074435 | 0.006446 | |
| 487 | 510 | NCYFPLQSYGFQPTNGVGYQPYRV | | 20 | 2798.3031 | RBD HEK 293-F | 9 h (28 °C) | 10.516221 | 0.089789 | 6.071718 | 0.004396 | |
| 487 | 510 | NCYFPLQSYGFQPTNGVGYQPYRV | | 20 | 2798.3031 | RBD Abdala | 10 s (ice) | 4.607908 | 0.07685 | 6.091069 | 0.006461 | |
| 487 | 510 | NCYFPLQSYGFQPTNGVGYQPYRV | | 20 | 2798.3031 | RBD Abdala | 10 s (23 °C) | 5.947615 | 0 | 6.091908 | 0 | |
| 487 | 510 | NCYFPLQSYGFQPTNGVGYQPYRV | | 20 | 2798.3031 | RBD Abdala | 100 s (23 °C) | 7.430488 | 0.213234 | 6.091828 | 0.018228 | |
| 487 | 510 | NCYFPLQSYGFQPTNGVGYQPYRV | | 20 | 2798.3031 | RBD Abdala | 1,000 s (23 °C) | 9.026747 | 0.038784 | 6.103254 | 0.00109 | |
| 487 | 510 | NCYFPLQSYGFQPTNGVGYQPYRV | | 20 | 2798.3031 | RBD Abdala | 10,000 s (23 °C) | 10.171265 | 0.030898 | 6.087644 | 0.009788 | |
| 487 | 510 | NCYFPLQSYGFQPTNGVGYQPYRV | | 20 | 2798.3031 | RBD Abdala | 9 h (28 °C) | 10.734467 | 0.03552 | 6.093465 | 0.006017 | |
| 487 | 510 | NCYFPLQSYGFQPTNGVGYQPYRV | | 20 | 2798.3031 | MaxD | Max | 13.145657 | 0.129086 | 6.053571 | 0.024969 | 0.308123316 |
| 487 | 512 | NCYFPLQSYGFQPTNGVGYQPYRVVV | | 22 | 2996.44 | RBD HEK 293-F | 10 s (ice) | 4.749975 | 0.063235 | 6.393941 | 0.004426 | |
| 487 | 512 | NCYFPLQSYGFQPTNGVGYQPYRVVV | | 22 | 2996.44 | RBD HEK 293-F | 10 s (23 °C) | 6.608698 | 0.045313 | 6.381463 | 0.003475 | |
| 487 | 512 | NCYFPLQSYGFQPTNGVGYQPYRVVV | | 22 | 2996.44 | RBD HEK 293-F | 100 s (23 °C) | 8.084307 | 0.149383 | 6.397512 | 0.005974 | |
| 487 | 512 | NCYFPLQSYGFQPTNGVGYQPYRVVV | | 22 | 2996.44 | RBD HEK 293-F | 1,000 s (23 °C) | 9.443849 | 0.101719 | 6.404031 | 0.001494 | |
| 487 | 512 | NCYFPLQSYGFQPTNGVGYQPYRVVV | | 22 | 2996.44 | RBD HEK 293-F | 10,000 s (23 °C) | 10.523219 | 0.094718 | 6.37485 | 0.00345 | |
| 487 | 512 | NCYFPLQSYGFQPTNGVGYQPYRVVV | | 22 | 2996.44 | RBD HEK 293-F | 9 h (28 °C) | 11.178174 | 0.065788 | 6.371531 | 0.004322 | |
| 487 | 512 | NCYFPLQSYGFQPTNGVGYQPYRVVV | | 22 | 2996.44 | RBD Abdala | 10 s (ice) | 4.828917 | 0.100853 | 6.393789 | 0.008857 | |
| 487 | 512 | NCYFPLQSYGFQPTNGVGYQPYRVVV | | 22 | 2996.44 | RBD Abdala | 10 s (23 °C) | 6.35317 | 0 | 6.393256 | 0 | |
| 487 | 512 | NCYFPLQSYGFQPTNGVGYQPYRVVV | | 22 | 2996.44 | RBD Abdala | 100 s (23 °C) | 7.794348 | 0.093023 | 6.39277 | 0.018709 | |
| 487 | 512 | NCYFPLQSYGFQPTNGVGYQPYRVVV | | 22 | 2996.44 | RBD Abdala | 1,000 s (23 °C) | 9.376457 | 0.072657 | 6.405931 | 0.000121 | |

| | | | | | | | | | | | | |
|---|---|---|---|---|---|---|---|---|---|---|---|---|
| 487 | 512 | NCYFPLQSYGFQPTNGVGYQPYRVVV | | 22 | 2996.44 | RBD Abdala | 10,000 s (23 °C) | 10.612594 | 0.007109 | 6.390727 | 0.010732 | |
| 487 | 512 | NCYFPLQSYGFQPTNGVGYQPYRVVV | | 22 | 2996.44 | RBD Abdala | 9 h (28 °C) | 11.387322 | 0.074629 | 6.398141 | 0.004926 | |
| 487 | 512 | NCYFPLQSYGFQPTNGVGYQPYRVVV | | 22 | 2996.44 | MaxD | Max | 15.617597 | 0.119565 | 6.35491 | 0.01473 | 0.252746555 |
| 490 | 495 | FPLQSY | | 4 | 754.377 | RBD HEK 293-F | 10 s (ice) | 0.549389 | 0.007709 | 3.935083 | 0.015831 | |
| 490 | 495 | FPLQSY | | 4 | 754.377 | RBD HEK 293-F | 10 s (23 °C) | 0.924973 | 0.012283 | 3.938412 | 0.023419 | |
| 490 | 495 | FPLQSY | | 4 | 754.377 | RBD HEK 293-F | 100 s (23 °C) | 0.913937 | 0.003532 | 3.945574 | 0.031102 | |
| 490 | 495 | FPLQSY | | 4 | 754.377 | RBD HEK 293-F | 1,000 s (23 °C) | 1.123271 | 0.002373 | 3.93958 | 0.029508 | |
| 490 | 495 | FPLQSY | | 4 | 754.377 | RBD HEK 293-F | 10,000 s (23 °C) | 2.072762 | 0.006223 | 3.938251 | 0.018714 | |
| 490 | 495 | FPLQSY | | 4 | 754.377 | RBD HEK 293-F | 9 h (28 °C) | 2.690254 | 0.023563 | 3.922851 | 0.003552 | |
| 490 | 495 | FPLQSY | | 4 | 754.377 | RBD Abdala | 10 s (ice) | 0.54285 | 0.004964 | 3.941546 | 0.01904 | |
| 490 | 495 | FPLQSY | | 4 | 754.377 | RBD Abdala | 10 s (23 °C) | 0.881436 | 0.01083 | 3.966974 | 0.004609 | |
| 490 | 495 | FPLQSY | | 4 | 754.377 | RBD Abdala | 100 s (23 °C) | 0.911271 | 0.007462 | 3.976628 | 0.004003 | |
| 490 | 495 | FPLQSY | | 4 | 754.377 | RBD Abdala | 1,000 s (23 °C) | 1.166091 | 0.016106 | 3.969996 | 0.006489 | |
| 490 | 495 | FPLQSY | | 4 | 754.377 | RBD Abdala | 10,000 s (23 °C) | 2.158472 | 0.000609 | 3.941235 | 0.017687 | |
| 490 | 495 | FPLQSY | | 4 | 754.377 | RBD Abdala | 9 h (28 °C) | 2.771656 | 0.020505 | 3.950854 | 0.026615 | |
| 490 | 495 | FPLQSY | | 4 | 754.377 | MaxD | Max | 3.189959 | 0.026238 | 3.925077 | 0.004944 | 0.160537105 |
| 490 | 510 | FPLQSYGFQPTNGVGYQPYRV | | 17 | 2418.1877 | RBD HEK 293-F | 10 s (ice) | 5.684401 | 0.040076 | 5.31854 | 0.005112 | |
| 490 | 510 | FPLQSYGFQPTNGVGYQPYRV | | 17 | 2418.1877 | RBD HEK 293-F | 10 s (23 °C) | 7.265738 | 0.002394 | 5.306161 | 0.006558 | |
| 490 | 510 | FPLQSYGFQPTNGVGYQPYRV | | 17 | 2418.1877 | RBD HEK 293-F | 100 s (23 °C) | 8.387774 | 0.057454 | 5.316972 | 0.008458 | |
| 490 | 510 | FPLQSYGFQPTNGVGYQPYRV | | 17 | 2418.1877 | RBD HEK 293-F | 1,000 s (23 °C) | 9.319075 | 0.022916 | 5.324074 | 0.006985 | |
| 490 | 510 | FPLQSYGFQPTNGVGYQPYRV | | 17 | 2418.1877 | RBD HEK 293-F | 10,000 s (23 °C) | 10.40254 | 0.029724 | 5.306252 | 0.007379 | |
| 490 | 510 | FPLQSYGFQPTNGVGYQPYRV | | 17 | 2418.1877 | RBD HEK 293-F | 9 h (28 °C) | 11.148555 | 0.077727 | 5.305048 | 0.004469 | |
| 490 | 510 | FPLQSYGFQPTNGVGYQPYRV | | 17 | 2418.1877 | RBD Abdala | 10 s (ice) | 5.658576 | 0.036691 | 5.316295 | 0.00302 | |
| 490 | 510 | FPLQSYGFQPTNGVGYQPYRV | | 17 | 2418.1877 | RBD Abdala | 10 s (23 °C) | 7.266487 | 0.050712 | 5.322974 | 0.004538 | |
| 490 | 510 | FPLQSYGFQPTNGVGYQPYRV | | 17 | 2418.1877 | RBD Abdala | 100 s (23 °C) | 8.119354 | 0.201336 | 5.324308 | 0.005105 | |
| 490 | 510 | FPLQSYGFQPTNGVGYQPYRV | | 17 | 2418.1877 | RBD Abdala | 1,000 s (23 °C) | 9.28622 | 0.214179 | 5.325896 | 0.00224 | |

| | | | | | | | | | | | | |
|---|---|---|---|---|---|---|---|---|---|---|---|---|
| 490 | 510 | FPLQSYGFQPTNGVGYQPYRV | | 17 | 2418.1877 | RBD Abdala | 10,000 s (23 °C) | 10.59044 | 0.092361 | 5.319005 | 0.007129 | |
| 490 | 510 | FPLQSYGFQPTNGVGYQPYRV | | 17 | 2418.1877 | RBD Abdala | 9 h (28 °C) | 11.343988 | 0.050877 | 5.320251 | 0.009749 | |
| 490 | 510 | FPLQSYGFQPTNGVGYQPYRV | | 17 | 2418.1877 | MaxD | Max | 14.335161 | 0.07061 | 5.297219 | 0.004235 | 0.112373932 |
| 490 | 512 | FPLQSYGFQPTNGVGYQPYRVVV | | 19 | 2616.3245 | RBD HEK 293-F | 10 s (ice) | 5.334417 | 0.070983 | 5.719341 | 0.00402 | |
| 490 | 512 | FPLQSYGFQPTNGVGYQPYRVVV | | 19 | 2616.3245 | RBD HEK 293-F | 10 s (23 °C) | 6.937183 | 0.01167 | 5.708832 | 0.00544 | |
| 490 | 512 | FPLQSYGFQPTNGVGYQPYRVVV | | 19 | 2616.3245 | RBD HEK 293-F | 100 s (23 °C) | 7.966682 | 0.025674 | 5.722442 | 0.005023 | |
| 490 | 512 | FPLQSYGFQPTNGVGYQPYRVVV | | 19 | 2616.3245 | RBD HEK 293-F | 1,000 s (23 °C) | 8.949548 | 0.001434 | 5.72731 | 0.004955 | |
| 490 | 512 | FPLQSYGFQPTNGVGYQPYRVVV | | 19 | 2616.3245 | RBD HEK 293-F | 10,000 s (23 °C) | 9.909529 | 0.011495 | 5.704273 | 0.00417 | |
| 490 | 512 | FPLQSYGFQPTNGVGYQPYRVVV | | 19 | 2616.3245 | RBD HEK 293-F | 9 h (28 °C) | 10.761632 | 0.093204 | 5.698879 | 0.003129 | |
| 490 | 512 | FPLQSYGFQPTNGVGYQPYRVVV | | 19 | 2616.3245 | RBD Abdala | 10 s (ice) | 5.312578 | 0.05021 | 5.717576 | 0.00644 | |
| 490 | 512 | FPLQSYGFQPTNGVGYQPYRVVV | | 19 | 2616.3245 | RBD Abdala | 10 s (23 °C) | 6.875149 | 0.015368 | 5.723959 | 0.005173 | |
| 490 | 512 | FPLQSYGFQPTNGVGYQPYRVVV | | 19 | 2616.3245 | RBD Abdala | 100 s (23 °C) | 7.779637 | 0.163221 | 5.72061 | 0.009059 | |
| 490 | 512 | FPLQSYGFQPTNGVGYQPYRVVV | | 19 | 2616.3245 | RBD Abdala | 1,000 s (23 °C) | 8.914671 | 0.164901 | 5.728741 | 0.000624 | |
| 490 | 512 | FPLQSYGFQPTNGVGYQPYRVVV | | 19 | 2616.3245 | RBD Abdala | 10,000 s (23 °C) | 10.057601 | 0.04538 | 5.716485 | 0.011269 | |
| 490 | 512 | FPLQSYGFQPTNGVGYQPYRVVV | | 19 | 2616.3245 | RBD Abdala | 9 h (28 °C) | 11.0002 | 0.049477 | 5.722192 | 0.006154 | |
| 490 | 512 | FPLQSYGFQPTNGVGYQPYRVVV | | 19 | 2616.3245 | MaxD | Max | 15.927732 | 0.119774 | 5.676476 | 0.028507 | 0.117577175 |
| 490 | 513 | FPLQSYGFQPTNGVGYQPYRVVVL | | 20 | 2729.4086 | RBD HEK 293-F | 10 s (ice) | 5.220669 | 0.093399 | 6.187915 | 0.004974 | |
| 490 | 513 | FPLQSYGFQPTNGVGYQPYRVVVL | | 20 | 2729.4086 | RBD HEK 293-F | 10 s (23 °C) | 6.804555 | 0.018725 | 6.174015 | 0.003367 | |
| 490 | 513 | FPLQSYGFQPTNGVGYQPYRVVVL | | 20 | 2729.4086 | RBD HEK 293-F | 100 s (23 °C) | 7.786516 | 0.054657 | 6.190385 | 0.006945 | |
| 490 | 513 | FPLQSYGFQPTNGVGYQPYRVVVL | | 20 | 2729.4086 | RBD HEK 293-F | 1,000 s (23 °C) | 8.80792 | 0.037706 | 6.19562 | 0.002717 | |
| 490 | 513 | FPLQSYGFQPTNGVGYQPYRVVVL | | 20 | 2729.4086 | RBD HEK 293-F | 10,000 s (23 °C) | 9.69583 | 3.73833E-05 | 6.169407 | 0.006081 | |
| 490 | 513 | FPLQSYGFQPTNGVGYQPYRVVVL | | 20 | 2729.4086 | RBD HEK 293-F | 9 h (28 °C) | 10.541438 | 0.071804 | 6.16767 | 0.005497 | |
| 490 | 513 | FPLQSYGFQPTNGVGYQPYRVVVL | | 20 | 2729.4086 | RBD Abdala | 10 s (ice) | 5.266394 | 0.052512 | 6.184196 | 0.009841 | |
| 490 | 513 | FPLQSYGFQPTNGVGYQPYRVVVL | | 20 | 2729.4086 | RBD Abdala | 10 s (23 °C) | 6.671117 | 0.100427 | 6.19441 | 0.007011 | |
| 490 | 513 | FPLQSYGFQPTNGVGYQPYRVVVL | | 20 | 2729.4086 | RBD Abdala | 100 s (23 °C) | 7.590833 | 0.185951 | 6.195313 | 0.018042 | |
| 490 | 513 | FPLQSYGFQPTNGVGYQPYRVVVL | | 20 | 2729.4086 | RBD Abdala | 1,000 s (23 °C) | 8.707223 | 0.216425 | 6.199959 | 0.00038 | |

| | | | | | | | | | | | | |
|---|---|---|---|---|---|---|---|---|---|---|---|---|
| 490 | 513 | FPLQSYGFQPTNGVGYQPYRVVVL | | 20 | 2729.4086 | RBD Abdala | 10,000 s (23 °C) | 9.783986 | 0.004463 | 6.18721 | 0.011957 | |
| 490 | 513 | FPLQSYGFQPTNGVGYQPYRVVVL | | 20 | 2729.4086 | RBD Abdala | 9 h (28 °C) | 10.700506 | 0.065441 | 6.190741 | 0.005221 | |
| 490 | 513 | FPLQSYGFQPTNGVGYQPYRVVVL | | 20 | 2729.4086 | MaxD | Max | 16.055983 | 0.070213 | 6.146472 | 0.020968 | 0.154948263 |
| 491 | 512 | PLQSYGFQPTNGVGYQPYRVVV | | 19 | 2469.2561 | RBD HEK 293-F | 10 s (ice) | 4.434958 | 0.093894 | 5.145593 | 0.005872 | |
| 491 | 512 | PLQSYGFQPTNGVGYQPYRVVV | | 19 | 2469.2561 | RBD HEK 293-F | 10 s (23 °C) | 5.802841 | 0.000618 | 5.134405 | 0.01094 | |
| 491 | 512 | PLQSYGFQPTNGVGYQPYRVVV | | 19 | 2469.2561 | RBD HEK 293-F | 100 s (23 °C) | 6.696267 | 0.149037 | 5.147266 | 0.010289 | |
| 491 | 512 | PLQSYGFQPTNGVGYQPYRVVV | | 19 | 2469.2561 | RBD HEK 293-F | 1,000 s (23 °C) | 7.554865 | 0.036796 | 5.146955 | 0.011091 | |
| 491 | 512 | PLQSYGFQPTNGVGYQPYRVVV | | 19 | 2469.2561 | RBD HEK 293-F | 10,000 s (23 °C) | 8.032869 | 0.085645 | 5.138104 | 0.008165 | |
| 491 | 512 | PLQSYGFQPTNGVGYQPYRVVV | | 19 | 2469.2561 | RBD HEK 293-F | 9 h (28 °C) | 8.397248 | 0.106227 | 5.137726 | 0.005083 | |
| 491 | 512 | PLQSYGFQPTNGVGYQPYRVVV | | 19 | 2469.2561 | RBD Abdala | 10 s (ice) | 4.397509 | 0.070242 | 5.146332 | 0.005192 | |
| 491 | 512 | PLQSYGFQPTNGVGYQPYRVVV | | 19 | 2469.2561 | RBD Abdala | 10 s (23 °C) | 5.63884 | 0.149453 | 5.154066 | 0.006604 | |
| 491 | 512 | PLQSYGFQPTNGVGYQPYRVVV | | 19 | 2469.2561 | RBD Abdala | 100 s (23 °C) | 6.495039 | 0.136499 | 5.156268 | 0.002748 | |
| 491 | 512 | PLQSYGFQPTNGVGYQPYRVVV | | 19 | 2469.2561 | RBD Abdala | 1,000 s (23 °C) | 7.453846 | 0.077466 | 5.150646 | 0.004469 | |
| 491 | 512 | PLQSYGFQPTNGVGYQPYRVVV | | 19 | 2469.2561 | RBD Abdala | 10,000 s (23 °C) | 8.158926 | 0.015349 | 5.15108 | 0.003639 | |
| 491 | 512 | PLQSYGFQPTNGVGYQPYRVVV | | 19 | 2469.2561 | RBD Abdala | 9 h (28 °C) | 8.484751 | 0.069273 | 5.150266 | 0.011018 | |
| 491 | 512 | PLQSYGFQPTNGVGYQPYRVVV | | 19 | 2469.2561 | MaxD | Max | 12.534978 | 0.060762 | 5.133996 | 0.003828 | 0.305541385 |
| 491 | 513 | PLQSYGFQPTNGVGYQPYRVVVL | | 20 | 2582.3402 | RBD HEK 293-F | 10 s (ice) | 4.807391 | 0.162705 | 5.713447 | 0.005275 | |
| 491 | 513 | PLQSYGFQPTNGVGYQPYRVVVL | | 20 | 2582.3402 | RBD HEK 293-F | 10 s (23 °C) | 6.359457 | 0.005372 | 5.701645 | 0.006476 | |
| 491 | 513 | PLQSYGFQPTNGVGYQPYRVVVL | | 20 | 2582.3402 | RBD HEK 293-F | 100 s (23 °C) | 7.388907 | 0.004919 | 5.714639 | 0.005367 | |
| 491 | 513 | PLQSYGFQPTNGVGYQPYRVVVL | | 20 | 2582.3402 | RBD HEK 293-F | 1,000 s (23 °C) | 8.300533 | 0.000423 | 5.719362 | 0.00624 | |
| 491 | 513 | PLQSYGFQPTNGVGYQPYRVVVL | | 20 | 2582.3402 | RBD HEK 293-F | 10,000 s (23 °C) | 8.710214 | 0.000816 | 5.697756 | 0.00743 | |
| 491 | 513 | PLQSYGFQPTNGVGYQPYRVVVL | | 20 | 2582.3402 | RBD HEK 293-F | 9 h (28 °C) | 9.006365 | 0.1902 | 5.696308 | 0.003651 | |
| 491 | 513 | PLQSYGFQPTNGVGYQPYRVVVL | | 20 | 2582.3402 | RBD Abdala | 10 s (ice) | 4.993463 | 0.088079 | 5.709967 | 0.007877 | |
| 491 | 513 | PLQSYGFQPTNGVGYQPYRVVVL | | 20 | 2582.3402 | RBD Abdala | 10 s (23 °C) | 6.403712 | 0 | 5.712888 | 0 | |
| 491 | 513 | PLQSYGFQPTNGVGYQPYRVVVL | | 20 | 2582.3402 | RBD Abdala | 100 s (23 °C) | 7.125632 | 0.177683 | 5.710239 | 0.009173 | |
| 491 | 513 | PLQSYGFQPTNGVGYQPYRVVVL | | 20 | 2582.3402 | RBD Abdala | 1,000 s (23 °C) | 8.050363 | 0.197236 | 5.723071 | 0.000592 | |

| | | | | | | | | | | | | |
|---|---|---|---|---|---|---|---|---|---|---|---|---|
| 491 | 513 | PLQSYGFQPTNGVGYQPYRVVVL | | 20 | 2582.3402 | RBD Abdala | 10,000 s (23 °C) | 8.84444 | 0.074894 | 5.715117 | 0.009254 | |
| 491 | 513 | PLQSYGFQPTNGVGYQPYRVVVL | | 20 | 2582.3402 | RBD Abdala | 9 h (28 °C) | 9.176976 | 0.043799 | 5.720098 | 0.007785 | |
| 491 | 513 | PLQSYGFQPTNGVGYQPYRVVVL | | 20 | 2582.3402 | MaxD | Max | 14.072174 | 0.065119 | 5.668708 | 0.024277 | 0.259359263 |
| 495 | 510 | YGFQPTNGVGYQPYRV | | 13 | 1845.8919 | RBD HEK 293-F | 10 s (ice) | 4.392166 | 0.028459 | 4.465223 | 0.016641 | |
| 495 | 510 | YGFQPTNGVGYQPYRV | | 13 | 1845.8919 | RBD HEK 293-F | 10 s (23 °C) | 5.708055 | 0.005945 | 4.455829 | 0.023811 | |
| 495 | 510 | YGFQPTNGVGYQPYRV | | 13 | 1845.8919 | RBD HEK 293-F | 100 s (23 °C) | 6.785023 | 0.025385 | 4.467945 | 0.024909 | |
| 495 | 510 | YGFQPTNGVGYQPYRV | | 13 | 1845.8919 | RBD HEK 293-F | 1,000 s (23 °C) | 7.59401 | 0.036997 | 4.463257 | 0.027388 | |
| 495 | 510 | YGFQPTNGVGYQPYRV | | 13 | 1845.8919 | RBD HEK 293-F | 10,000 s (23 °C) | 7.543737 | 0.016617 | 4.464497 | 0.015655 | |
| 495 | 510 | YGFQPTNGVGYQPYRV | | 13 | 1845.8919 | RBD HEK 293-F | 9 h (28 °C) | 7.583733 | 0.029825 | 4.453323 | 0.004808 | |
| 495 | 510 | YGFQPTNGVGYQPYRV | | 13 | 1845.8919 | RBD Abdala | 10 s (ice) | 4.420141 | 0.050776 | 4.467474 | 0.017858 | |
| 495 | 510 | YGFQPTNGVGYQPYRV | | 13 | 1845.8919 | RBD Abdala | 10 s (23 °C) | 5.690157 | 0.020313 | 4.488429 | 0.002492 | |
| 495 | 510 | YGFQPTNGVGYQPYRV | | 13 | 1845.8919 | RBD Abdala | 100 s (23 °C) | 6.553268 | 0.216314 | 4.500131 | 0.004374 | |
| 495 | 510 | YGFQPTNGVGYQPYRV | | 13 | 1845.8919 | RBD Abdala | 1,000 s (23 °C) | 7.417916 | 0.193281 | 4.491118 | 0.004164 | |
| 495 | 510 | YGFQPTNGVGYQPYRV | | 13 | 1845.8919 | RBD Abdala | 10,000 s (23 °C) | 7.623771 | 0.020294 | 4.471273 | 0.016201 | |
| 495 | 510 | YGFQPTNGVGYQPYRV | | 13 | 1845.8919 | RBD Abdala | 9 h (28 °C) | 7.69328 | 0.02972 | 4.479722 | 0.02427 | |
| 495 | 510 | YGFQPTNGVGYQPYRV | | 13 | 1845.8919 | MaxD | Max | 9.94498 | 0.028896 | 4.452413 | 0.003859 | 0.194738462 |
| 495 | 512 | YGFQPTNGVGYQPYRVVV | | 15 | 2044.0287 | RBD HEK 293-F | 10 s (ice) | 4.471566 | 0.036657 | 4.954451 | 0.009247 | |
| 495 | 512 | YGFQPTNGVGYQPYRVVV | | 15 | 2044.0287 | RBD HEK 293-F | 10 s (23 °C) | 5.73218 | 0.010227 | 4.942419 | 0.013233 | |
| 495 | 512 | YGFQPTNGVGYQPYRVVV | | 15 | 2044.0287 | RBD HEK 293-F | 100 s (23 °C) | 6.665751 | 0.023166 | 4.948825 | 0.017441 | |
| 495 | 512 | YGFQPTNGVGYQPYRVVV | | 15 | 2044.0287 | RBD HEK 293-F | 1,000 s (23 °C) | 7.495771 | 0.001691 | 4.951731 | 0.013059 | |
| 495 | 512 | YGFQPTNGVGYQPYRVVV | | 15 | 2044.0287 | RBD HEK 293-F | 10,000 s (23 °C) | 7.548473 | 0.007049 | 4.949311 | 0.011516 | |
| 495 | 512 | YGFQPTNGVGYQPYRVVV | | 15 | 2044.0287 | RBD HEK 293-F | 9 h (28 °C) | 7.630433 | 0.014245 | 4.948607 | 0.006085 | |
| 495 | 512 | YGFQPTNGVGYQPYRVVV | | 15 | 2044.0287 | RBD Abdala | 10 s (ice) | 4.505787 | 0.020026 | 4.956413 | 0.00321 | |
| 495 | 512 | YGFQPTNGVGYQPYRVVV | | 15 | 2044.0287 | RBD Abdala | 10 s (23 °C) | 5.657688 | 0.041373 | 4.965489 | 0.006238 | |
| 495 | 512 | YGFQPTNGVGYQPYRVVV | | 15 | 2044.0287 | RBD Abdala | 100 s (23 °C) | 6.612249 | 0.153843 | 4.971707 | 0.001446 | |
| 495 | 512 | YGFQPTNGVGYQPYRVVV | | 15 | 2044.0287 | RBD Abdala | 1,000 s (23 °C) | 7.414362 | 0.195223 | 4.961619 | 0.004093 | |

| | | | | | | | | | | | | |
|---|---|---|---|---|---|---|---|---|---|---|---|---|
| 495 | 512 | YGFQPTNGVGYQPYRVVV | | 15 | 2044.0287 | RBD Abdala | 10,000 s (23 °C) | 7.63085 | 0.077939 | 4.96092 | 0.000536 | |
| 495 | 512 | YGFQPTNGVGYQPYRVVV | | 15 | 2044.0287 | RBD Abdala | 9 h (28 °C) | 7.698646 | 0.036352 | 4.957498 | 0.01027 | |
| 495 | 512 | YGFQPTNGVGYQPYRVVV | | 15 | 2044.0287 | MaxD | Max | 11.872114 | 0.039739 | 4.951275 | 0.005914 | 0.166869193 |
| 495 | 513 | YGFQPTNGVGYQPYRVVVL | | 16 | 2157.1128 | RBD HEK 293-F | 10 s (ice) | 4.453679 | 0.049259 | 5.589312 | 0.005284 | |
| 495 | 513 | YGFQPTNGVGYQPYRVVVL | | 16 | 2157.1128 | RBD HEK 293-F | 10 s (23 °C) | 5.688235 | 0.014829 | 5.578684 | 0.005587 | |
| 495 | 513 | YGFQPTNGVGYQPYRVVVL | | 16 | 2157.1128 | RBD HEK 293-F | 100 s (23 °C) | 6.751592 | 0.010842 | 5.589301 | 0.00566 | |
| 495 | 513 | YGFQPTNGVGYQPYRVVVL | | 16 | 2157.1128 | RBD HEK 293-F | 1,000 s (23 °C) | 7.626781 | 0.002293 | 5.594429 | 0.006879 | |
| 495 | 513 | YGFQPTNGVGYQPYRVVVL | | 16 | 2157.1128 | RBD HEK 293-F | 10,000 s (23 °C) | 7.60745 | 0.001978 | 5.576892 | 0.005442 | |
| 495 | 513 | YGFQPTNGVGYQPYRVVVL | | 16 | 2157.1128 | RBD HEK 293-F | 9 h (28 °C) | 7.669443 | 0.049148 | 5.576152 | 0.002478 | |
| 495 | 513 | YGFQPTNGVGYQPYRVVVL | | 16 | 2157.1128 | RBD Abdala | 10 s (ice) | 4.483163 | 0.042922 | 5.585715 | 0.00541 | |
| 495 | 513 | YGFQPTNGVGYQPYRVVVL | | 16 | 2157.1128 | RBD Abdala | 10 s (23 °C) | 5.670421 | 0.031366 | 5.594778 | 0.007698 | |
| 495 | 513 | YGFQPTNGVGYQPYRVVVL | | 16 | 2157.1128 | RBD Abdala | 100 s (23 °C) | 6.605182 | 0.197991 | 5.591391 | 0.006631 | |
| 495 | 513 | YGFQPTNGVGYQPYRVVVL | | 16 | 2157.1128 | RBD Abdala | 1,000 s (23 °C) | 7.49313 | 0.233738 | 5.596947 | 0.000321 | |
| 495 | 513 | YGFQPTNGVGYQPYRVVVL | | 16 | 2157.1128 | RBD Abdala | 10,000 s (23 °C) | 7.680161 | 0.048354 | 5.588001 | 0.008534 | |
| 495 | 513 | YGFQPTNGVGYQPYRVVVL | | 16 | 2157.1128 | RBD Abdala | 9 h (28 °C) | 7.760204 | 0.041118 | 5.59305 | 0.007267 | |
| 495 | 513 | YGFQPTNGVGYQPYRVVVL | | 16 | 2157.1128 | MaxD | Max | 12.76944 | 0.066027 | 5.565621 | 0.009681 | 0.159905263 |
| 496 | 510 | GFQPTNGVGYQPYRV | | 12 | 1682.8285 | RBD HEK 293-F | 10 s (ice) | 4.104302 | 0.022496 | 3.912745 | 0.022356 | |
| 496 | 510 | GFQPTNGVGYQPYRV | | 12 | 1682.8285 | RBD HEK 293-F | 10 s (23 °C) | 4.997939 | 0.018785 | 3.910867 | 0.027722 | |
| 496 | 510 | GFQPTNGVGYQPYRV | | 12 | 1682.8285 | RBD HEK 293-F | 100 s (23 °C) | 6.079531 | 0.032388 | 3.920931 | 0.034477 | |
| 496 | 510 | GFQPTNGVGYQPYRV | | 12 | 1682.8285 | RBD HEK 293-F | 1,000 s (23 °C) | 6.896718 | 0.008744 | 3.915757 | 0.035033 | |
| 496 | 510 | GFQPTNGVGYQPYRV | | 12 | 1682.8285 | RBD HEK 293-F | 10,000 s (23 °C) | 6.894011 | 0.006502 | 3.917522 | 0.022298 | |
| 496 | 510 | GFQPTNGVGYQPYRV | | 12 | 1682.8285 | RBD HEK 293-F | 9 h (28 °C) | 6.954443 | 0.031025 | 3.901745 | 0.004395 | |
| 496 | 510 | GFQPTNGVGYQPYRV | | 12 | 1682.8285 | RBD Abdala | 10 s (ice) | 4.101627 | 0.02194 | 3.921388 | 0.021622 | |
| 496 | 510 | GFQPTNGVGYQPYRV | | 12 | 1682.8285 | RBD Abdala | 10 s (23 °C) | 4.994382 | 0.007474 | 3.949363 | 0.003173 | |
| 496 | 510 | GFQPTNGVGYQPYRV | | 12 | 1682.8285 | RBD Abdala | 100 s (23 °C) | 5.932491 | 0.17028 | 3.960182 | 0.001151 | |
| 496 | 510 | GFQPTNGVGYQPYRV | | 12 | 1682.8285 | RBD Abdala | 1,000 s (23 °C) | 6.790041 | 0.179684 | 3.953085 | 0.004359 | |

| | | | | | | | | | | | | |
|---|---|---|---|---|---|---|---|---|---|---|---|---|
| 496 | 510 | GFQPTNGVGYQPYRV | | 12 | 1682.8285 | RBD Abdala | 10,000 s (23 °C) | 6.953661 | 0.017696 | 3.928339 | 0.018831 | |
| 496 | 510 | GFQPTNGVGYQPYRV | | 12 | 1682.8285 | RBD Abdala | 9 h (28 °C) | 7.016181 | 0.026413 | 3.936103 | 0.02908 | |
| 496 | 510 | GFQPTNGVGYQPYRV | | 12 | 1682.8285 | MaxD | Max | 9.324576 | 0.029393 | 3.906217 | 0.003439 | 0.182054737 |
| 496 | 512 | GFQPTNGVGYQPYRVVV | | 14 | 1880.9654 | RBD HEK 293-F | 10 s (ice) | 4.052934 | 0.038519 | 4.540077 | 0.012797 | |
| 496 | 512 | GFQPTNGVGYQPYRVVV | | 14 | 1880.9654 | RBD HEK 293-F | 10 s (23 °C) | 4.900758 | 0.010906 | 4.530001 | 0.023409 | |
| 496 | 512 | GFQPTNGVGYQPYRVVV | | 14 | 1880.9654 | RBD HEK 293-F | 100 s (23 °C) | 5.93146 | 0.025024 | 4.537959 | 0.025558 | |
| 496 | 512 | GFQPTNGVGYQPYRVVV | | 14 | 1880.9654 | RBD HEK 293-F | 1,000 s (23 °C) | 6.752283 | 0.006473 | 4.539916 | 0.025445 | |
| 496 | 512 | GFQPTNGVGYQPYRVVV | | 14 | 1880.9654 | RBD HEK 293-F | 10,000 s (23 °C) | 6.769036 | 0.017572 | 4.540603 | 0.01547 | |
| 496 | 512 | GFQPTNGVGYQPYRVVV | | 14 | 1880.9654 | RBD HEK 293-F | 9 h (28 °C) | 6.819156 | 0.011304 | 4.530216 | 0.009478 | |
| 496 | 512 | GFQPTNGVGYQPYRVVV | | 14 | 1880.9654 | RBD Abdala | 10 s (ice) | 4.058296 | 0.047156 | 4.541419 | 0.01558 | |
| 496 | 512 | GFQPTNGVGYQPYRVVV | | 14 | 1880.9654 | RBD Abdala | 10 s (23 °C) | 4.874093 | 0.006431 | 4.560612 | 0.004031 | |
| 496 | 512 | GFQPTNGVGYQPYRVVV | | 14 | 1880.9654 | RBD Abdala | 100 s (23 °C) | 5.799294 | 0.187288 | 4.569515 | 0.001638 | |
| 496 | 512 | GFQPTNGVGYQPYRVVV | | 14 | 1880.9654 | RBD Abdala | 1,000 s (23 °C) | 6.641583 | 0.162123 | 4.56529 | 0.0039 | |
| 496 | 512 | GFQPTNGVGYQPYRVVV | | 14 | 1880.9654 | RBD Abdala | 10,000 s (23 °C) | 6.835203 | 0.031957 | 4.547965 | 0.01359 | |
| 496 | 512 | GFQPTNGVGYQPYRVVV | | 14 | 1880.9654 | RBD Abdala | 9 h (28 °C) | 6.914369 | 0.035271 | 4.55394 | 0.019457 | |
| 496 | 512 | GFQPTNGVGYQPYRVVV | | 14 | 1880.9654 | MaxD | Max | 11.007239 | 0.035741 | 4.521267 | 0.004529 | 0.172388045 |
| 496 | 513 | GFQPTNGVGYQPYRVVVL | | 15 | 1994.0494 | RBD HEK 293-F | 10 s (ice) | 4.021538 | 0.034664 | 5.159995 | 0.007592 | |
| 496 | 513 | GFQPTNGVGYQPYRVVVL | | 15 | 1994.0494 | RBD HEK 293-F | 10 s (23 °C) | 4.856399 | 0.000455 | 5.14909 | 0.010501 | |
| 496 | 513 | GFQPTNGVGYQPYRVVVL | | 15 | 1994.0494 | RBD HEK 293-F | 100 s (23 °C) | 5.947354 | 0.010338 | 5.156053 | 0.012762 | |
| 496 | 513 | GFQPTNGVGYQPYRVVVL | | 15 | 1994.0494 | RBD HEK 293-F | 1,000 s (23 °C) | 6.771993 | 0.014096 | 5.161725 | 0.011762 | |
| 496 | 513 | GFQPTNGVGYQPYRVVVL | | 15 | 1994.0494 | RBD HEK 293-F | 10,000 s (23 °C) | 6.765794 | 0.00289 | 5.151752 | 0.008534 | |
| 496 | 513 | GFQPTNGVGYQPYRVVVL | | 15 | 1994.0494 | RBD HEK 293-F | 9 h (28 °C) | 6.841788 | 0.022334 | 5.150608 | 0.005866 | |
| 496 | 513 | GFQPTNGVGYQPYRVVVL | | 15 | 1994.0494 | RBD Abdala | 10 s (ice) | 4.053679 | 0.028107 | 5.161136 | 0.003499 | |
| 496 | 513 | GFQPTNGVGYQPYRVVVL | | 15 | 1994.0494 | RBD Abdala | 10 s (23 °C) | 4.848924 | 0.024236 | 5.171672 | 0.005774 | |
| 496 | 513 | GFQPTNGVGYQPYRVVVL | | 15 | 1994.0494 | RBD Abdala | 100 s (23 °C) | 5.816279 | 0.14609 | 5.172311 | 0.003965 | |
| 496 | 513 | GFQPTNGVGYQPYRVVVL | | 15 | 1994.0494 | RBD Abdala | 1,000 s (23 °C) | 6.656072 | 0.186982 | 5.169516 | 0.003784 | |

| | | | | | | | | | | | | |
|---|---|---|---|---|---|---|---|---|---|---|---|---|
| 496 | 513 | GFQPTNGVGYQPYRVVVL | | 15 | 1994.0494 | RBD Abdala | 10,000 s (23 °C) | 6.852259 | 0.052391 | 5.167123 | 0.00402 | |
| 496 | 513 | GFQPTNGVGYQPYRVVVL | | 15 | 1994.0494 | RBD Abdala | 9 h (28 °C) | 6.925465 | 0.032534 | 5.165861 | 0.010085 | |
| 496 | 513 | GFQPTNGVGYQPYRVVVL | | 15 | 1994.0494 | MaxD | Max | 11.929518 | 0.032547 | 5.150194 | 0.004584 | 0.162840842 |
| 496 | 514 | GFQPTNGVGYQPYRVVVLS | | 16 | 2081.0815 | RBD HEK 293-F | 10 s (ice) | 4.077139 | 0.024351 | 4.68117 | 0.009041 | |
| 496 | 514 | GFQPTNGVGYQPYRVVVLS | | 16 | 2081.0815 | RBD HEK 293-F | 10 s (23 °C) | 4.929618 | 0.024225 | 4.67311 | 0.014064 | |
| 496 | 514 | GFQPTNGVGYQPYRVVVLS | | 16 | 2081.0815 | RBD HEK 293-F | 100 s (23 °C) | 5.989774 | 0.040008 | 4.67948 | 0.016429 | |
| 496 | 514 | GFQPTNGVGYQPYRVVVLS | | 16 | 2081.0815 | RBD HEK 293-F | 1,000 s (23 °C) | 6.850986 | 0.019092 | 4.679503 | 0.013429 | |
| 496 | 514 | GFQPTNGVGYQPYRVVVLS | | 16 | 2081.0815 | RBD HEK 293-F | 10,000 s (23 °C) | 6.7694 | 0.033966 | 4.682547 | 0.011312 | |
| 496 | 514 | GFQPTNGVGYQPYRVVVLS | | 16 | 2081.0815 | RBD HEK 293-F | 9 h (28 °C) | 6.934781 | 0.080774 | 4.676695 | 0.006747 | |
| 496 | 514 | GFQPTNGVGYQPYRVVVLS | | 16 | 2081.0815 | RBD Abdala | 10 s (ice) | 4.086676 | 0.035725 | 4.682443 | 0.005429 | |
| 496 | 514 | GFQPTNGVGYQPYRVVVLS | | 16 | 2081.0815 | RBD Abdala | 10 s (23 °C) | 4.865392 | 0.059957 | 4.695621 | 0.006254 | |
| 496 | 514 | GFQPTNGVGYQPYRVVVLS | | 16 | 2081.0815 | RBD Abdala | 100 s (23 °C) | 5.859394 | 0.141804 | 4.703138 | 0.000544 | |
| 496 | 514 | GFQPTNGVGYQPYRVVVLS | | 16 | 2081.0815 | RBD Abdala | 1,000 s (23 °C) | 6.761652 | 0.140032 | 4.69765 | 0.004243 | |
| 496 | 514 | GFQPTNGVGYQPYRVVVLS | | 16 | 2081.0815 | RBD Abdala | 10,000 s (23 °C) | 6.969013 | 0.039654 | 4.689303 | 0.003262 | |
| 496 | 514 | GFQPTNGVGYQPYRVVVLS | | 16 | 2081.0815 | RBD Abdala | 9 h (28 °C) | 6.972717 | 0.062352 | 4.6891 | 0.014053 | |
| 496 | 514 | GFQPTNGVGYQPYRVVVLS | | 16 | 2081.0815 | MaxD | Max | 12.956386 | 0.01111 | 4.668169 | 0.006595 | 0.147606184 |
| 497 | 512 | FQPTNGVGYQPYRVVV | | 13 | 1823.9439 | RBD HEK 293-F | 10 s (ice) | 3.945236 | 0.024453 | 4.373683 | 0.017949 | |
| 497 | 512 | FQPTNGVGYQPYRVVV | | 13 | 1823.9439 | RBD HEK 293-F | 10 s (23 °C) | 4.391969 | 0.013098 | 4.365981 | 0.024086 | |
| 497 | 512 | FQPTNGVGYQPYRVVV | | 13 | 1823.9439 | RBD HEK 293-F | 100 s (23 °C) | 5.128937 | 0.008155 | 4.38022 | 0.027409 | |
| 497 | 512 | FQPTNGVGYQPYRVVV | | 13 | 1823.9439 | RBD HEK 293-F | 1,000 s (23 °C) | 6.01864 | 0.038581 | 4.375621 | 0.028437 | |
| 497 | 512 | FQPTNGVGYQPYRVVV | | 13 | 1823.9439 | RBD HEK 293-F | 10,000 s (23 °C) | 5.986917 | 0.011849 | 4.3766 | 0.017467 | |
| 497 | 512 | FQPTNGVGYQPYRVVV | | 13 | 1823.9439 | RBD HEK 293-F | 9 h (28 °C) | 6.019524 | 0.01741 | 4.365662 | 0.00593 | |
| 497 | 512 | FQPTNGVGYQPYRVVV | | 13 | 1823.9439 | RBD Abdala | 10 s (ice) | 3.950855 | 0.033821 | 4.379684 | 0.019041 | |
| 497 | 512 | FQPTNGVGYQPYRVVV | | 13 | 1823.9439 | RBD Abdala | 10 s (23 °C) | 4.346048 | 0.043346 | 4.402415 | 0.005865 | |
| 497 | 512 | FQPTNGVGYQPYRVVV | | 13 | 1823.9439 | RBD Abdala | 100 s (23 °C) | 5.058706 | 0.107953 | 4.417116 | 0.001579 | |
| 497 | 512 | FQPTNGVGYQPYRVVV | | 13 | 1823.9439 | RBD Abdala | 1,000 s (23 °C) | 5.870493 | 0.170295 | 4.407223 | 0.006282 | |

| | | | | | | | | | | | | |
|---|---|---|---|---|---|---|---|---|---|---|---|---|
| 497 | 512 | FQPTNGVGYQPYRVVV | | 13 | 1823.9439 | RBD Abdala | 10,000 s (23 °C) | 6.033313 | 0.027513 | 4.383802 | 0.014914 | |
| 497 | 512 | FQPTNGVGYQPYRVVV | | 13 | 1823.9439 | RBD Abdala | 9 h (28 °C) | 6.107908 | 0.02122 | 4.394106 | 0.025387 | |
| 497 | 512 | FQPTNGVGYQPYRVVV | | 13 | 1823.9439 | MaxD | Max | 10.304573 | 0.05708 | 4.360259 | 0.003809 | 0.165621619 |
| 503 | 512 | VGYQPYRVVV | | 8 | 1179.6521 | RBD HEK 293-F | 10 s (ice) | 0.247551 | 0.026487 | 3.919049 | 0.018834 | |
| 503 | 512 | VGYQPYRVVV | | 8 | 1179.6521 | RBD HEK 293-F | 10 s (23 °C) | 0.437218 | 0.006992 | 3.919542 | 0.026996 | |
| 503 | 512 | VGYQPYRVVV | | 8 | 1179.6521 | RBD HEK 293-F | 100 s (23 °C) | 1.118225 | 0.017243 | 3.926499 | 0.033605 | |
| 503 | 512 | VGYQPYRVVV | | 8 | 1179.6521 | RBD HEK 293-F | 1,000 s (23 °C) | 1.971956 | 0.042551 | 3.923847 | 0.03174 | |
| 503 | 512 | VGYQPYRVVV | | 8 | 1179.6521 | RBD HEK 293-F | 10,000 s (23 °C) | 1.966426 | 0.003969 | 3.92422 | 0.021348 | |
| 503 | 512 | VGYQPYRVVV | | 8 | 1179.6521 | RBD HEK 293-F | 9 h (28 °C) | 1.996625 | 0.020876 | 3.90906 | 0.005364 | |
| 503 | 512 | VGYQPYRVVV | | 8 | 1179.6521 | RBD Abdala | 10 s (ice) | 0.290594 | 0.023049 | 3.92341 | 0.020763 | |
| 503 | 512 | VGYQPYRVVV | | 8 | 1179.6521 | RBD Abdala | 10 s (23 °C) | 0.468922 | 0.013987 | 3.949635 | 0.003824 | |
| 503 | 512 | VGYQPYRVVV | | 8 | 1179.6521 | RBD Abdala | 100 s (23 °C) | 1.115288 | 0.031982 | 3.958896 | 0.002052 | |
| 503 | 512 | VGYQPYRVVV | | 8 | 1179.6521 | RBD Abdala | 1,000 s (23 °C) | 1.949046 | 0.083704 | 3.95232 | 0.007071 | |
| 503 | 512 | VGYQPYRVVV | | 8 | 1179.6521 | RBD Abdala | 10,000 s (23 °C) | 2.013142 | 0.00266 | 3.928113 | 0.017944 | |
| 503 | 512 | VGYQPYRVVV | | 8 | 1179.6521 | RBD Abdala | 9 h (28 °C) | 2.020639 | 0.016722 | 3.938729 | 0.027917 | |
| 503 | 512 | VGYQPYRVVV | | 8 | 1179.6521 | MaxD | Max | 6.232109 | 0.034535 | 3.899752 | 0.003099 | 0.179985658 |
| 503 | 513 | VGYQPYRVVVL | | 9 | 1292.7361 | RBD HEK 293-F | 10 s (ice) | 0.326441 | 0.0097 | 4.73577 | 0.009851 | |
| 503 | 513 | VGYQPYRVVVL | | 9 | 1292.7361 | RBD HEK 293-F | 10 s (23 °C) | 0.458051 | 0.009913 | 4.724834 | 0.01628 | |
| 503 | 513 | VGYQPYRVVVL | | 9 | 1292.7361 | RBD HEK 293-F | 100 s (23 °C) | 1.170939 | 0.017053 | 4.735653 | 0.022178 | |
| 503 | 513 | VGYQPYRVVVL | | 9 | 1292.7361 | RBD HEK 293-F | 1,000 s (23 °C) | 2.003466 | 0.003612 | 4.735585 | 0.018551 | |
| 503 | 513 | VGYQPYRVVVL | | 9 | 1292.7361 | RBD HEK 293-F | 10,000 s (23 °C) | 2.021542 | 0.007809 | 4.734002 | 0.012633 | |
| 503 | 513 | VGYQPYRVVVL | | 9 | 1292.7361 | RBD HEK 293-F | 9 h (28 °C) | 2.027578 | 0.020312 | 4.731786 | 0.006278 | |
| 503 | 513 | VGYQPYRVVVL | | 9 | 1292.7361 | RBD Abdala | 10 s (ice) | 0.29534 | 0.036952 | 4.735023 | 0.007639 | |
| 503 | 513 | VGYQPYRVVVL | | 9 | 1292.7361 | RBD Abdala | 10 s (23 °C) | 0.436053 | 0.017056 | 4.749919 | 0.00431 | |
| 503 | 513 | VGYQPYRVVVL | | 9 | 1292.7361 | RBD Abdala | 100 s (23 °C) | 1.162942 | 0.029726 | 4.760791 | 0.001124 | |
| 503 | 513 | VGYQPYRVVVL | | 9 | 1292.7361 | RBD Abdala | 1,000 s (23 °C) | 1.958654 | 0.0551 | 4.750129 | 0.003768 | |

| | | | | | | | | | | | | |
|---|---|---|---|---|---|---|---|---|---|---|---|---|
| 503 | 513 | VGYQPYRVVVL | | 9 | 1292.7361 | RBD Abdala | 10,000 s (23 °C) | 2.079441 | 0.040997 | 4.744284 | 0.003864 | |
| 503 | 513 | VGYQPYRVVVL | | 9 | 1292.7361 | RBD Abdala | 9 h (28 °C) | 2.062954 | 0.035498 | 4.74384 | 0.014618 | |
| 503 | 513 | VGYQPYRVVVL | | 9 | 1292.7361 | MaxD | Max | 7.163681 | 0.042613 | 4.727028 | 0.004327 | 0.162142573 |
| 511 | 515 | VVLSF | | 4 | 564.3392 | RBD HEK 293-F | 10 s (ice) | 0.082025 | 0.059099 | 4.759084 | 0.012064 | |
| 511 | 515 | VVLSF | | 4 | 564.3392 | RBD HEK 293-F | 10 s (23 °C) | 0.123879 | 0.003708 | 4.749392 | 0.016532 | |
| 511 | 515 | VVLSF | | 4 | 564.3392 | RBD HEK 293-F | 100 s (23 °C) | 0.095212 | 0.003268 | 4.758574 | 0.020877 | |
| 511 | 515 | VVLSF | | 4 | 564.3392 | RBD HEK 293-F | 1,000 s (23 °C) | 0.172963 | 0.018634 | 4.754462 | 0.017699 | |
| 511 | 515 | VVLSF | | 4 | 564.3392 | RBD HEK 293-F | 10,000 s (23 °C) | 0.372936 | 0.004739 | 4.76199 | 0.011569 | |
| 511 | 515 | VVLSF | | 4 | 564.3392 | RBD HEK 293-F | 9 h (28 °C) | 0.673048 | 0.016557 | 4.755104 | 0.005661 | |
| 511 | 515 | VVLSF | | 4 | 564.3392 | RBD Abdala | 10 s (ice) | 0.075388 | 0.043436 | 4.760583 | 0.011193 | |
| 511 | 515 | VVLSF | | 4 | 564.3392 | RBD Abdala | 10 s (23 °C) | 0.070444 | 0.004569 | 4.772406 | 0.003942 | |
| 511 | 515 | VVLSF | | 4 | 564.3392 | RBD Abdala | 100 s (23 °C) | 0.071783 | 0.009488 | 4.784488 | 0.000686 | |
| 511 | 515 | VVLSF | | 4 | 564.3392 | RBD Abdala | 1,000 s (23 °C) | 0.16995 | 0.013441 | 4.781564 | 0.005969 | |
| 511 | 515 | VVLSF | | 4 | 564.3392 | RBD Abdala | 10,000 s (23 °C) | 0.385119 | 0.00039 | 4.76631 | 0.009309 | |
| 511 | 515 | VVLSF | | 4 | 564.3392 | RBD Abdala | 9 h (28 °C) | 0.722694 | 0.010394 | 4.770541 | 0.01577 | |
| 511 | 515 | VVLSF | | 4 | 564.3392 | MaxD | Max | 2.391891 | 0.030632 | 4.749972 | 0.004147 | 0.370555 |
| 514 | 533 | SFELLHAPATVCGPKKSTNL | | 17 | 2113.111 | RBD HEK 293-F | 10 s (ice) | 9.20291 | 0.066037 | 4.010375 | 0.022259 | |
| 514 | 533 | SFELLHAPATVCGPKKSTNL | | 17 | 2113.111 | RBD HEK 293-F | 10 s (23 °C) | 10.472912 | 0.016109 | 4.007203 | 0.027098 | |
| 514 | 533 | SFELLHAPATVCGPKKSTNL | | 17 | 2113.111 | RBD HEK 293-F | 100 s (23 °C) | 10.845785 | 0.061982 | 4.016683 | 0.035343 | |
| 514 | 533 | SFELLHAPATVCGPKKSTNL | | 17 | 2113.111 | RBD HEK 293-F | 1,000 s (23 °C) | 11.149696 | 0.034506 | 4.01478 | 0.034186 | |
| 514 | 533 | SFELLHAPATVCGPKKSTNL | | 17 | 2113.111 | RBD HEK 293-F | 10,000 s (23 °C) | 11.141946 | 0.004692 | 4.013181 | 0.022551 | |
| 514 | 533 | SFELLHAPATVCGPKKSTNL | | 17 | 2113.111 | RBD HEK 293-F | 9 h (28 °C) | 11.218568 | 0.059576 | 3.998094 | 0.004244 | |
| 514 | 533 | SFELLHAPATVCGPKKSTNL | | 17 | 2113.111 | MaxD | Max | 10.655337 | 0.03908 | 3.994025 | 0.003638 | 0.340226811 |
| 515 | 533 | FELLHAPATVCGPKKSTNL | | 16 | 2026.079 | RBD HEK 293-F | 10 s (ice) | 8.496143 | 0.076591 | 3.816741 | 0.022871 | |
| 515 | 533 | FELLHAPATVCGPKKSTNL | | 16 | 2026.079 | RBD HEK 293-F | 10 s (23 °C) | 9.626004 | 0.010395 | 3.814132 | 0.03025 | |
| 515 | 533 | FELLHAPATVCGPKKSTNL | | 16 | 2026.079 | RBD HEK 293-F | 100 s (23 °C) | 9.990907 | 0.11575 | 3.823932 | 0.036514 | |

| | | | | | | | | | | | | |
|---|---|---|---|---|---|---|---|---|---|---|---|---|
| 515 | 533 | FELLHAPATVCGPKKSTNL | | 16 | 2026.079 | RBD HEK 293-F | 1,000 s (23 °C) | 10.101537 | 1.64791E-05 | 3.821242 | 0.037185 | |
| 515 | 533 | FELLHAPATVCGPKKSTNL | | 16 | 2026.079 | RBD HEK 293-F | 10,000 s (23 °C) | 10.050822 | 0.023595 | 3.821611 | 0.022871 | |
| 515 | 533 | FELLHAPATVCGPKKSTNL | | 16 | 2026.079 | RBD HEK 293-F | 9 h (28 °C) | 10.085921 | 0.079516 | 3.805773 | 0.005412 | |
| 515 | 533 | FELLHAPATVCGPKKSTNL | | 16 | 2026.079 | MaxD | Max | 9.572831 | 0.028667 | 3.808566 | 3.28E-05 | 0.370208487 |
| 515 | 547 | FELLHAPATVCGPKKSTNLVKNKCVNFHHHHHH | | 30 | 3780.9227 | RBD HEK 293-F | 10 s (ice) | 9.753713 | 0.095082 | 2.600027 | 0.001974 | |
| 515 | 547 | FELLHAPATVCGPKKSTNLVKNKCVNFHHHHHH | | 30 | 3780.9227 | RBD HEK 293-F | 10 s (23 °C) | 10.510044 | 0.081045 | 2.588054 | 0.000362 | |
| 515 | 547 | FELLHAPATVCGPKKSTNLVKNKCVNFHHHHHH | | 30 | 3780.9227 | RBD HEK 293-F | 100 s (23 °C) | 10.973179 | 0.106355 | 2.59259 | 8.74E-05 | |
| 515 | 547 | FELLHAPATVCGPKKSTNLVKNKCVNFHHHHHH | | 30 | 3780.9227 | RBD HEK 293-F | 1,000 s (23 °C) | 11.211149 | 0.04703 | 2.598487 | 0.005157 | |
| 515 | 547 | FELLHAPATVCGPKKSTNLVKNKCVNFHHHHHH | | 30 | 3780.9227 | RBD HEK 293-F | 10,000 s (23 °C) | 11.479261 | 0.102017 | 2.59644 | 0.002232 | |
| 515 | 547 | FELLHAPATVCGPKKSTNLVKNKCVNFHHHHHH | | 30 | 3780.9227 | RBD HEK 293-F | 9 h (28 °C) | 11.262657 | 0.189994 | 2.596981 | 0.001413 | |
| 515 | 547 | FELLHAPATVCGPKKSTNLVKNKCVNFHHHHHH | | 30 | 3780.9227 | MaxD | Max | 10.555298 | 0.143293 | 2.581913 | 0.004563 | 0.629638667 |
| 516 | 533 | ELLHAPATVCGPKKSTNL | | 15 | 1879.0106 | RBD HEK 293-F | 10 s (ice) | 8.150978 | 0.058666 | 2.896507 | 0.001201 | |
| 516 | 533 | ELLHAPATVCGPKKSTNL | | 15 | 1879.0106 | RBD HEK 293-F | 10 s (23 °C) | 9.217471 | 0.013724 | 2.888566 | 0.002202 | |
| 516 | 533 | ELLHAPATVCGPKKSTNL | | 15 | 1879.0106 | RBD HEK 293-F | 100 s (23 °C) | 9.582265 | 0.017046 | 2.890705 | 0.001076 | |
| 516 | 533 | ELLHAPATVCGPKKSTNL | | 15 | 1879.0106 | RBD HEK 293-F | 1,000 s (23 °C) | 9.652309 | 0.005053 | 2.892328 | 0.001645 | |
| 516 | 533 | ELLHAPATVCGPKKSTNL | | 15 | 1879.0106 | RBD HEK 293-F | 10,000 s (23 °C) | 9.671722 | 0.004769 | 2.89536 | 0.000462 | |
| 516 | 533 | ELLHAPATVCGPKKSTNL | | 15 | 1879.0106 | RBD HEK 293-F | 9 h (28 °C) | 9.652852 | 0.015085 | 2.892108 | 0.001684 | |
| 516 | 533 | ELLHAPATVCGPKKSTNL | | 15 | 1879.0106 | MaxD | Max | 9.186337 | 0.069909 | 2.893674 | 0.001302 | 0.355344772 |
| 534 | 547 | VKNKCVNFHHHHHH | | 13 | 1773.8615 | RBD HEK 293-F | 10 s (ice) | 3.200221 | 0.080725 | 0.73739 | 0.000974 | |
| 534 | 547 | VKNKCVNFHHHHHH | | 13 | 1773.8615 | RBD HEK 293-F | 10 s (23 °C) | 3.051398 | 0.001267 | 0.738151 | 0.000752 | |
| 534 | 547 | VKNKCVNFHHHHHH | | 13 | 1773.8615 | RBD HEK 293-F | 100 s (23 °C) | 3.002264 | 0.004824 | 0.738434 | 0.000751 | |
| 534 | 547 | VKNKCVNFHHHHHH | | 13 | 1773.8615 | RBD HEK 293-F | 1,000 s (23 °C) | 3.150832 | 0.027095 | 0.737619 | 0.001227 | |
| 534 | 547 | VKNKCVNFHHHHHH | | 13 | 1773.8615 | RBD HEK 293-F | 10,000 s (23 °C) | 3.174372 | 0.002728 | 0.738487 | 0.000113 | |
| 534 | 547 | VKNKCVNFHHHHHH | | 13 | 1773.8615 | RBD HEK 293-F | 9 h (28 °C) | 3.297984 | 0.012358 | 0.739993 | 0.000478 | |
| 534 | 547 | VKNKCVNFHHHHHH | | 13 | 1773.8615 | MaxD | Max | 2.869624 | 0.013415 | 0.742837 | 0.003806 | 0.767641781 |
| 504 | 532 | FELLHAPATVCGPKKGGSGGSSSSSSSSS | | 26 | 2709.278449 | RBD Abdala | 10 s (ice) | 14.770892 | 0.259943 | 3.242494 | 0.01636 | |

| | | | | | | | | | | | | |
|---|---|---|---|---|---|---|---|---|---|---|---|---|
| 504 | 532 | FELLHAPATVCGPKKGGSGGSSSSSSSSS | | 26 | 2709.278449 | RBD Abdala | 10 s (23 °C) | 15.745322 | 0.017148 | 3.265329 | 0.000932 | |
| 504 | 532 | FELLHAPATVCGPKKGGSGGSSSSSSSSS | | 26 | 2709.278449 | RBD Abdala | 100 s (23 °C) | 15.225141 | 0.450025 | 3.270987 | 0.001936 | |
| 504 | 532 | FELLHAPATVCGPKKGGSGGSSSSSSSSS | | 26 | 2709.278449 | RBD Abdala | 1,000 s (23 °C) | 15.173838 | 0.62133 | 3.26744 | 0.002022 | |
| 504 | 532 | FELLHAPATVCGPKKGGSGGSSSSSSSSS | | 26 | 2709.278449 | RBD Abdala | 10,000 s (23 °C) | 15.808626 | 0.182312 | 3.247443 | 0.017278 | |
| 504 | 532 | FELLHAPATVCGPKKGGSGGSSSSSSSSS | | 26 | 2709.278449 | RBD Abdala | 9 h (28 °C) | 16.162814 | 0.191231 | 3.253594 | 0.019791 | |
| 504 | 532 | FELLHAPATVCGPKKGGSGGSSSSSSSSS | | 26 | 2709.278449 | MaxD | Max | 14.516802 | 0.234413 | 3.2319 | 0.00034 | 0.412275223 |
| 505 | 530 | ELLHAPATVCGPKKGGSGGSSSSSSS | | 23 | 2388.145978 | RBD Abdala | 10 s (ice) | 12.403979 | 0.250137 | 2.183404 | 0.005477 | |
| 505 | 530 | ELLHAPATVCGPKKGGSGGSSSSSSS | | 23 | 2388.145978 | RBD Abdala | 10 s (23 °C) | 13.077066 | 0.094029 | 2.176273 | 0.008024 | |
| 505 | 530 | ELLHAPATVCGPKKGGSGGSSSSSSS | | 23 | 2388.145978 | RBD Abdala | 100 s (23 °C) | 12.628304 | 0.259336 | 2.17502 | 0.00836 | |
| 505 | 530 | ELLHAPATVCGPKKGGSGGSSSSSSS | | 23 | 2388.145978 | RBD Abdala | 1,000 s (23 °C) | 12.653061 | 0.556343 | 2.169622 | 0.001445 | |
| 505 | 530 | ELLHAPATVCGPKKGGSGGSSSSSSS | | 23 | 2388.145978 | RBD Abdala | 10,000 s (23 °C) | 13.380487 | 0.277678 | 2.177465 | 0.009948 | |
| 505 | 530 | ELLHAPATVCGPKKGGSGGSSSSSSS | | 23 | 2388.145978 | RBD Abdala | 9 h (28 °C) | 13.536549 | 0.198457 | 2.176941 | 0.009222 | |
| 505 | 530 | ELLHAPATVCGPKKGGSGGSSSSSSS | | 23 | 2388.145978 | MaxD | Max | 12.236289 | 0.298639 | 2.160544 | 0.040294 | 0.439986773 |
| 505 | 532 | ELLHAPATVCGPKKGGSGGSSSSSSSSS | | 25 | 2562.210035 | RBD Abdala | 10 s (ice) | 15.119743 | 0.284033 | 2.160235 | 0.005919 | |
| 505 | 532 | ELLHAPATVCGPKKGGSGGSSSSSSSSS | | 25 | 2562.210035 | RBD Abdala | 10 s (23 °C) | 16.180664 | 0.10505 | 2.151677 | 0.004051 | |
| 505 | 532 | ELLHAPATVCGPKKGGSGGSSSSSSSSS | | 25 | 2562.210035 | RBD Abdala | 100 s (23 °C) | 15.426227 | 0.504251 | 2.145799 | 0.011571 | |
| 505 | 532 | ELLHAPATVCGPKKGGSGGSSSSSSSSS | | 25 | 2562.210035 | RBD Abdala | 1,000 s (23 °C) | 15.294505 | 0.779801 | 2.141646 | 0.002881 | |
| 505 | 532 | ELLHAPATVCGPKKGGSGGSSSSSSSSS | | 25 | 2562.210035 | RBD Abdala | 10,000 s (23 °C) | 16.181853 | 0.053968 | 2.152994 | 0.006024 | |
| 505 | 532 | ELLHAPATVCGPKKGGSGGSSSSSSSSS | | 25 | 2562.210035 | RBD Abdala | 9 h (28 °C) | 16.474156 | 0.067846 | 2.153348 | 0.009044 | |
| 505 | 532 | ELLHAPATVCGPKKGGSGGSSSSSSSSS | | 25 | 2562.210035 | MaxD | Max | 15.040717 | 0.380112 | 2.132179 | 0.041804 | 0.366706653 |
| 505 | 541 | ELLHAPATVCGPKKGGSGGSSSSSSSSSSIEHHHHHH | | 34 | 3713.722192 | RBD Abdala | 10 s (ice) | 14.842056 | 0.573113 | 1.144431 | 0.318588 | |
| 505 | 541 | ELLHAPATVCGPKKGGSGGSSSSSSSSSSIEHHHHHH | | 34 | 3713.722192 | RBD Abdala | 10 s (23 °C) | 15.774333 | 0.326624 | 0.946819 | 0.290131 | |
| 505 | 541 | ELLHAPATVCGPKKGGSGGSSSSSSSSSSIEHHHHHH | | 34 | 3713.722192 | RBD Abdala | 100 s (23 °C) | 15.024753 | 0.134515 | 1.054586 | 0.326489 | |
| 505 | 541 | ELLHAPATVCGPKKGGSGGSSSSSSSSSSIEHHHHHH | | 34 | 3713.722192 | RBD Abdala | 1,000 s (23 °C) | 15.604682 | 0.731239 | 0.780209 | 0.000284 | |
| 505 | 541 | ELLHAPATVCGPKKGGSGGSSSSSSSSSSIEHHHHHH | | 34 | 3713.722192 | RBD Abdala | 10,000 s (23 °C) | 15.737896 | 0.612686 | 1.124181 | 0.331504 | |
| 505 | 541 | ELLHAPATVCGPKKGGSGGSSSSSSSSSSIEHHHHHH | | 34 | 3713.722192 | RBD Abdala | 9 h (28 °C) | 16.450854 | 0.640104 | 1.009153 | 0.314148 | |

| | | | | | | | | | | | | |
|---|---|---|---|---|---|---|---|---|---|---|---|---|
| 505 | 541 | ELLHAPATVCGPKKGGSGGSSSSSSSSSSIEHHHHHH | | 34 | 3713.722192 | MaxD | Max | 13.865491 | 0 | 1.418041 | 0 | 0.570727833 |
| 505 | 541 | ELLHAPATVCGPKKGGSGGSSSSSSSSSSIEHHHHHH | Man6 (S29) | 34 | 3713.722192 | RBD Abdala | 10 s (ice) | 17.4304 | 0 | 1.144431 | 0.318588 | |
| 505 | 541 | ELLHAPATVCGPKKGGSGGSSSSSSSSSSIEHHHHHH | Man6 (S29) | 34 | 3713.722192 | RBD Abdala | 10 s (23 °C) | 18.1124 | 0 | 0.946819 | 0.290131 | |
| 505 | 541 | ELLHAPATVCGPKKGGSGGSSSSSSSSSSIEHHHHHH | Man6 (S29) | 34 | 3713.722192 | RBD Abdala | 100 s (23 °C) | 17.0427 | 0.0731 | 1.054586 | 0.326489 | |
| 505 | 541 | ELLHAPATVCGPKKGGSGGSSSSSSSSSSIEHHHHHH | Man6 (S29) | 34 | 3713.722192 | RBD Abdala | 1,000 s (23 °C) | 17.6918 | 0.7246 | 0.780209 | 0.000284 | |
| 505 | 541 | ELLHAPATVCGPKKGGSGGSSSSSSSSSSIEHHHHHH | Man6 (S29) | 34 | 3713.722192 | RBD Abdala | 10,000 s (23 °C) | 17.8654 | 0 | 1.124181 | 0.331504 | |
| 505 | 541 | ELLHAPATVCGPKKGGSGGSSSSSSSSSSIEHHHHHH | Man6 (S29) | 34 | 3713.722192 | RBD Abdala | 9 h (28 °C) | 17.618 | 0 | 1.009153 | 0.314148 | |
| 505 | 541 | ELLHAPATVCGPKKGGSGGSSSSSSSSSSIEHHHHHH | Man6 (S29) | 34 | 3713.722192 | MaxD | Max | 15.7211 | 0 | 1.418041 | 0 | 0.513278638 |
